# Spatiotemporal and ontogenetic variation, microbial selection, and predicted antifungal function in the skin-associated microbiome of a Rocky Mountain amphibian

**DOI:** 10.1101/2022.06.01.494434

**Authors:** Kenen B. Goodwin, Jaren D. Hutchinson, Zachariah Gompert

## Abstract

Host-associated microbiomes play important roles in host health and pathogen defense. In amphibians, the skin-associated microbiome serves as an innate immune defense with potential implications for disease management. Few studies have examined season-long temporal variation in the amphibian skin-associated microbiome, and the interactions between bacteria and fungi on amphibian skin remain poorly understood. We characterize season-long temporal variation in the skin-associated microbiome of the western tiger salamander (*Ambystoma mavortium*) for both bacteria and fungi between sites and across salamander life stages. 207 skin-associated microbiome samples were collected from salamanders at two Rocky Mountain lakes throughout the summer and fall of 2018, and 127 additional microbiome samples were collected from lake water and lake substrate. We used 16S and ITS next-generation sequencing data with Bayesian Dirichlet-multinomial regression to estimate the relative abundances of bacterial and fungal taxa, test for differential abundance, examine microbial selection, and derive alpha and beta diversity. The antifungal function of bacterial communities was predicted using stochastic character mapping and a database of antifungal bacterial isolates. We examined microbial absolute abundances using Bayesian negative binomial LASSO coupled with synthetic gene spike-ins. For both bacteria and fungi, we observed variation in community composition through time, between sites, and with salamander age and life stage. We found salamander skin to be selective for microbes, with many taxa disproportionately represented relative to the environment, and we observed selection for predicted antifungal bacteria. Ultimately, this ecological knowledge may assist in the conservation of amphibian species threatened by chytridiomycosis and other emerging diseases.

## INTRODUCTION

Host-associated microbiomes can interact with their hosts in many ways. Specialized metabolites produced by microbes can influence various aspects of host biology (Sharon *et al*. 2014), and host production of antimicrobial peptides can in turn influence microbial community structure (McFall-Ngai *et al*. 2013). Microbial communities are increasingly recognized as providing beneficial and necessary services for their hosts (Dethlefsen *et al*. 2007; Grice & Segre 2011) and maintaining and restoring healthy microbiomes can be important for host health (Tosh & McDonald 2011). Host-associated microbiomes can inhibit pathogens or parasites through competition, the activation of host immune responses, and the production of inhibitory secondary metabolites (Lee & Mazmanian 2010; Britton & Young 2014; Grunseich *et al*. 2019). An imbalance in the host-associated microbiome can permit transient opportunistic pathogens and resident microbes with pathogenic potential to harm the host (Lee & Mazmanian 2010).

As an innate immune defense, much attention has been given to the amphibian skin-associated microbiome for its potential in disease management (Walke & Belden 2016). For example, probiotic bioaugmentation and habitat management have the potential to influence the amphibian skin-associated microbiome and susceptibility to chytridiomycosis (Harris *et al*. 2009; Kueneman *et al*. 2016a; Grant *et al*. 2018), a devastating disease caused by the fungal pathogen *Batrachochytrium dendrobatidis* (hereafter *Bd*; Longcore *et al*. 1999; Skerratt *et al*. 2007). While amphibian skin-associated microbiomes are species-specific, vary with life history stage, and are distinct from environmental microbiomes (*i.e.*, soil, lake substrate, and lake water microbiomes), some variation in the microbiomes is attributable to location and abiotic water quality (McKenzie *et al*. 2011; Kueneman *et al*. 2013; Walke *et al*. 2014; Bletz *et al*. 2017a; Bletz *et al*. 2017b; Ellison *et al*. 2019). In particular, temperature influences the production of antifungal metabolites in amphibian skin-associated microbiomes (Daskin *et al*. 2014) and also impacts amphibian gut microbiome community structure (Kohl & Yahn 2016). Habitat disturbance, diet, captivity, pollutants, and pathogens can also influence the amphibian skin-associated microbiome (Antwis *et al*. 2014; Jani & Briggs 2014; Krynak 2015; Krynak *et al*. 2016; Kueneman *et al*. 2016a; Jani & Briggs 2018).

Although many studies have worked to characterize species-specific and spatial variation in the amphibian skin-associated microbiome, season-long temporal variation remains a major gap in our knowledge of the amphibian skin-associated microbiome with few applicable studies (Jiménez & Sommer 2016). Since both *Bd* infection prevalence and amphibian skin-associated microbiomes show seasonal and year-to-year variation (Savage *et al*. 2011; Longo *et al*. 2015; Familiar López *et al*. 2017; Douglas *et al*. 2021), season-long temporal variation in the amphibian skin-associated microbiome warrants investigation for its implications in disease management. Using a database of amphibian skin-associated microbiome antifungal bacterial isolates and their 16S rRNA gene sequences (Woodhams *et al*. 2015), Bletz *et al*. (2017a) found that despite significant changes in the skin-associated microbiome community structure of salamandrid newts, the relative abundances of bacteria with *Bd*-inhibitory potential did not change significantly during a 12-week sampling period nor across life history stages in two of the three species studied. The database by Woodhams *et al*. (2015) contains bacterial isolates demonstrated to have antifungal function in *in vitro* assays. However, the application of this database to predict amphibian skin-associated microbiome antifungal function is limited by our knowledge of how these bacterial isolates function on amphibian skin, and observing fungal responses to changes in bacterial abundances could assist in detecting bacterial-fungal relationships.

Despite the focus of many amphibian skin-associated microbiome studies on bacteria, few studies have examined how bacteria interact with fungi and other microeukaryotes on amphibian skin (Jiménez & Sommer 2016). Based on the maturation of defensive components in amphibian systems and the relative abundances of bacteria and fungi across life history stages, Kueneman (2015) proposed that larval stages of amphibians may depend on high relative abundances of antifungal bacteria before metamorphosis and the maturation of the host immune system. Further, Kueneman *et al*. (2016b) found many correlations between bacterial and fungal taxa on the skin of the western toad (*Anaxyrus boreas*). Hence, the interactions between bacteria and fungi on amphibian skin may have substantial implications for host health and disease management.

Broadly, our study aims to investigate temporal variation in the amphibian skin-associated microbiome using the western tiger salamander (*Ambystoma mavortium*; hereafter salamander) as a model amphibian. In the Rocky Mountains of North America, the western tiger salamander serves as an apex predator in many fishless high alpine lakes. When the snow melts at these lakes, adult salamanders travel to the lakes to breed, and some of these salamanders remain in the lakes throughout the early summer. During the summer months, eggs hatch and larval salamanders may follow several life history strategies, including metamorphosing during the same year as hatching, overwintering as larvae and metamorphosing the following year, and becoming sexually mature in the larval stage as neotenes (Sexton & Bizer 1978). Due to their local abundance and the presence of at least one life stage throughout the warm field season (June to September, hereafter season) at fishless high alpine lakes, the western tiger salamander is an ideal amphibian for consistently obtaining skin-associated microbiome samples through time.

In this study, we first examine season-long temporal variation of both bacteria and fungi in the salamander skin-associated microbiome between sites and across life history stages. Based on these data we identify differentially abundant microbes between salamander skin and the environment and compare the predictive ability of spatiotemporal and water quality covariates on microbial community composition. We then ask (i) whether variation in the salamander skin-associated microbiome influences predicted antifungal function, and (ii) whether predicted antifungal function is correlated with the observed relative abundance of *Bd*.

## METHODS

### Study Sites

Salamanders were sampled from the largest of the Gibson Lakes (Franklin County, ID; 447845 easting, 4654056 northing, NAD 83 UTM Zone 12; elevation: 2,579 m) and Ponds Lake (Summit County, UT; 503020 easting, 4503670 northing, NAD 83 UTM Zone 12; elevation: 3,058 m). These lakes were chosen for sampling due to their differences in geology, substrate, and water conditions. Both lakes are fishless, have no tributaries or outlets, and are located in different subranges of the Rocky Mountains. Gibson Lakes is a ∼2.5-ha shallow lake in a limestone basin of the Bear River Mountains. Patches of submerged vegetation cover much of the lake bottom, and the lake substrate is primarily composed of soft sticky mud. Ponds Lake is a ∼2.3-ha lake in a granitic basin of the Uinta Mountains. The lake substrate is a thick layer of loose vegetative material, and some parts of the shoreline have floating mats of vegetation. The water in Ponds Lake is stained red with dissolved organic carbon.

In 2018, access to Gibson Lakes was blocked due to snow at lower elevations until June 9^th^, when salamander eggs were observed attached to submerged vegetation. By the next week, when field sampling began, most of the previously observed eggs had hatched. Data from NRCS SNOTEL sites (see Supplemental Information) suggest that snow melted at both lakes at about the same time in 2018, possibly within days of each other, and snow typically melts at these lakes about a week apart. Based on these data, it is likely that salamanders laid eggs in 2018 at about the same time at both lakes.

### Sampling Design

To ensure that the sampled salamanders were distributed throughout the lakes, the lakes were sampled by strata. Gibson Lakes was assigned 4 strata and Ponds Lake was assigned 5 strata (Figure 1). Within a lake, all strata had roughly the same area, and their areas remained roughly the same as each other as water levels dropped throughout the season. Three age classes of salamanders could be distinguished based on length and weight measurements, age-0, age-1, and age-2+. During each visit to a lake (hereafter sampling event), we collected up to 20 salamanders from each age class with a maximum of five and four salamanders per stratum at Gibson Lakes and Ponds Lake, respectively. Each lake was sampled every other week during the 2018 season. Sampling began shortly after snowmelt and continued until the lakes became too cold to safely catch salamanders. Sampling began at Gibson Lakes on June 16^th^ and Ponds Lake on June 23^rd^. Gibson Lakes was too cold to sample on September 29^th^, marking the end of the field season.

**Figure 1.**
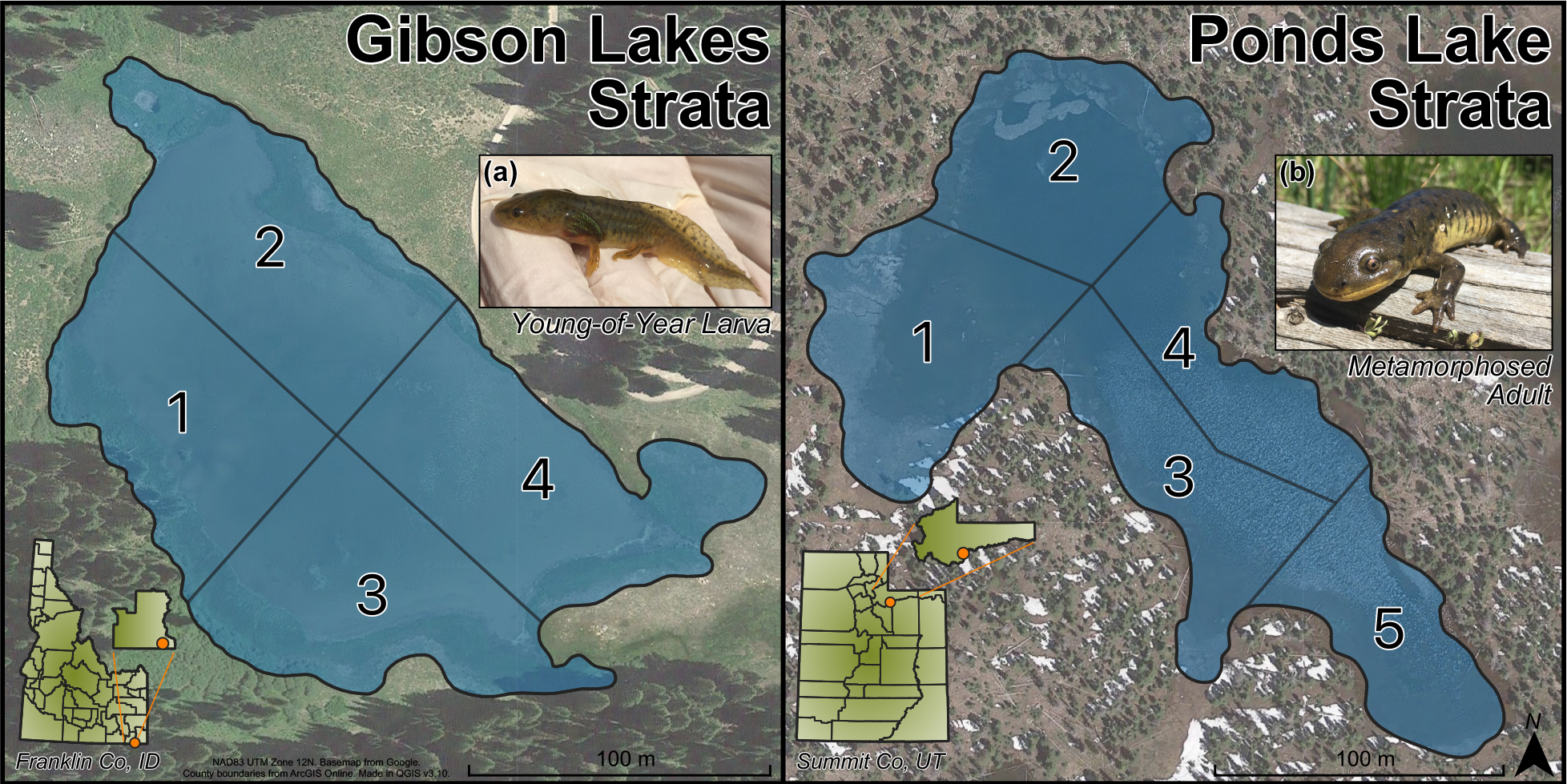
Strata for Gibson Lakes (left) and Ponds Lake (right). Images of (a) a young-of-year larval salamander and (b) a metamorphosed adult salamander.

Salamanders were considered larvae if they retained any of their larval gill structures, and salamanders were considered metamorphosed individuals once all traces of their gill structures were absorbed. For each age class, larval and metamorphosed individuals were encountered, which we refer to as life stages, and we refer to the six possible combinations of age class and life stage as stage classes. We expect most age-2+ individuals to be sexually mature adults, at which point gilled individuals are considered neotenes.

### Data Collection

Upon arriving at a lake, environmental microbiome samples and water quality data were collected. During the first visit to each lake, a location was selected just offshore in each stratum to collect these samples and data. These locations were chosen to have relatively homogeneous depths across strata and to minimize the distance which the sampling location would need to move with receding water levels. Water quality data was collected prior to collecting environmental microbiome samples to minimize disturbance to the water. Water temperature, pH, electrical conductivity, and dissolved oxygen (ppm and percent) were measured just below the water surface using handheld meters (Hannah Instruments HI98129 & HI9146). For sampling the lake water microbiome, 500 mL of lake water was collected from the water surface in a laboratory Nalgene bottle. Following collection of a lake water microbiome sample, a lake substrate microbiome sample was collected from the top ∼10 cm of pond substrate using a small PVC clam gun. The substrate column was deposited into a 15-mL conical tube, and excess water was decanted. The substrate was thoroughly stirred with a teasing needle, and ∼1.5 mL of substrate was deposited into a sterile 2-mL microcentrifuge tube. The microcentrifuge tubes containing substrate samples were placed in a cooler while in the field. New latex gloves were worn for each environmental microbiome sample, and the clam gun and teasing needle were rinsed with 95% ethanol between substrate samples and rinsed thoroughly with distilled water between sampling events. Nalgene bottles were rinsed thoroughly with distilled water and autoclaved for 20 minutes at 121 °C between holding lake water samples. During each sampling event, the depth of a predefined rock was measured to determine relative lake elevation, the water level of the lake relative to its height at the beginning of the season.

After collecting environmental microbiomes and water quality data for all strata, salamanders were captured for each stratum. Salamanders were collected by hand and dip net, and salamanders were stored in 5-gallon buckets filled with lake water. Each salamander was handled with new latex gloves, and snout-vent length (SVL) and weight measurements were taken to verify age classes (Figure S1). Sex was determined for age-2+ salamanders. The ventral surface of each salamander was rinsed with 50 mL of distilled water (Bletz *et al*. 2017a) to remove environmental material and transient microbes (Culp *et al*. 2007; Lauer *et al*. 2007), and the salamander’s ventral surface was swabbed with a sterile rayon-tipped swab (MW113 Medical Wire and Equipment, Corsham, UK). Swabbing was performed by stroking the swab across the ventral surface 10 times (1 time = an up and back stroke along the full length of the belly; Bletz *et al*. 2017a). Swabs used to sample salamander skin-associated microbiomes were stored in individual sterile 2-mL microcentrifuge tubes and placed in a cooler while in the field. After processing salamanders for a stratum was complete, the salamanders were released back into the stratum, and salamander collection began at the next stratum. While it is possible that salamanders sampled in one stratum may have been sampled again in another stratum during the same sampling event, few salamanders were observed to have swum far from their point of release.

For each sampling event, the lake was surveyed for salamanders for a minimum of 5 person-hours divided evenly among the lake’s strata. Salamanders were processed after the stratum minimum sampling time was reached or the maximum number of individuals from all available age classes had been collected, and the search for salamanders then proceeded to the next stratum. After field sampling and while still at the lake, wet and dry negative control swabs were taken. Wet control swabs were sprayed with 50 mL of distilled water, and nothing was done to the dry control swabs. Wet and dry control swabs were placed in individual sterile 2-mL microcentrifuge tubes and stored in a cooler while in the field.

Following field sampling and on the same day, lake water samples were prefiltered through a 5.0-μm prefilter membrane to remove debris followed by filtration with a 0.22-μm filter membrane to catch microbes (Millipore Sigma SVLP02500 and GSWP04700, respectively). Multiple 5.0-μm prefilter membranes were used for each water sample as necessary, whereas samples which experienced clogging on the 0.22-μm filter membrane were discarded. Following filtration, 0.22-μm filter membranes were folded and stored in 2-mL microcentrifuge tubes. For autoclavable filtration equipment, the equipment was rinsed thoroughly with distilled water between water samples followed by autoclaving for 20 minutes at 121 °C. Non-autoclavable filtration equipment was rinsed with 6% bleach solution followed by a thorough rinse with distilled water between water samples. Every four or five sampling events, five 500-mL distilled water samples were filtered as negative controls.

All samples were transferred to a −80 °C freezer for storage. Salamanders were collected, stored, handled, and released according to an approved Utah State University Institutional Animal Care and Use Committee protocol (#2798), a Utah Division of Wildlife Resources Certificate of Registration (#2COLL10232), and an Idaho Department of Fish and Game Wildlife Collection/Banding/Possession Permit (#180110).

### DNA Extraction, Sequencing, and Processing

DNA was extracted with the DNeasy PowerSoil Pro Kit (Qiagen, Inc.) following the manufacturer’s protocol, and 12 empty extractions were performed as blank negative controls. Substrate samples were centrifuged for 30 s at 10,000 x *g*, excess liquid was removed with a pipette, and a scoopula was used to collect 250 mg of substrate from each sample for DNA extraction. Water sample filter membranes were finely diced using scissors and forceps into reagent reservoirs before being transferred to DNA extraction tubes. Swab samples were transferred to DNA extraction tubes using a different pair of forceps than that used for water samples. Pre-DNA extraction sample preparation work was performed under a fume hood, and the scoopula, scissors, and forceps were rinsed with 95% ethanol, flamed, and rinsed thoroughly with distilled water between samples. Reagent troughs were rinsed thoroughly with distilled water and autoclaved for 20 minutes at 121 °C between water samples.

Following DNA extraction, two samples of ZymoBIOMICS Microbial Community DNA Standard (Zymo Research D6305) were added as mock community positive sequencing controls. 6 μL of a control oligo pool was added to 30 μL of full concentration DNA extract. The control oligo pool contained 0.01 pg/μL each of 16S and ITS well-specific cross contamination oligos (hereafter coligos; Hawkins *et al*. 2018) and 0.03 pg/μL each of 16S and ITS synthetic genes (hereafter synthgenes, which will be used in estimating absolute abundances of microbial taxa; Tourlousse *et al*. 2017). Sample DNA concentrations were measured via absorption and normalized to 10 ng/μL with an automated liquid handler. Combinatorial dual indexing was performed on the samples with two-stage polymerase chain reaction (PCR). First stage PCR amplified the 16S rRNA and ITS genetic barcoding regions, added unique dual index combinations to each sample, and added a portion of the Illumina Nextera adapter. For each sample, two first-stage PCR replicates were performed and subsequently pooled. Second stage PCR completed Illumina adapter addition. The 16S rRNA V4 region was amplified using the primers 515F (forward; Parada *et al*. 2016) and 806R (reverse; Caporaso 2011). The ITS1 region was amplified using the primers ITS1-F (forward; Gardes & Bruns 1993) and ITS2 (reverse; White *et al*. 1990). A modified AxyPrep MagBead PCR Clean-up protocol was used to purify the amplified DNA after each PCR reaction. Library preparation occurred at the University of Wyoming Genome Technologies Laboratory (Laramie, WY). See Supplemental Information for library preparation details.

Paired-end DNA sequencing of pooled amplicon product was performed on both Illumina MiSeq (v3 600-cycle kit, 2 x 300 base pair [bp] reads) and Illumina NextSeq (v2 300-cycle kit, 2 x 150 bp reads) platforms at the Utah State University Center for Integrated Biosystems (Logan, UT). We used MiSeq sequences as study-specific 16S and ITS reference libraries to enhance the taxonomic precision of our NextSeq data, for which there was greater sequencing depth. MiSeq produced 19 million paired-end reads and NextSeq produced 187 million.

MiSeq reads were partitioned into 16S and ITS datasets based on their primer regions using Perl (version 5.18.1), and sample barcodes were removed. Since variable length sample barcodes were used, MiSeq reads were trimmed to 290 bp using cutadapt (version 2.10; Martin 2011) to ensure that non-overlapping sequences did not appear different simply due to read length. Using cutadapt, read pairs that contained Ns were removed, and forward primers and reverse complements of reverse primers were trimmed (with a maximum error rate of 0.15, a minimum trimmed length of 1 bp, and discarding untrimmed read pairs), with trimming the reverse primer’s reverse complement being required for 16S read pairs.

The DADA2 bioinformatics pipeline (version 3.10; Callahan *et al*. 2016) was used in the R statistical software program (version 4.0.2; R Core Team 2020) for quality filtering, phiX removal, denoising, merging pairs, chimera removal, and taxonomic assignment of MiSeq reads (see Supplemental Information for details). For ITS sequences, non-overlapping read pairs were retained in the pipeline as concatenated sequences with 10-N spacers. A naïve Bayesian classifier (Wang *et al*. 2007) was used to classify unique sequences in the MiSeq 16S and ITS datasets using Silva (version 138; Quast *et al*. 2012) and UNITE (general dynamic FASTA release for fungi; version 8.2; Nilsson *et al*. 2019) reference libraries, respectively. To create study-specific 16S and ITS reference libraries, NextSeq-length forward and reverse reads were created from the classified MiSeq 16S and ITS sequences, and consensus taxonomies and MiSeq-length sequences (for antifungal prediction) were generated for duplicate reference read pairs (see Supplemental Information for details). Integers were appended to reference taxa names to differentiate each amplicon sequence variant (ASV) associated with a taxon.

NextSeq reads were assigned to barcode regions and samples using Perl while allowing 1 bp mismatches in sample barcodes. phiX reads were discarded, and sample barcodes were removed. The following steps were performed sequentially on the NextSeq reads using cutadapt: reads were trimmed to 140 bp to make all reads the same length, read pairs with Ns were removed, forward primers and reverse complements of reverse primers were trimmed (with the same settings as the MiSeq data but without requiring trimming of the reverse primers’ reverse complements).

Using exact matching in R, 21.4 million of 54.3 million NextSeq 16S reads were identified to 15,792 reference sequences, and 60.4 million of 113.9 million NextSeq ITS reads were identified to 3,488 reference sequences. All samples were checked for between-well cross contamination through use of the coligos. Three salamander samples and one blank control sample were removed from the 16S dataset due to high amounts of between-well contamination (having a ratio of any contaminant coligo to non-contaminant coligo greater than 0.1 after summing coligo read counts across PCR replicates). Two salamander samples were removed from the ITS dataset due to lack of detection of any non-synthgene and non-coligo sequences in both PCR replicates. Coligos were removed from the datasets for all subsequent analyses. In the mock community samples, we observed strong amplification bias in the ITS data (Figure S2; see Supplemental Information), and one fungal taxon was split into three substantial ASVs. In an effort to mitigate the potential impact of fungal taxa being split into multiple ASVs, we merged fungal ASVs which were assigned the same taxonomy into the same taxa. We chose to forego rarefaction of our samples as it increases uncertainty in relative abundances (McMurdie & Holmes 2014).

We performed principal component analyses (PCAs) on the proportional abundances of taxa across PCR replicate and sample type (Figures S3-S5; see Supplemental Information). Taxa proportional abundances within samples were similar across PCR replicates, so read counts were summed across PCR replicates for each sample. There were 10 salamander samples which grouped closely with wet swab and dry swab negative controls in the 16S PCAs on sample type, so these samples were removed from the 16S data for all subsequent analyses. Following Harrison *et al*. (2021), we calculated point estimates for microbial absolute abundance in each sample, and we observed higher microbial absolute abundance in field samples compared to their associated negative controls (Figure S6).

We examined microbial diversity between field sample types (*i.e.*, salamander, water, and substrate) through visualization. For each barcode region, each microbiome sample had its taxa sorted by descending read count, and the read count for each taxon was iteratively added to the cumulative read count of the preceding taxa. The median and interquartile range of the cumulative proportion of reads by cumulative number of sorted taxa was plotted for each field sample type and barcode region (Figure S7). Additionally, we checked the proportion of reads in each salamander sample which was classified to each taxonomic level by barcode region, life stage, and site using boxplots (Figure S8).

### Antifungal Prediction

A database of amphibian skin-associated microbiome antifungal bacterial isolates (Woodhams *et al*. 2015) was used to predict which bacteria observed in our datasets exhibit *Bd*-inhibitory properties (see Supplemental Information for details). We trimmed sequences in the Woodhams database to the 16S rRNA V4 region using our 16S amplification primers with R, and we aligned the MiSeq 16S sequences of taxa detected in our NextSeq 16S field samples with the Woodhams sequences using Clustal Omega (version 1.2.4; Sievers *et al*. 2011). We used FastTree 2 (version 2.1.11; Price *et al*. 2010) to create a phylogenetic tree, and we used stochastic character mapping with the make.simmap function in the phytools package (version 0.7.70; Revell 2012) to predict the *Bd*-inhibition statuses of our observed taxa. We visualized the phylogenetic tree with posterior probabilities of our taxa being *Bd*-inhibitory using the Interactive Tree of Life (Figure S9; version 6.5.4; Letunic & Bork 2021).

The vast majority of our taxa had low confidence in their antifungal statuses (99.5% of posterior probabilities were between 47.9% and 52.5%), whereas most posterior probabilities which were < 40% or > 60% were also ≤ 10% or ≥ 90% (33 of 39). Therefore, we considered our bacterial taxa to be *Bd*-inhibitory if their posterior probabilities of *Bd*-inhibition were ≥ 90%, and we considered our bacterial taxa to be non-*Bd*-inhibitory if their posterior probabilities of *Bd*-inhibition were ≤ 10%. Otherwise, we considered our bacterial taxa to have an uncertain *Bd*-inhibition status.

### Microbial Composition Modeling

For both bacterial and fungal communities, we fit Bayesian Dirichlet-multinomial regression models to the salamander, water, and substrate microbiome data to identify differentially abundant microbes and to evaluate differences in overall community composition. Our Dirichlet-multinomial regression model was adapted from the Dirichlet regression model of Sennhenn-Reulen (2018), and we used backwards variable selection by WAIC to optimize predictive accuracy. In our model, sample read counts are distributed according to the Dirichlet-multinomial distribution. Each taxon receives a linear predictor combination, and the softmax function normalizes linear predictor combinations for all taxa into expected proportions. The last taxon serves as a reference category, and its intercept and regression coefficients are set to zero to allow for model identifiability. A precision parameter controls the degree of overdispersion relative to the multinomial distribution. See Supplemental Information for model details.

To keep model run-times practical, we opted to select the 100 most proportionally abundant taxa plus an “other” category for inclusion in the composition models. To select these taxa, we averaged point estimates of taxa proportional abundances in each metabarcoding region of the salamander samples by combinations of site and life stage. We then averaged across these averages, and we took the 100 taxa with the highest averaged proportional abundances for each barcode region for use in modeling. Other taxa which were not included in the top 100 for each barcode region had their read counts merged into an “other” category. Datasets used in the modeling of water and substrate microbial communities included the same taxa as used for the salamander modeling, plus their own “other” categories. Since not all top microbial taxa in the salamander samples were detected in the water and substrate samples, the water and substrate datasets used in modeling had fewer than 101 taxa.

As water quality was highly correlated with space and time (Figure S10), we fit models with two different sets of predictors. One predictor set included spatiotemporal covariates, while the other predictor set substituted spatiotemporal covariates with water quality. The spatiotemporal predictor set included four-way interactions between age, life stage, site, and a second-degree polynomial for week, all lower-level interactions, and the individual predictors. Stratum was also included as a predictor and treated as a hierarchical effect. The water quality predictor set included a five-way interaction between age, life stage, temperature (°C), pH, and dissolved oxygen (ppm), all lower-level interactions, and the individual predictors. Models for water and substrate lacked age and life stage predictors. Site and life stage were treated as categorical predictors, and age and week were treated as continuous predictors. Age took whole integers from zero (age-0) to two (age-2+), and week represented the number of weeks since June 9^th^, 2018.

Models were fit in Stan (version 2.21.0; Carpenter *et al*. 2017) using the rstan R interface (version 2.21.2; Stan Development Team 2020) with four Hamiltonian Monte Carlo (HMC) chains, 500 warmup iterations, 500 sampling iterations, and no thinning. *R̂* convergence diagnostics and trace plots of the posteriors were used to assess model convergence. Since the effect of regression coefficients on proportional abundances is difficult to interpret (see Supplemental Information for a discussion), we opted for a graphical interpretation of the best-fit models (*i.e.*, the models selected by backwards variable selection). We generated posterior predictions of proportional abundances for each combination of non-stratum predictors observed in the datasets, where predictions were for the average stratum (see Supplemental Information for prediction details).

Measures of alpha and beta diversity were derived from the posterior distributions of taxa proportional abundances. For salamander microbiomes, we derived Hill’s diversity index with α = 2, which is robust to the many rare taxa expected in microbial communities (Haegeman *et al*. 2013). For each salamander stage class, we also derived posterior distributions for Bray-Curtis dissimilarity (calculated with the vegdist function in the vegan package; version 2.5.6; Oksanen *et al*. 2019) between all dates which the stage class was observed. The posterior distributions for taxa proportional abundances, Hill’s diversity index, and Bray-Curtis dissimilarity were summarized with 95% credible intervals, 50% credible intervals, and their median values.

To estimate the antifungal function of microbial communities, we summed the posterior proportional abundance predictions of taxa belonging to each *Bd*-inhibition category, with “other” bacterial taxa belonging to none of these categories. We summarized the proportional abundances of each *Bd*-inhibition category with 95% credible intervals, 50% credible intervals, and their median values. Of the top 100 bacterial taxa in the salamander composition models, 11 were *Bd*-inhibitory, four were non-*Bd*-inhibitory, and 85 had an uncertain *Bd*-inhibition status (Table S1).

To examine which taxa were disproportionately abundant on salamander skin relative to the environment, we considered the proportional abundance of microbes that salamanders experience in their environments to be a mixture between water and substrate proportional abundances, with the mixing ratio being a product of salamander behavior. Although we do not know this ratio, we expect that the result of this mixture is between the lower of the 0.025 quantiles and the upper of the 0.975 quantiles of the proportional abundance posterior predictions from water and substrate, and we consider this range to represent the proportional abundance of a taxon in the environment. In determining this range, if a taxon was not detected in the water or substrate samples, and therefore was not included in the compositional abundance modeling for that sample type, it was considered to have 0.025 and 0.975 quantiles of proportional abundance predictions for that sample type of zero.

### Microbial Absolute Abundance Modeling

Following DNA extraction and prior to PCR, fixed amounts of 16S and ITS synthgenes were added to a constant volume of each sample’s DNA extract. The synthgene read counts give us a benchmark to compare taxon read counts with, serving as the basis of our absolute abundance modeling. To model the density of the top 100 microbial taxa on salamander skin, we used a Bayesian negative binomial LASSO model for each taxon. Briefly, we modeled expected taxon read count as a product of taxon density (arbitrary units), synthgene read count, and an estimated value proportional to swabbed area. Taxon density is related to a linear predictor combination via a log link, and a size parameter controls the degree of overdispersion relative to the Poisson distribution. Our model borrows from the Bayesian LASSO (Park & Casella 2008) and Bayesian group LASSO (Xu & Ghosh 2015). We treated stratum as a grouped predictor and included four-way interactions between a second-degree polynomial for age, life stage, site, and a fourth-degree polynomial for week, all lower-level interactions, and the individual predictors. We generated posterior predictions of microbial densities for each combination of non-stratum predictors observed in the salamander datasets, where predictions were for the average stratum. See Supplemental Information for details.

## RESULTS

### Field Sampling

Total sample counts are included in Table 1, and a breakdown of salamander skin-associated microbiome samples are displayed in Figure 2. The residency of different salamander age classes varied through time, and age-0 salamanders were too small to sample during the early season. Based on observed sizes of males and females, most age-2+ salamanders are thought to have been adults (see Supplemental Information). Temperature and dissolved oxygen followed similar temporal trends at both lakes, while pH and conductivity were higher at Gibson Lakes (Figure S10).

**Figure 2.**
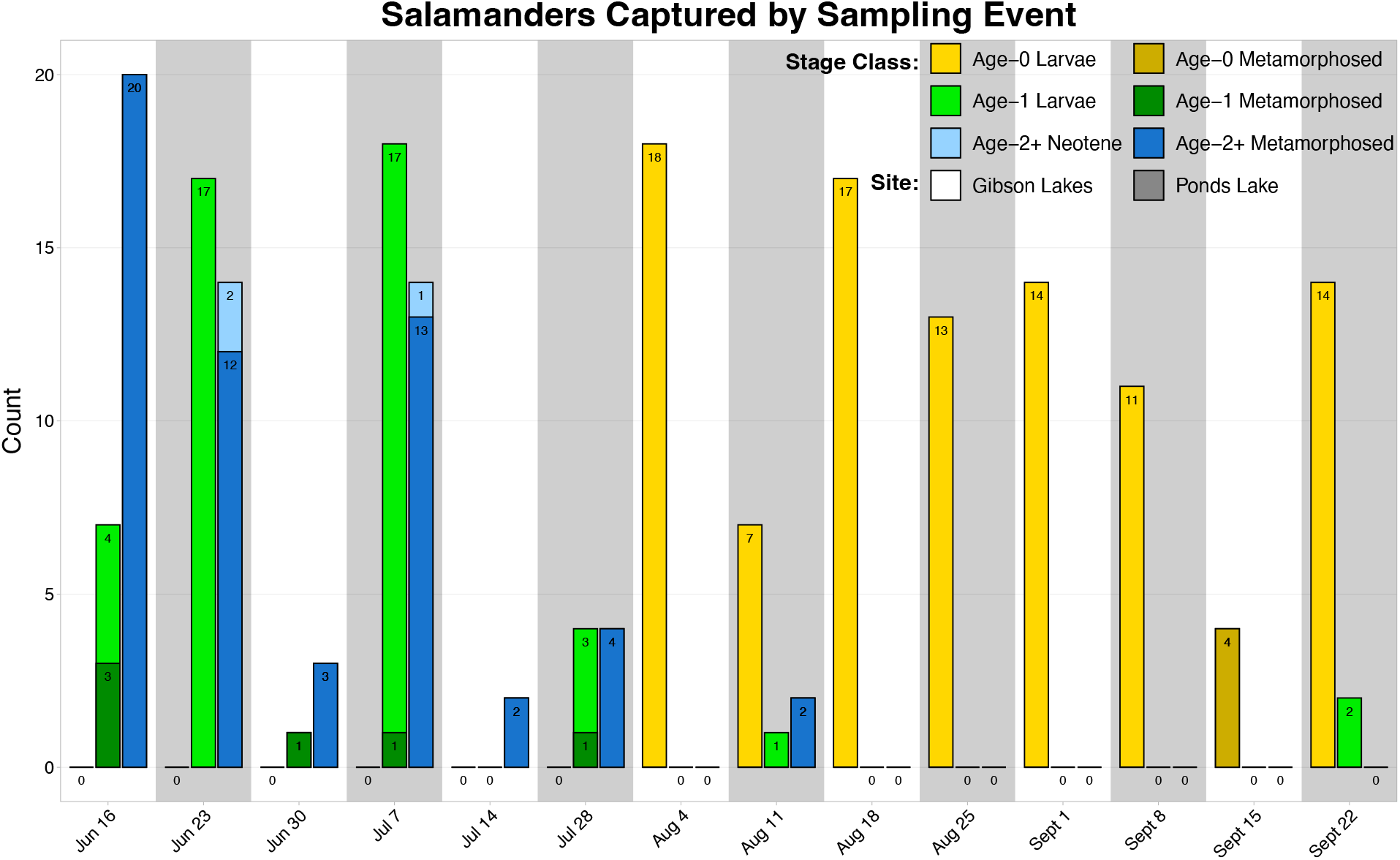
Counts of salamander skin-associated microbiome samples through time.

**Table 1.** Microbiome sample counts.

| Format | Type | Count |
| --- | --- | --- |
| Swab | Salamander | 207 |
|  | Wet Negative Control | 11 |
|  | Dry Negative Control | 13 |
| Water | Sample | 60 |
|  | Negative Control | 20 |
| Substrate | Sample | 67 |
| Blank | Negative Control | 12 |
| Mock Community | Positive Control | 2 |
| Total |  | 392 |

### Microbial Diversity Across Sample Types

We detected 15,690 bacterial taxa (6, 529 for salamander, 8,873 for water, and 14,591 for substrate) and 469 fungal taxa (289 for salamander, 224 for water, and 413 for substrate) in our field samples (using the NextSeq data). Graphically, substrate samples were the most diverse for both bacterial and fungal communities (Figure S7). Salamander and water samples were the least diverse graphically for bacterial and fungal communities, respectively.

### Antifungal Prediction

Only 33 of the 15,690 taxa detected in our NextSeq 16S field samples were classified as *Bd*-inhibitory or non-*Bd*-inhibitory (*i.e.*, posterior probabilities ≥ 90% or ≤ 10%), and of these, only 15 taxa were individually included (*i.e.*, were members of the top 100) in the composition modeling (Table S1). Of the 872 Woodhams sequences which were used in the alignment, there were 361 unique sequences, and 79 of these unique sequences occurred across multiple bacterial isolates. We note that 41 of these 79 sequences (51.9%) had inconsistent antifungal statuses (*i.e.*, statuses varied across isolates associated with the same sequence). We also note that the aligned Woodhams sequences provided limited phylogenetic coverage of the bacterial taxa detected in our field samples (Figure S9), with only 5,330 (34.0%) of our bacterial taxa belonging to phyla included in the Woodhams database.

### Microbial Composition

Composition models with spatiotemporal predictors fit better than models with water quality predictors for salamander samples (Table 2), suggesting that our spatiotemporal predictors were better able to predict salamander microbial composition. All but one of the best-fitting spatiotemporal composition models included stratum as a predictor, suggesting compositional variation in microbial communities within the lakes, with the salamander ITS model being the exception. Except for stratum for salamander fungal communities, all best-fitting spatiotemporal composition models included all individual predictors or their interactions, suggesting that all of our measured variables contributed to our ability to predict microbial communities.

**Table 2.**
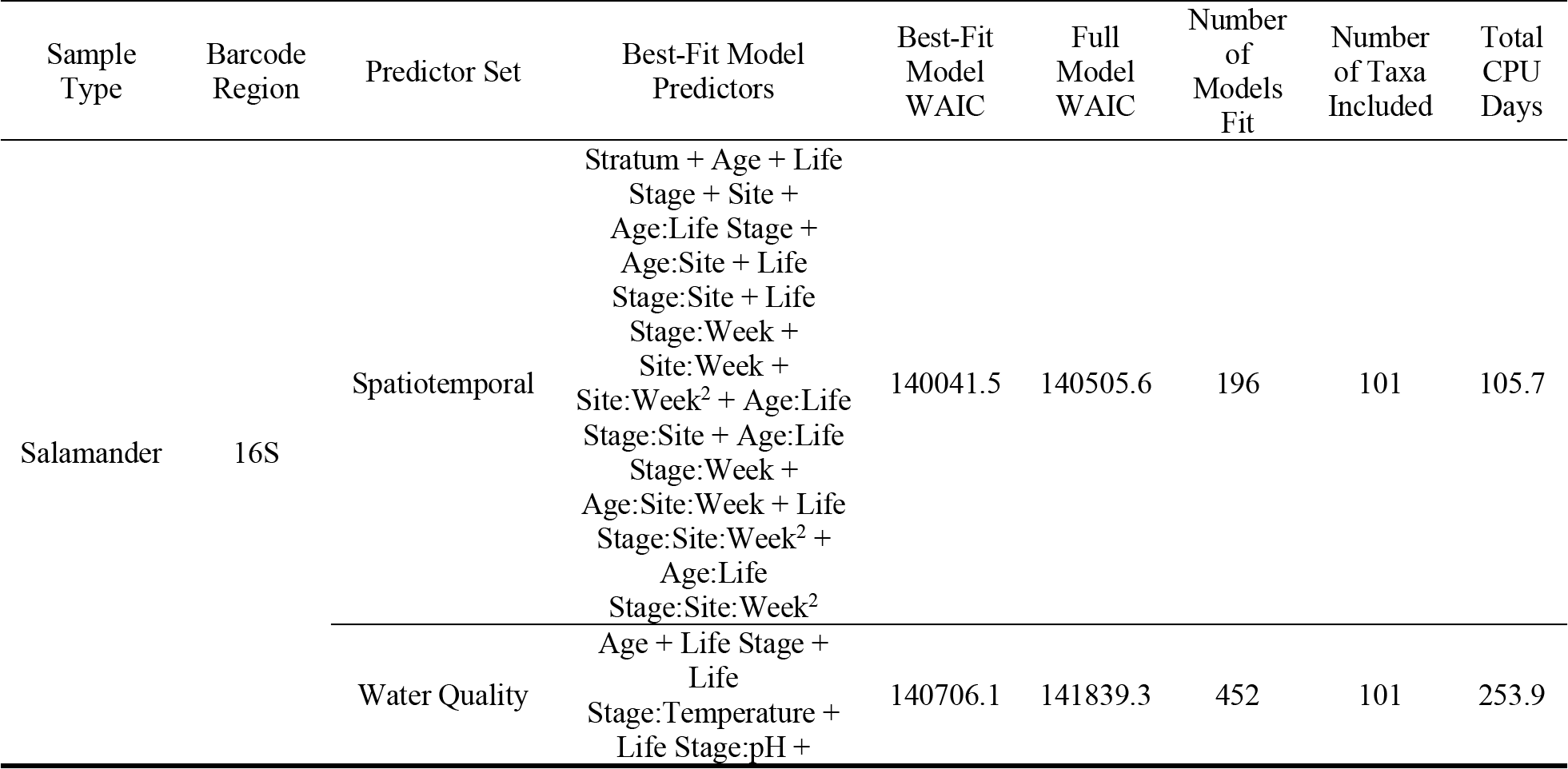
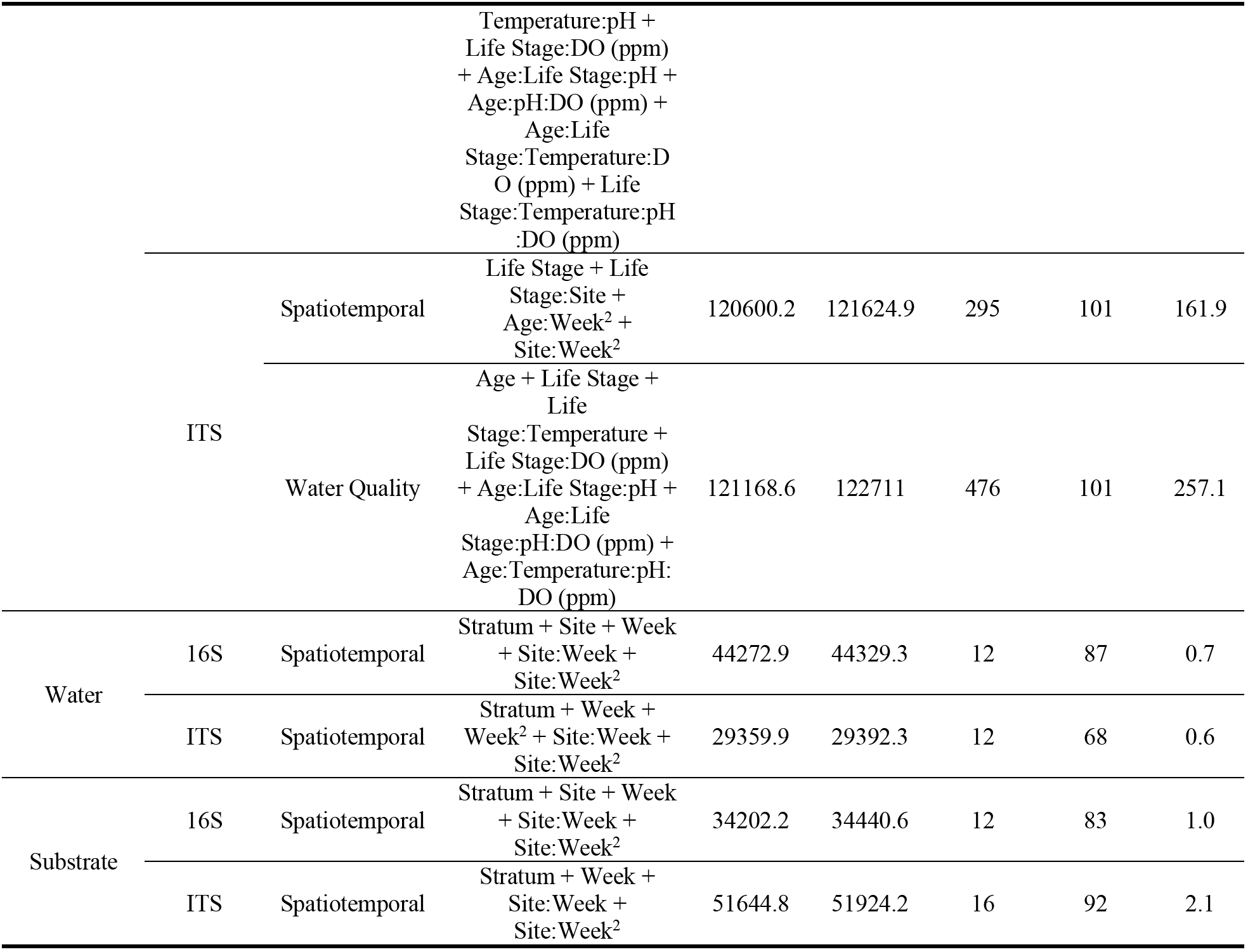
Predictors included in the best-fitting microbial composition models. Best-fit models are those selected by backwards variable selection by WAIC, and the full model is the initial model fit during backwards variable selection which includes all predictors. DO is dissolved oxygen.

We observed temporal, spatial, and ontogenetic variation in the proportional abundances of the top 100 bacterial taxa (Figures 3 & S11-S15), with the degree of variation depending on the taxon. Proportional abundance trends were often taxon-specific, although patterns were observed across some taxa. Examples of temporal variation include an increase in the proportional abundance of Comamonadaceae 2 (*i.e.*, the second ASV classified as Comamonadaceae) through time in Gibson Lakes age-0 larvae and a decrease in the proportional abundance of Candidatus Methylopumilus 1 through time in Ponds Lake age-1 larvae. Ontogenetic variation is apparent among many of the top 100 bacterial taxa. For example, the proportional abundances of Comamonadaceae 3 and 6 in Ponds Lake were consistently higher for age-2+ metamorphosed salamanders than other stage classes. In Gibson Lakes age-0 individuals, we observed higher proportional abundances of certain bacterial taxa on larvae followed by a sharp decline post-metamorphosis (*e.g.*, Comamonadaceae 2, Limnohabitans 1, Methylotenera 7, Methylotenera 2, and *Sphingorhabdus rigui* 1).

**Figure 3.**
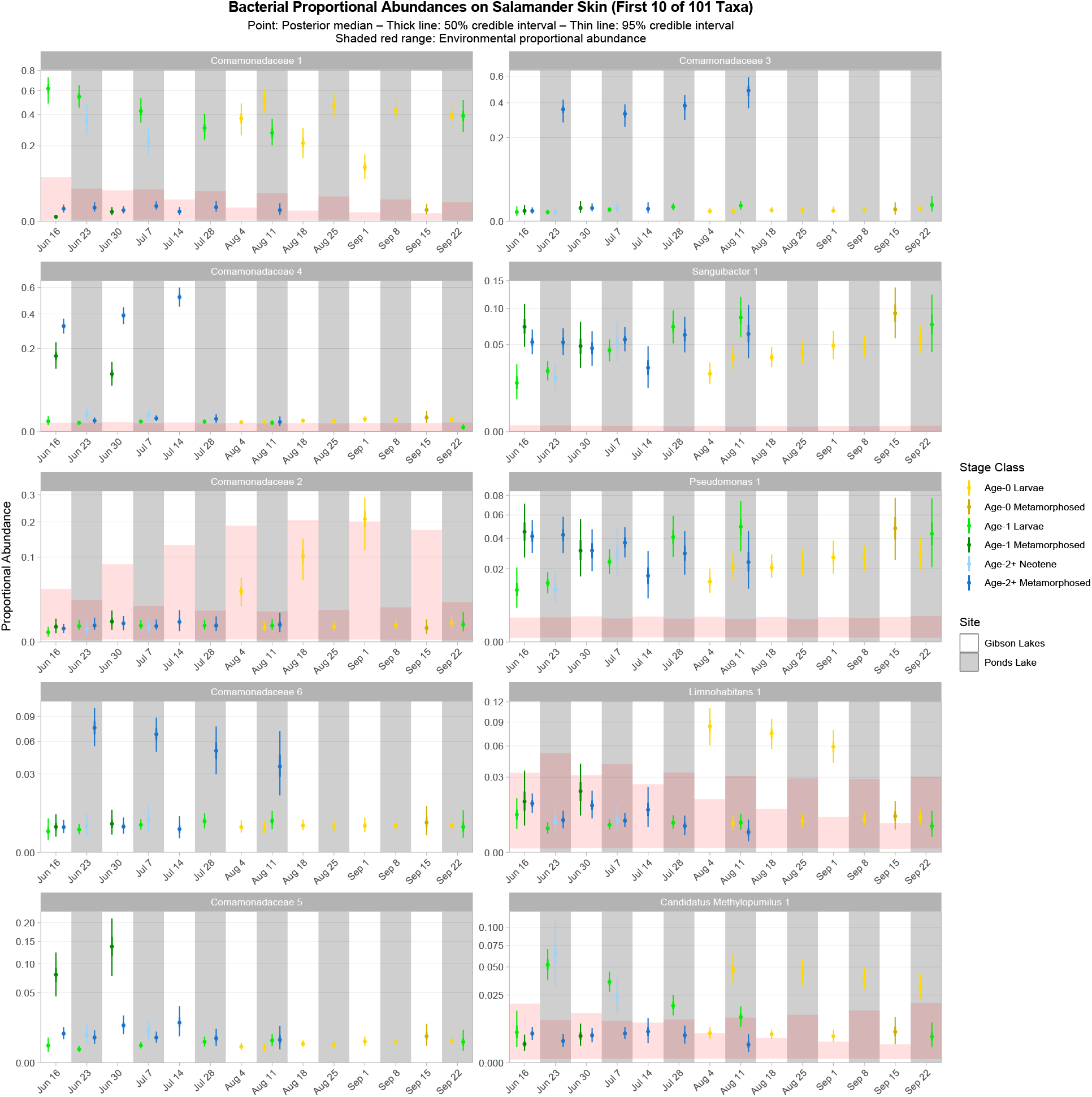
Proportional abundance predictions for the first ten bacterial taxa from the top 100. Points, thick lines, and thin lines represent posterior medians, 50% credible intervals, and 95% credible intervals, respectively. Shaded red ranges represent environmental proportional abundances. Taxa without shaded red ranges were not detected in either water or substrate. Note the square root scale on the y-axis. Taxa are ordered (left to right, top to bottom) by descending average proportion of reads in salamander samples in which each combination of site and life stage receives equal weight. Proportional abundance prediction plots for the remaining top 100 bacterial taxa can be found in the Supplemental Information.

90 of the top 100 bacterial taxa detected in salamander samples were also detected in environmental samples. We detected 86 and 82 of the top 100 bacterial taxa in water and substrate samples, respectively. 32 of the top 100 bacterial taxa were disproportionately more abundant on salamander skin relative to the environment for the majority of combinations of stage class and sampling event (Figures 3 & S11-S15), including Sanguibacter 1 and Gracilibacteria 3. Taxa detected exclusively on salamander skin include *Pseudochrobactrum kiredjianiae* 1 and Roseomonas 2. None of the top 100 bacterial taxa had lower proportional abundances on salamander skin than in the environment for the majority of combinations of stage class and sampling event.

Salamander bacterial diversity was highest among early-season age-1 metamorphosed individuals at Gibson Lakes, age-2+ neotenes at Ponds Lake in July, and age-0 individuals after mid-August at Gibson Lakes (Figure 4). Bacterial diversity for age-1 larvae increased throughout the early season at Ponds Lake when most of this stage class was observed. Bacterial diversity in age-2+ metamorphosed individuals tended to decrease through time at both lakes. After mid-August, bacterial diversity for all observed stage classes was higher at Gibson Lakes than Ponds Lake. For a given stage class, salamander bacterial communities were more dissimilar between sites than through time, with this pattern being more pronounced for age-2+ metamorphosed than age-0 larval individuals (Figure S16).

**Figure 4.**
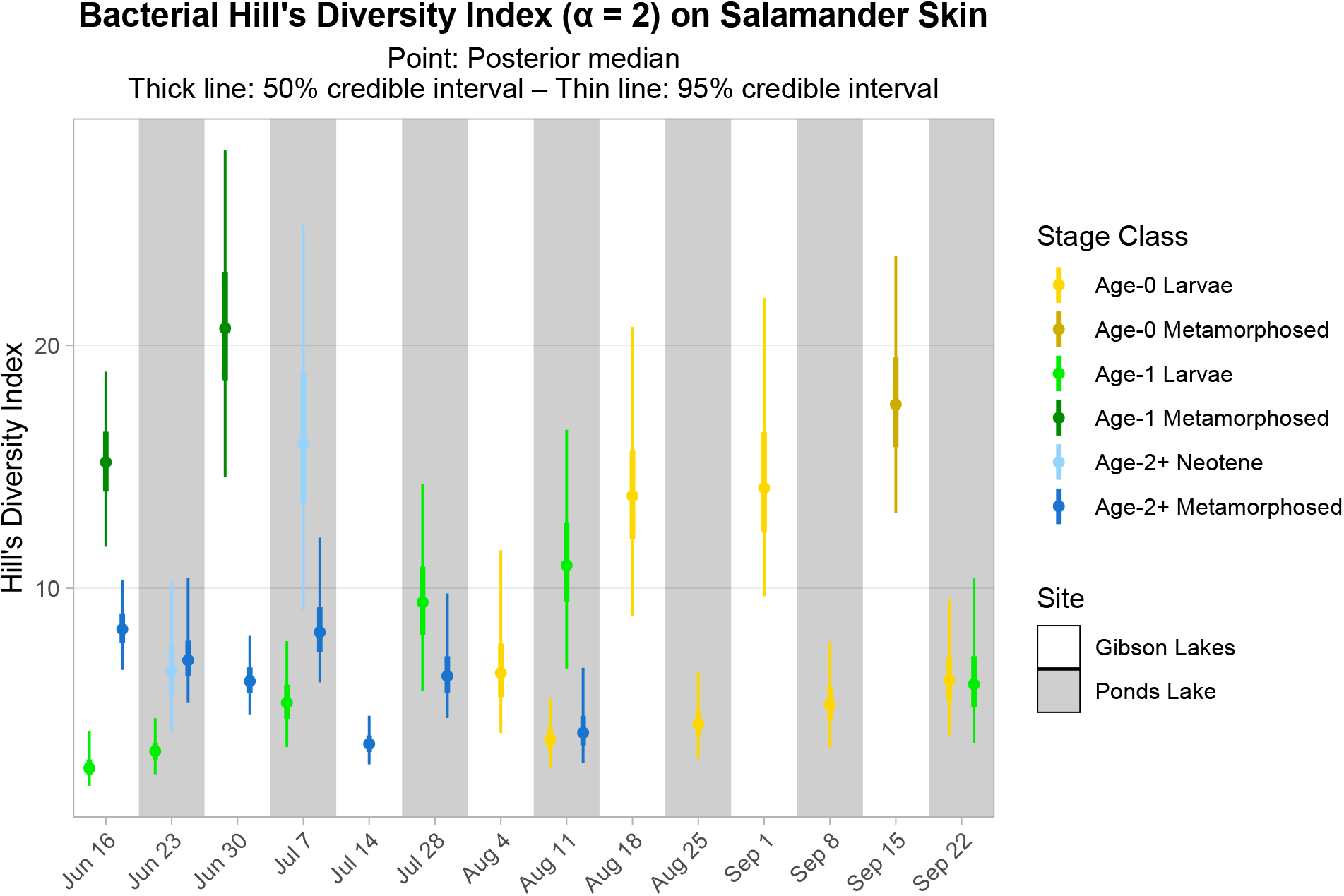
Hill’s diversity index (α = 2) for bacterial communities on salamander skin. Points, thick lines, and thin lines represent posterior medians, 50% credible intervals, and 95% credible intervals, respectively.

As for bacteria, we observed spatiotemporal and ontogenetic variation in the proportional abundances of the top 100 fungal taxa (Figures 5 & S17-S21). *Naganishia diffluens* experienced an increase in proportional abundance through time for age-0 and age-2+ salamanders at both lakes, and the proportional abundance of *Vishniacozyma victoriae* increased through time for Gibson Lakes age-0 larvae. Notably, the proportional abundance of *Bd* in Ponds Lake metamorphosed individuals was very high (between 35% and 55%) compared to Ponds Lake larvae and either life stage at Gibson Lakes (all < 2.5%). In contrast, metamorphosed individuals at Ponds Lake had lower proportional abundances of *Cystobasidium slooffiae* compared to Ponds Lake larvae and either life stage at Gibson Lakes, the opposite of the pattern observed for *Bd*.

**Figure 5.**
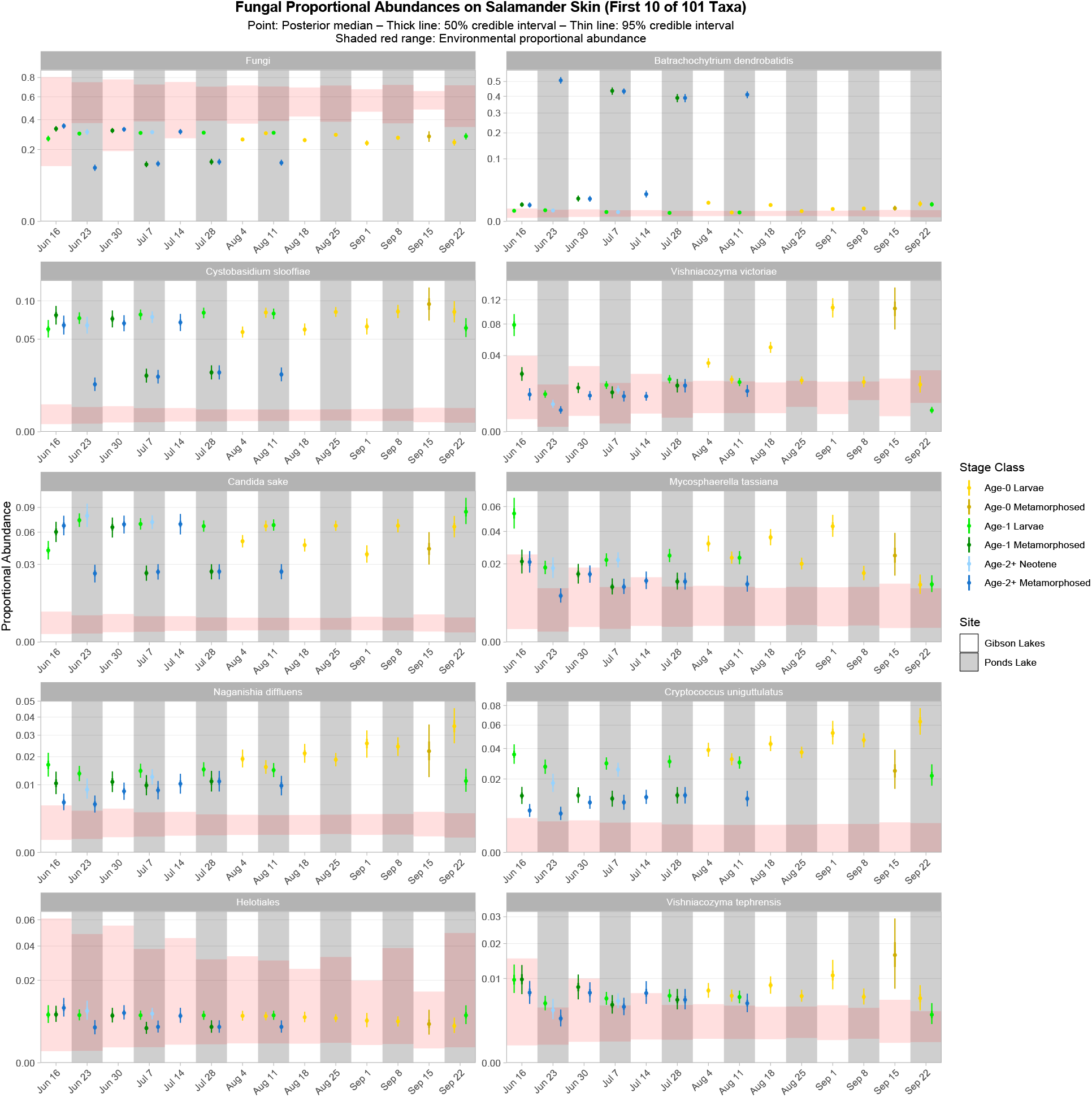
Proportional abundance predictions for the first ten fungal taxa from the top 100. Points, thick lines, and thin lines represent posterior medians, 50% credible intervals, and 95% credible intervals, respectively. Shaded red ranges represent environmental proportional abundances. Note the square root scale on the y-axis. Taxa are ordered (left to right, top to bottom) by descending average proportion of reads in salamander samples in which each combination of site and life stage receives equal weight. Proportional abundance prediction plots for the remaining top 100 fungal taxa can be found in the Supplemental Information.

93 of the top 100 fungal taxa detected in salamander samples were also detected in environmental samples. We detected 67 and 91 of the top 100 fungal taxa in water and substrate samples, respectively. 27 of the top 100 fungal taxa were disproportionately more abundant on salamander skin relative to the environment for the majority of combinations of stage class and sampling event (Figures 5 & S17-S21), including *Candida sake*, *Wallemia muriae*, and Vishniacozyma. Taxa detected exclusively on salamander skin include Pleosporales, *Melanodiplodia tianschanica*, and *Buckleyzyma aurantiaca*. Four of the top 100 fungal taxa had lower proportional abundances on salamander skin than in the environment for the majority of combinations of stage class and sampling event, including Ascomycota, Basidiomycota, and Rozellomycota.

Patterns of microbial diversity for fungi differed than those for bacteria. Salamander fungal diversity was highest among age-0 larvae at Gibson Lakes, age-1 larvae at the beginning of the season at Ponds Lake, and late-season larvae at Ponds Lake (Figure 6). Fungal diversity increased throughout the early season for metamorphosed individuals at both lakes, with metamorphosed individuals at Ponds Lake having lower fungal diversity than at Gibson Lakes, possibly due to the high proportional abundance of *Bd* in Ponds Lake metamorphosed salamanders. Similar to the bacterial community dissimilarity predictions, for a given stage class, salamander fungal communities were more dissimilar between sites than through time, with this pattern being more pronounced for age-2+ metamorphosed than age-0 larval individuals (Figure S22).

**Figure 6.**
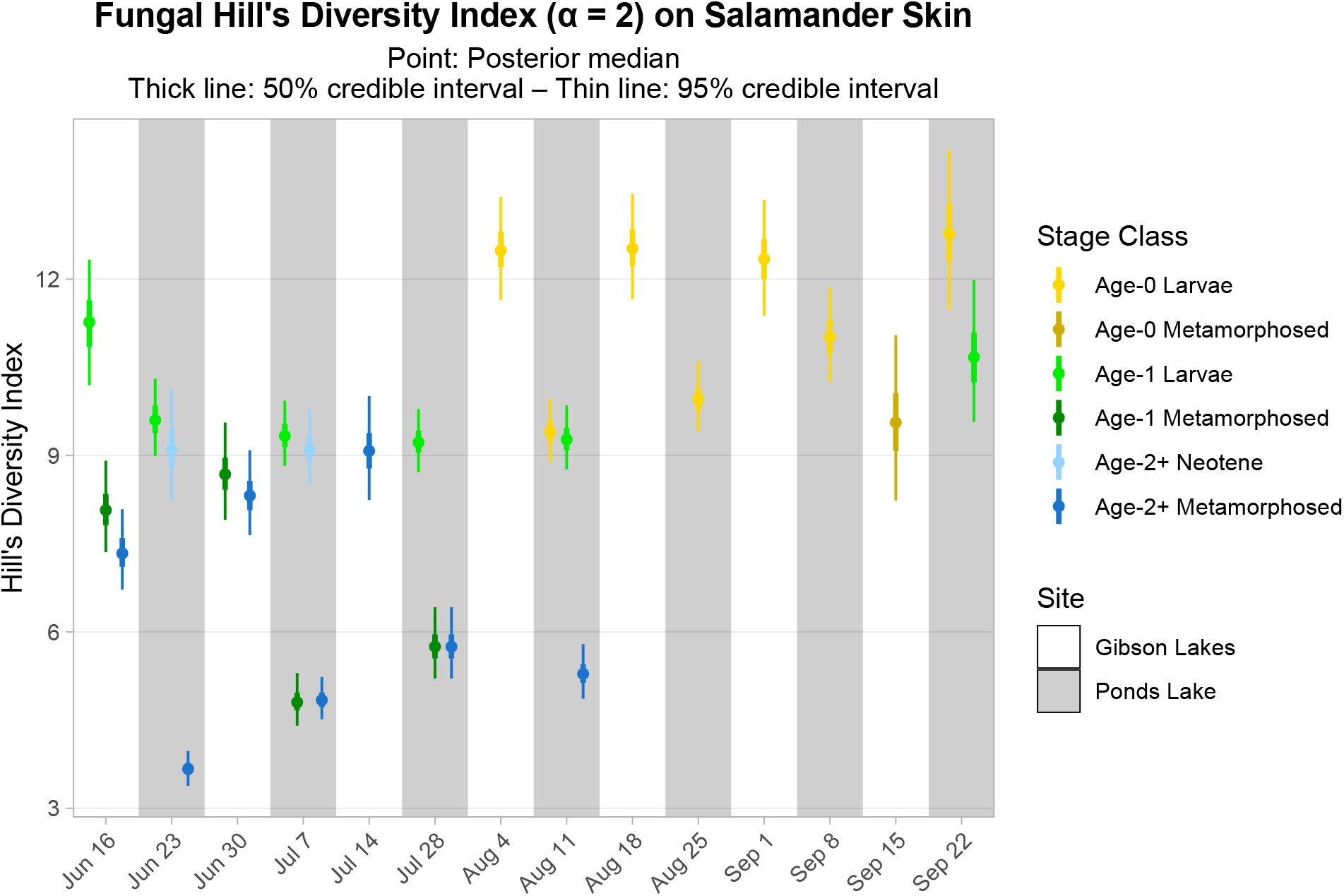
Hill’s diversity index (α = 2) for fungal communities on salamander skin. Points, thick lines, and thin lines represent posterior medians, 50% credible intervals, and 95% credible intervals, respectively.

Within the top 100 bacterial taxa, *Bd*-inhibitory taxa were disproportionately more abundant on salamander skin relative to the environment for all stage classes at both lakes throughout the season (Figure 7). Non-*Bd*-inhibitory bacterial taxa within the top 100 were disproportionately more abundant on salamander skin relative to the environment for most combinations of stage class and sampling event at Ponds Lake, but we were unable to detect differences in the proportional abundances of non-*Bd*-inhibitory bacterial taxa between salamander skin and the environment for any combination of stage class and sampling event at Gibson Lakes. The top 100 bacterial taxa of uncertain *Bd*-inhibition status were disproportionately more abundant on salamander skin compared to the environment for most combinations of stage class and sampling event, reflecting a broader trend among the top 100 bacterial taxa.

**Figure 7.**
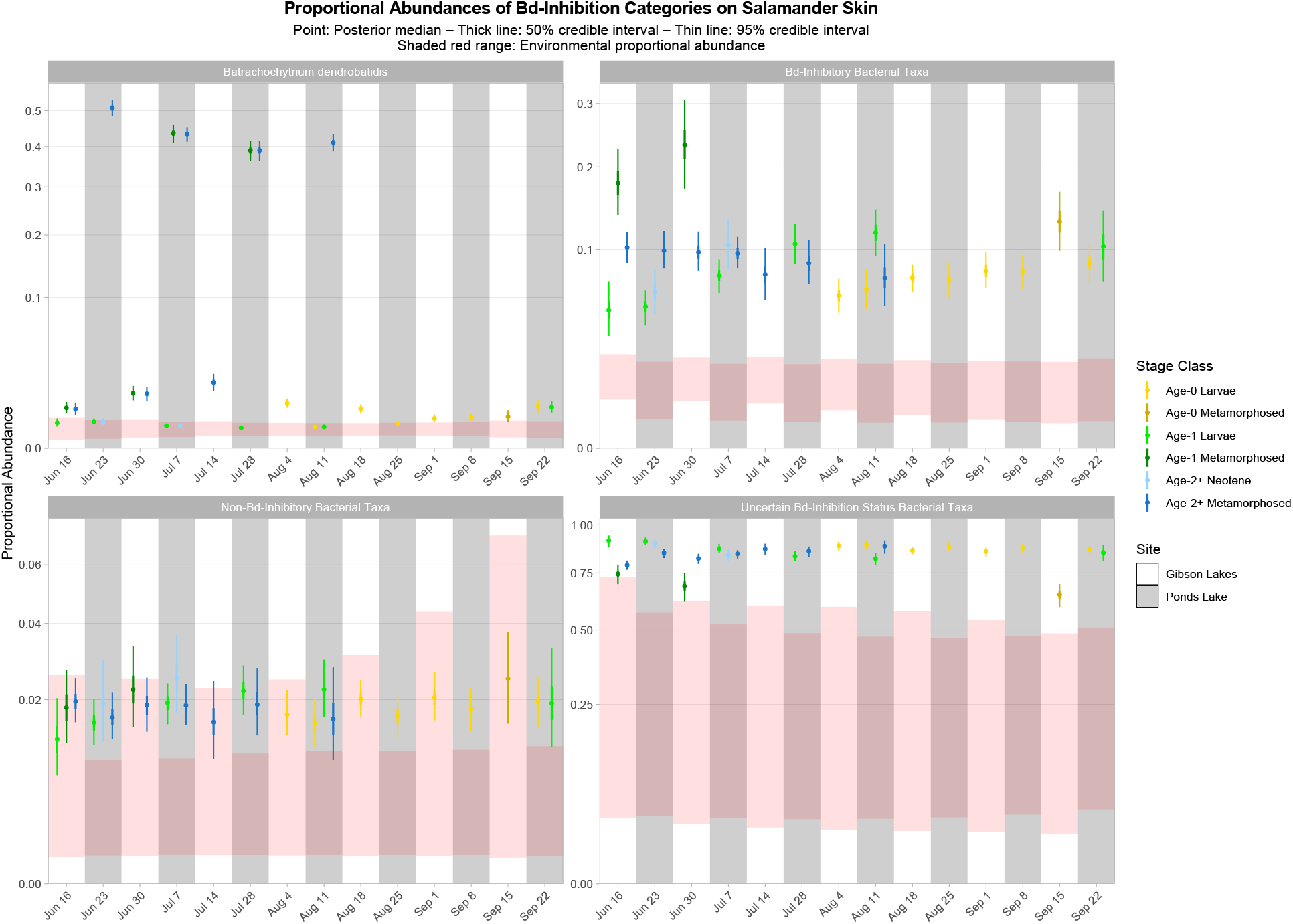
Proportional abundance predictions of *Batrachochytrium dendrobatidis* (*Bd*) and *Bd*-inhibitory bacterial taxa from the top 100. Points, thick lines, and thin lines represent posterior medians, 50% credible intervals, and 95% credible intervals, respectively. Shaded red ranges represent environmental proportional abundances. Note the square root scale on the y-axis.

### Microbial Absolute Abundance

As in our predictions of microbial composition, the densities of the many of the top 100 bacterial taxa exhibited spatiotemporal and ontogenetic variation (Figures S23-S27). While overlapping credible intervals complicate interpretation, some emerging trends can be observed. The density of Rhodoferax 1 appears to decrease over time. The densities of some taxa exhibit a possible mid-season slump, where density drops in the early season and then rises during the later season (*e.g.*, Sanguibacter 1, Stenotrophomonas 1, Pseudomonas 4, *Pseudochrobactrum kiredjianiae* 1, and Comamonadaceae 28). We observed higher densities of FukuN18 freshwater group 1 and hgcI clade 2 at Ponds Lake, and inversely, we observed higher densities of *Polynucleobacter difficilis* 1 at Gibson Lakes. Parachlamydiaceae 1 and Comamonadaceae 13 had higher densities on larvae than metamorphosed individuals, and Comamonadaceae 3 had higher densities on age-2+ metamorphosed individuals at Ponds Lake than other stage class-site combinations.

We were unable to detect spatiotemporal or ontogenetic variation in the densities of many of the top 100 fungal taxa (Figures S28-S32), possibly due to the strong control over overfitting of our LASSO models and the poor taxonomic resolution of many fungal taxa (Figure S8). The density of *Bd* was generally highest on age-1 and age-2+ metamorphosed individuals at Ponds Lake. While many credible intervals overlap, the density of *Bd*-inhibitory bacterial taxa within the top 100 shows a potential mid-season slump (Figure S33).

## DISCUSSION

We observed spatiotemporal and ontogenetic variation in the relative abundances and microbial diversity of bacterial and fungal taxa in the skin-associated microbiome of the western tiger salamander at two high alpine Rocky Mountain lakes. Our best-fitting models for microbial composition included all predictors or their interactions except for the model of fungal communities on salamander skin, for which the stratum predictor was excluded. The inclusion of stratum as a predictor in the best-fitting composition model of bacteria on salamander skin, as well as for composition models of bacteria and fungi in the environment, suggests that we observed spatial variation in microbial community composition within lakes in addition to between lakes. For salamander skin-associated bacterial and fungal communities, composition was better explained by spatiotemporal than water quality covariates, and we observed greater community dissimilarity between lakes than through time for each salamander stage class. In agreement with other amphibian skin-associated microbiome studies, we found that the skin of the western tiger salamander is a selective environment with taxa disproportionately represented compared to their relative abundances in water and substrate (Kueneman *et al*. 2013; Walke *et al*. 2014; Bletz *et al*. 2017a).

We detected *Bd* on salamander skin at both lakes, with the relative and absolute abundances of *Bd* being highest for age-1 and age-2+ metamorphosed salamanders at Ponds Lake. The higher abundance of *Bd* on the skin of metamorphosed compared to larval amphibians is supported by other studies and is thought to be the result of increased keratin, a substrate for *Bd*, in amphibian skin following metamorphosis, during which structural changes to the skin occur (Berger *et al*. 1998; Marantelli *et al*. 2004). We are unsure why differences in the relative abundance of *Bd* is much less pronounced between larval and metamorphosed individuals at Gibson Lakes. Since *Bd* was absent in all negative control samples, we are confident that *Bd* was present at Gibson Lakes and that its detection was not the result of contamination from Ponds Lake samples.

We observed that *Bd*-inhibitory bacterial taxa which were members of the top 100 were disproportionately more abundant on salamander skin relative to the environment for all stage classes at both lakes and throughout the season. If members of the top 100 bacterial taxa for which we have high confidence in their *Bd*-inhibition statuses can be considered a random sample of our observed bacterial taxa, then this could be taken as evidence that salamander skin selects for *Bd*-inhibitory bacteria. While common bacterial taxa on salamander skin (*i.e.*, our top 100) tend to have disproportionately higher abundances compared to the environment, the apparent selection for *Bd*-inhibitory taxa is much stronger than for non-*Bd*-inhibitory taxa, suggesting that selection for *Bd*-inhibitory taxa may indeed be occurring. We did not, however, observe correlations between the relative or absolute abundances of *Bd*-inhibitory bacteria and *Bd*.

For both relative and absolute abundances, we observed a strong negative correlation between *Bd* and *Cystobasidium slooffiae*, and we observed strong positive correlations between *Bd* and Comamonadaceae 3 and 6. Comamonadaceae has been found to be abundant on the skin of multiple amphibian species, including the tiger salamander (McKenzie *et al*. 2011), and some members show evidence of *Bd*-inhibition or negative co-occurrence with fungal taxa (Woodhams *et al*. 2015; Kueneman *et al*. 2016b). Despite this, both our study and Walke *et al*. (2015) found positive or very weak correlations between members of Comamonadaceae and *Bd*. While we were unable to confidently predict the *Bd*-inhibition status of Comamonadaceae 3 and 6, we did predict one member of Comamonadaceae to be *Bd*-inhibitory (Comamonadaceae 5). Still, we observed no correlation between the relative or absolute abundances of this taxon and *Bd*. We note that, despite harboring *Bd*, tiger salamanders have been found to be resistant to chytridiomycosis (Davidson *et al*. 2003).

A key aim of amphibian skin-associated microbiome studies relates to understanding what role microbial communities play in protecting their hosts against cutaneous diseases such as chytridiomycosis. While DNA metabarcoding is commonly employed to characterize the composition of microbial communities, we experienced challenges relating community composition to functional activity. Using 16S rRNA gene sequences, we were unable to predict *Bd*-inhibition statuses for the vast majority of our bacterial taxa with any reasonable certainty. This is not surprising given that, after trimming to our amplicon region, the majority of sequences in the Woodhams database belonged to multiple bacterial isolates with variable antifungal statuses, and these isolates provided limited phylogenetic coverage of our bacterial taxa. Similarly, Becker *et al*. (2015) found bacterial congeners to frequently range from complete inhibition to facilitation of *Bd*. Another approach to exploring the functional activity of microbial communities involves metatranscriptomics, the sequencing of RNA within a microbiome to investigate gene expression (Nichols & Davenport 2021). With a metatranscriptomics approach to exploring functional activity, antifungal secondary metabolite production by microbes experiencing real-world biotic and abiotic conditions on salamander skin could be observed, and a precise knowledge of community composition, while still informative, would not be a pre-requisite for inference.

We highlight that our Bayesian Dirichlet-multinomial regression and negative binomial LASSO models provide readily-interpretable predictions of microbial relative and absolute abundances. We believe that these are valuable analytical tools, and in this study, we show how these models can be applied to amphibian skin-associated microbial ecology, including inference for differential abundance, microbial selection, and alpha and beta diversity. While existing applications of the Woodhams antifungal isolates database tend to use similarity clustering or local alignments to categorize bacterial taxa as “potentially” *Bd*-inhibitory (*e.g.*, Kueneman *et al*. 2016b; Bletz *et al*. 2017a; Kruger 2020), our use of stochastic character mapping provides the benefit of yielding probabilistic predictions that bacterial taxa are actually *Bd*-inhibitory.

We caution that our absolute abundance models rely on assumptions of salamander growth and swabbing intensity. In deriving estimates of swabbed area from salamander length, we assumed that height and width grow proportionally. While swabbing, the 10 strokes along the length of the belly were distributed across the belly’s width. This means that the same belly area was swabbed more times for smaller salamanders than for larger salamanders. For inferences from our absolute abundance models to be valid, an assumption must be made that the number of microbes collected asymptotes after a certain swabbing intensity, and we must further assume that we reached this threshold of swabbing intensity. We suggest that the need for these assumptions can be avoided by using a different swabbing protocol. For example, instead of stroking a swab across the ventral surface a certain number of times while covering an area of interest, one could swab the full area of interest (*e.g.*, the belly) a certain number of times and measure the swabbed area, a method which is already applied in studies of *Bd* load (North & Alford 2008).

Our study emphasizes two traditionally understudied areas of amphibian skin-associated microbial ecology, temporal variation in community composition and expanding our view of the microbiome to include fungi in addition to bacteria. Temporal variation in community composition could prove challenging for studies examining spatial variation, where temporal and spatial variation may be confounded. We also identified additional sources of variation in community composition which are not typically considered. Within life stages, we identified additional variation with salamander age, and within lakes, we identified additional variation between strata. Furthermore, we observed that the relationships between community composition and spatiotemporal and stage class covariates are interdependent, complex, and best described using interactions.

Through this study, we have gained a greater understanding of microbial ecology on amphibian skin through the examination of season-long temporal variation of bacterial and fungal communities. In addition to identifying further sources of variation in community composition, we have identified differentially abundant taxa, have examined salamander microbial selection, have investigated alpha and beta diversity in microbial communities, and have predicted antifungal activity. Ultimately, this ecological knowledge may assist in the conservation of amphibian species threatened by chytridiomycosis.

## ACKNOWLEDGEMENTS

We would like to thank our volunteers for fieldwork assistance, with special thanks to Sarah Shaw. We would also like to thank Trisha Atwood, Bonnie Waring, and Kevin Landom for their support and feedback. This project was supported with funding from the Utah State University (USU) Quinney College of Natural Resources Undergraduate Research Grant, the USU Office of Research Undergraduate Research and Creative Opportunities Grant, the USU Honors Program Honors Research Fund, the Society for the Study of Amphibians and Reptiles Roger Conant Undergraduate Research Grant in Herpetology, and the National Science Foundation (DBI 1638768 to Z.G.). We thank the Uinta-Wasatch-Cache and Caribou-Targhee National Forests for permission to conduct fieldwork on National Forest lands. The support and resources from the Center for High Performance Computing at the University of Utah are gratefully acknowledged.

## DATA ACCESSIBILITY

Raw sequence reads are deposited in the Sequence Read Archive (BioProject PRJNA843333). Sample metadata will be made available on Dryad. Scripts used in the analyses are available on GitHub (https://github.com/Urodelan/2022_Salamander_Microbiome).

## AUTHOR CONTRIBUTIONS

K.B.G. and Z.G. designed the research. K.B.G. and J.D.H. conducted field sampling. K.B.G. performed the lab work and analyses. K.B.G. and J.D.H. wrote the manuscript with revisions from all authors. Z.G. provided guidance, support, and feedback at all stages of the project.

## Notes

### Competing Interest Statement

The authors have declared no competing interest.

