## Supplement for "Spatiotemporal and ontogenetic variation, microbial selection, and predicted antifungal function in the skin-associated microbiome of a Rocky Mountain amphibian"

### Methods

#### *SNOTEL snowmelt timing data*

Although Ponds Lake was not visited early enough to observe salamander eggs, snowmelt data suggests that salamanders likely laid eggs at about the same time at both study sites. Data from a NRCS SNOTEL site (#828) within ~1.2 km distance and ~50 m elevation from Ponds Lake first reported a 2018 snow water equivalent of zero on May $24^{\text{th}}$, and data from a SNOTEL site (#484) within ~2.9 km and ~80 m elevation from Gibson Lakes first reported a 2018 snow water equivalent of zero on May $23^{\text{rd}}$. The 30-year median snow water equivalent reaches zero at the SNOTEL sites near Ponds Lake and Gibson Lakes on June $11^{\text{th}}$ and June $4^{\text{th}}$, respectively. Taken together, this suggests that snow melted at these lakes at about the same time in 2018, possibly within days of each other, and snow typically melts at these lakes about a week apart.

#### *Salamander length and weight*

Salamander length and weight measurements were taken to verify age classes (Figure S1).

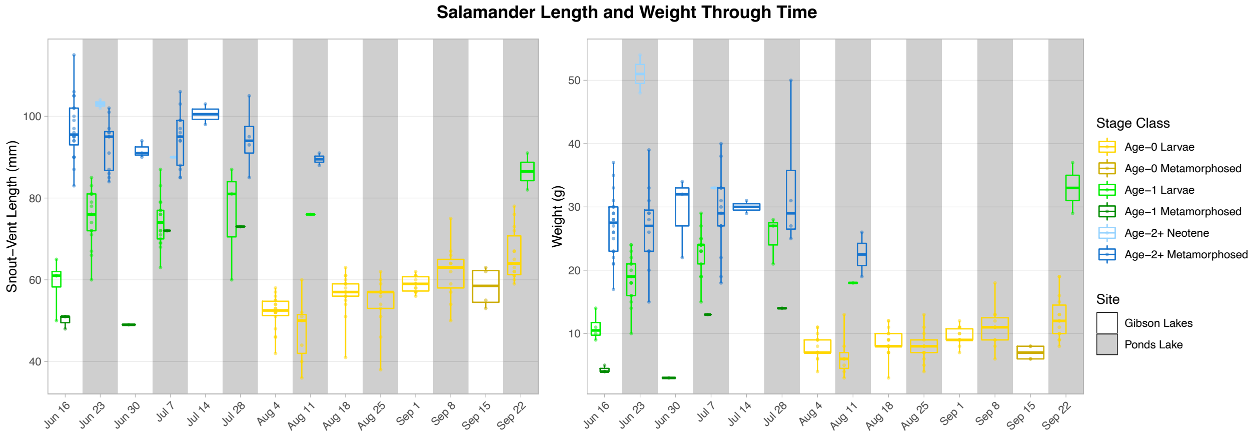

Figure S1. Salamander length and weight through time.

#### *Library preparation*

Library preparation for sequencing the 16S rRNA and ITS genetic barcoding regions for bacteria and fungi involved two-stage polymerase chain reaction (PCR) with MagBead cleanups in between. First stage PCR amplified the 16S and ITS genes, added unique dual index combinations to each sample, and added a portion of the Illumina Nextera adapter. Second stage PCR completed Illumina adapter addition. The 16S rRNA V4 region was amplified using the primers 515F (forward; Parada *et al.* 2016) and 806R (reverse; Caporaso 2011). The ITS1 region was amplified using the primers ITS1-F (forward; Gardes & Bruns 1993) and ITS2 (reverse; White *et al.* 1990). First stage PCR was a 15 µL reaction with 3 µL 5X Phusion HF Buffer, 0.45 µL 10M dNTPs, 0.3 µL Kapa HiFi HotStart DNA Polymerase, 3.25 µL high-performance liquid chromatography-grade water, 6 µL appropriate 0.25 µM paired primers, and 2 µL template DNA. First stage PCR was performed with the settings: 95 °C for 3 min, 15 cycles at 98 °C for 30 s, 15 cycles at 62 °C for 30 s, 15 cycles at 72 °C for 30 s, 72 °C for 5 min, and 4 °C for ever. Two first-stage PCR replicates were performed for each sample. Following first stage PCR, replicates were pooled together for each sample (*i.e.*, 15 µL of one replicate was transferred to the other) and purified using a modified manual AxyPrep MagBead PCR Clean-up protocol. For this protocol, the MagBead bottle was gently shaken to resuspend particles and equilibrated to room temperature. 24 µL of MagBead solution was added to each sample and pipette mixed 10 times followed by a 5-min incubation at room temperature. Samples were placed on a magnet plate and incubated for 5 min at room temperature or until the solutions were clear. The solutions were aspirated and discarded, and the beads were rinsed twice with 100 µL fresh 80% ethanol. The rinse involved adding ethanol, incubating for 30 s at room temperature, and aspirating and discarding the ethanol. Samples were re-aspirated to assure maximum ethanol removal and air dried for 7 min. Samples were removed from the magnet plate, 40 µL TE buffer was added to each sample and pipette mixed 10 times, and samples were incubated for 2 min at room temperature. Samples were placed back on the magnet plate for 5 min or until the solutions were clear, and 10 µL of solution was transferred for use as template DNA in the second stage PCR reaction.

Second stage PCR was a 15 µL reaction with 3 µL 5X Phusion HF Buffer, 0.45 µL 10M dNTPs, 0.3 µL Kapa HiFi HotStart DNA Polymerase, 0.5 µL 10 µM forward and reverse flow cell primers, 0.75 µL high-performance liquid chromatography-grade water, and 10 µL template DNA. Second stage PCR was performed with the settings: 95 °C for 3 min, 19 cycles at 98 °C for 30 s, 19 cycles at 55 °C for 30 s, 19 cycles at 72 °C for 30 s, 72 °C for 5 min, and 4 °C for ever. Following second stage PCR, 15 µL water was added to each sample. Samples were then purified using the same modified AxyPrep MagBead PCR Clean-up protocol as used before, except for 40 µL of solution (instead of 10 µL) was transferred during the final step to serve as the cleaned amplicon product.

Amplicon product concentrations were measured via absorption with Synergy HTX Take 3 Trio. Since amplicon product concentrations were of relatively similar concentrations (within an order of magnitude), 2 µL of each sample was combined across 8 tubes. The 8 tubes were vortexed and spun before 50 µL of each was combined into a final pooled tube.

#### *DADA2 bioinformatics pipeline*

Read pairs were quality filtered using DADA2's (version 3.10; Callahan *et al.* 2016) filterAndTrim function (truncating reads at the first instance of a quality score ≤ 2, discarding reads with more than 2 expected errors after truncation, discarding reads of less than 10 bp after truncation, and discarding reads which match against the phiX genome). DADA2's error-rate learning (learnErrors) and application of the core sample inference algorithm (dada) were performed using default settings. For the MiSeq 16S dataset, read pairs were merged using DADA2's mergePairs function (minimum overlap of 12 bp and allowing for no mismatches in the overlap region). For the MiSeq ITS dataset, which has variable length amplicon sequences whose read pairs may not overlap, mergePairs was applied as above but with rejects returned (returnRejects = TRUE). For unsuccessfully merged MiSeq ITS pairs, read pairs were concatenated (mergePairs with justConcatenate = TRUE) with a 10-N spacer between them if the overlap region was less than 12 bp (the DADA2 pipeline is made to work with these 10-N spacers), and unsuccessfully merged pairs with mismatches in an overlap region of at least 12 bp were discarded (following DADA2's default behavior, we consider 12 bp to be a true overlap). Concatenated MiSeq ITS read pairs which were of forward or reverse reads which were previously truncated were discarded (to ensure unique non-overlapping sequences do not appear different because of differing lengths). To remove potential partial overlap between the remaining concatenated MiSeq ITS read pairs, the maximum possible length of the overlap region (the sum of the number of matches, number of mismatches, and number of indels in the overlap region from the mergePairs output) for each of these read pairs was trimmed from the reverse read segment on the end which could potentially overlap with the forward read. MiSeq sequences which were the synthgenes, coligos, or had lengths between or equal to 250 and 256 bp for 16S and greater than 125 bp for ITS were kept. Chimeric reads were removed for each of the MiSeq 16S and ITS datasets using DADA2's removeBimeraDenovo function with default settings, and synthgene and coligo sequences were subsequently removed. A naïve Bayesian classifier (Wang *et al.* 2007) was used to classify each of the MiSeq 16S and ITS datasets with DADA2's assignTaxonomy function using Silva (formatted by DADA2 with down to genus-level taxonomies; version 138; Quast *et al.* 2012) and UNITE (general dynamic FASTA release for fungi; version 8.2; Nilsson *et al.* 2019) reference libraries with default settings. For the MiSeq 16S dataset, species-level assignments were made through exact matching against a DADA2-maintained species-level Silva database using DADA2's addSpecies function with default settings. The MiSeq 16S dataset was subsetted to sequences which were classified as bacteria, and the MiSeq ITS dataset was subsetted to sequences which were classified as fungi.

#### *Creating study-specific reference libraries*

To create study-specific 16S and ITS reference libraries for classifying the NextSeq sequences, forward and reverse reads were created from the classified MiSeq 16S and ITS sequences which were the same lengths of the forward and reverse NextSeq 16S and ITS reads. The Biostrings R package (version 2.54.0; Pagès *et al.* 2019) was used to generate reverse complements of the MiSeq sequences during the creation of reverse reads. For the MiSeq 16S data, most truncated reference sequences were still unique after trimming reads to NextSeq lengths, with 104 out of 16,845 reference sequences becoming duplicates of each other. In R, these 104 non-unique truncated reference sequences were collapsed into 52 unique sequences and assigned a consensus taxonomy and consensus MiSeq-length sequence (for later use in alignment and antifungal prediction). The consensus taxonomy was the most precise agreed upon taxonomy of the duplicated reference sequences. The consensus MiSeq-length sequences were created by replacing inconsistencies in the full-length reference sequences with the most parsimonious ambiguous nucleotide (all differences were substitutions). For the MiSeq ITS data, the same process was applied for creating consensus taxonomies for 87 non-unique truncated reference sequences out of 3,954 total. Since a reference taxon can occur multiple times in the reference libraries (being associated with multiple amplicon sequence variants [ASVs]), reference taxon names were appended with integers so that all reference taxa names were unique. Forward and reverse reference reads were also created for the 16S and ITS coligos and sythgenes, which were added to their respective study-specific reference libraries.

#### *Mock community visualization*

The proportional abundances (excluding the synthgenes from the calculations and after summing read counts across PCR replicates) of taxa detected in the mock community samples were visualized (Figure S2). The actual DNA composition of the mock communities were 12% *Listeria monocytogenes* (bacterium), 12% *Pseudomonas aeruginosa* (bacterium), 12% *Bacillus subtilis* (bacterium), 12% *Escherichia coli* (bacterium), 12% *Salmonella enterica* (bacterium), 12% *Lactobacillus fermentum* (bacterium), 12% *Enterococcus faecalis* (bacterium), 12% *Staphylococcus aureus* (bacterium), 2% *Saccharomyces cerevisiae* (fungus), and 2% *Cryptococcus neoformans* (fungus). Given this composition, we expected eight bacterial taxa in the 16S mock community data with uniform proportional abundances, and we expected two fungal taxa in the ITS mock community data also with uniform proportional abundances. We observed some amplification bias in the 16S data, and two 16S taxa (*Bacillus* and *Salmonella*) were split into two ASVs each (with the degree of the split being more pronounced for *Bacillus*). In our ITS mock community data, we observed much more extreme amplification bias, and one taxon (*Saccharomyces cerevisiae*) was split into three substantial ASVs. In an effort to mitigate the potential impact of fungal taxa being split into multiple ASVs, we merged fungal ASVs which were assigned the same taxonomy into the same taxa (and trailing integers in the ITS taxa names were removed).

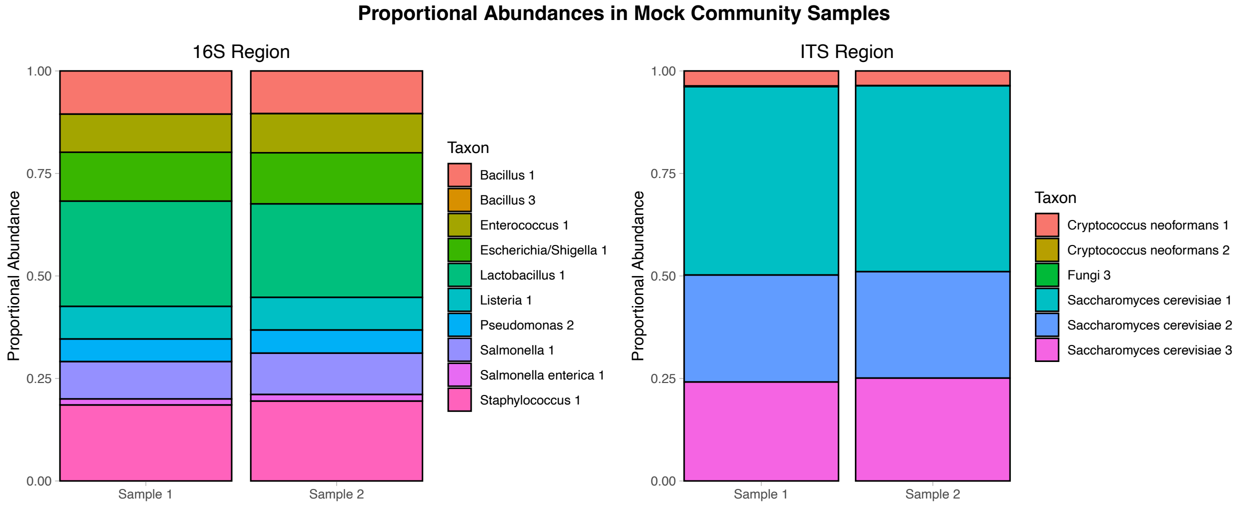

Figure S2. Observed composition of mock community samples.

#### *Quality assessment of microbiome data*

We performed principal component analyses (PCAs) on the proportional abundances of sample taxa (excluding the synthgenes from the calculations) across PCR replicates (Figure S3). The PCA plots on PCR replicate for both 16S and ITS samples suggest that taxa proportional abundances in both PCR replicates were similar, and read counts were summed across PCR replicates for each sample. We then performed PCAs on the proportional abundances of sample taxa (excluding the synthgenes from the calculations) across sample types (*i.e.*, salamander, wet swab, dry swab, blank, water, water control, substrate, and mock community samples) to check for cross contamination between samples preceding library preparation. We performed separate 16S and ITS PCAs with random subsets of salamander samples (Figures S4 & S5) so the number of salamander, water, and substrate samples included in the PCAs was similar (51 to 67 of each type). There were 10 salamander samples which grouped closely with wet swab and dry swab negative controls in the 16S PCA plots, so these samples were removed from the 16S data for all subsequent analyses.

Following Harrison *et al.* (2021), we calculated point estimates for the absolute abundances of microbial taxa relative to the absolute abundance of the synthgene in each of the 16S and ITS datasets. There were 13 samples in which the synthgene was not detected (all were non-control water and substrate samples), and the synthgene read counts for these samples were set to one for the purpose of estimating taxa absolute abundances (this only affects the following visualization). We visualized total absolute abundances along with total read counts, both excluding the synthgene in sample totals, within each sample (Figure S6), and we observed higher point estimates of absolute microbial abundance for field samples (*i.e.*, non-control salamander, water, and substrate samples) than for their associated negative controls.

While we observed 16S sequences classified as mitochondrial and chloroplast DNA, the composition of these sequences was minor (2.6% of salamander sample reads). We used online nucleotide BLAST (default settings; Zhang *et al.* 2000) to evaluate whether the mitochondrial and chloroplast sequences with the most salamander sample reads could be bacterial. A BLAST of the MiSeq-length Mitochondria 1 sequence yielded four top hits with > 97% identity. Two hits were for uncultured bacteria while the others were for Aphanomyces water molds. Similarly, a BLAST of the MiSeq-length Chloroplast 7 sequence yielded exact hits to uncultured bacteria in addition to plant chloroplast. Since amphibian skin microbiome studies do not typically remove mitochondrial or chloroplast sequences (*e.g.*, Longo *et al.* 2015; Kueneman *et al.* 2016b; Bletz *et al.* 2017a; Kruger 2020), and because it is possible that these sequences represent uncultured bacteria, we retain these sequences in our analyses.

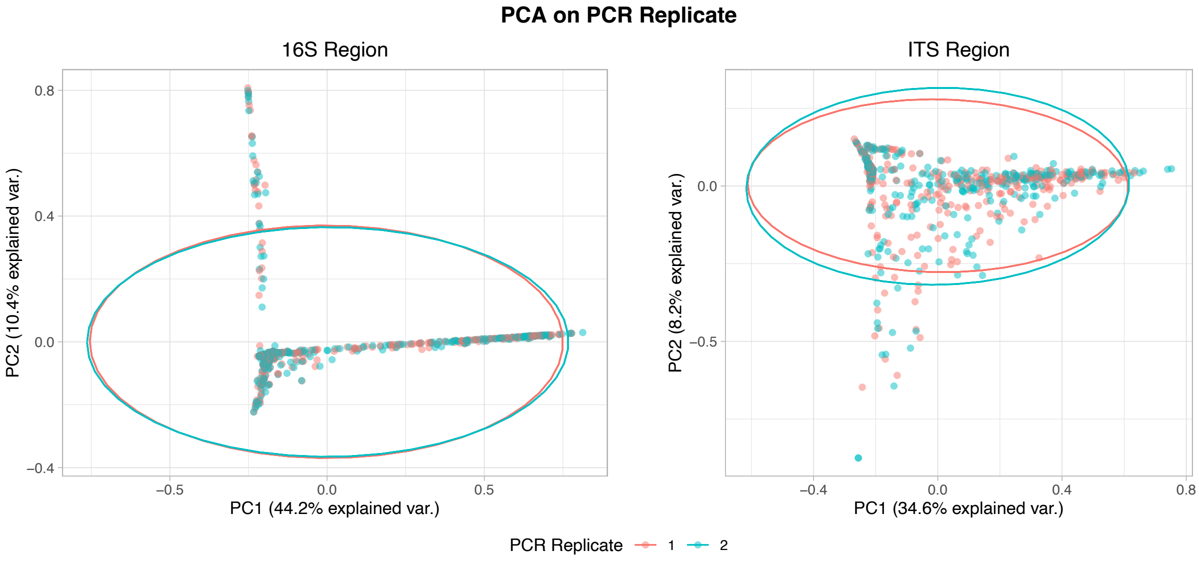

Figure S3. Principal component analyses (PCAs) on PCR replicate with 95% ellipses.

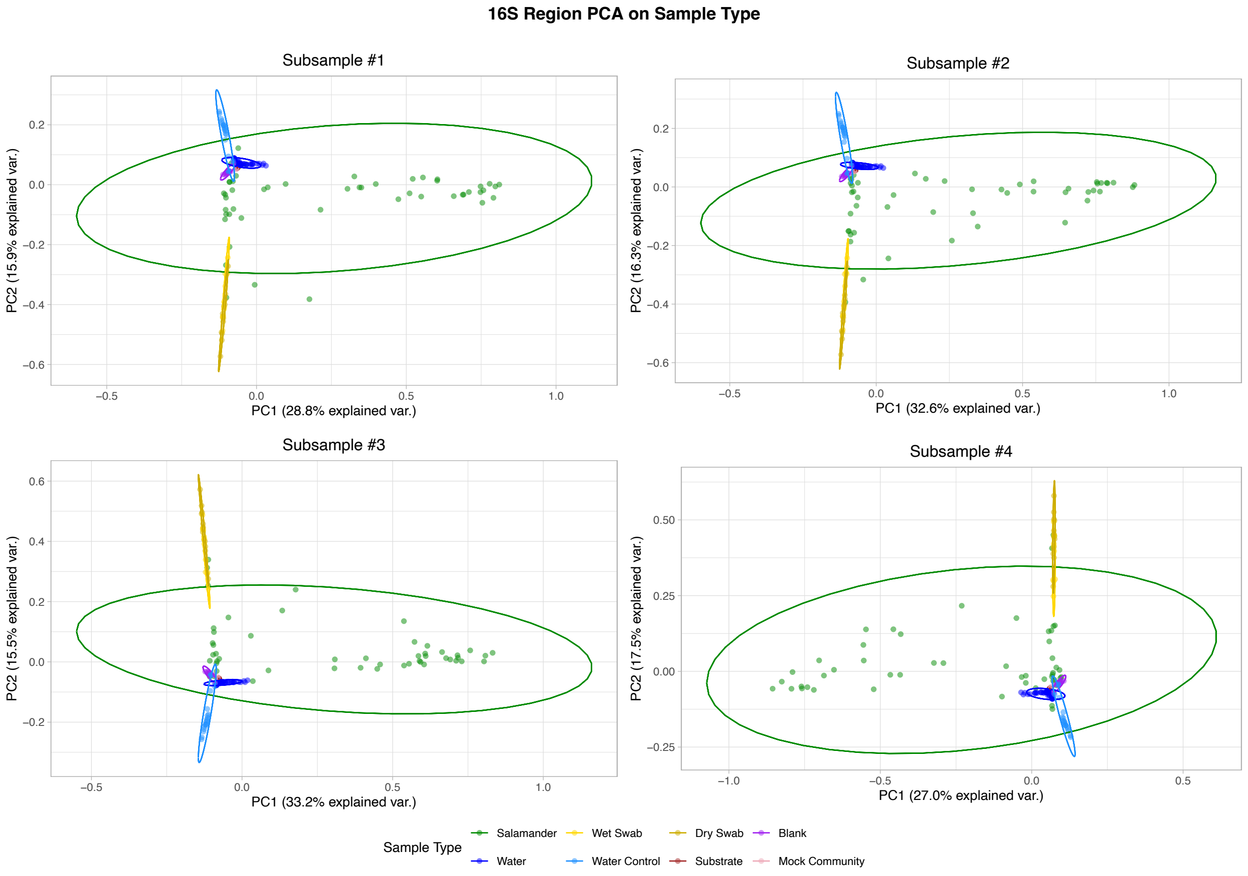

Figure S4. Principal component analyses (PCAs) on sample type with 95% ellipses for 16S data. The PCA in each plot contains a random subset of salamander samples.

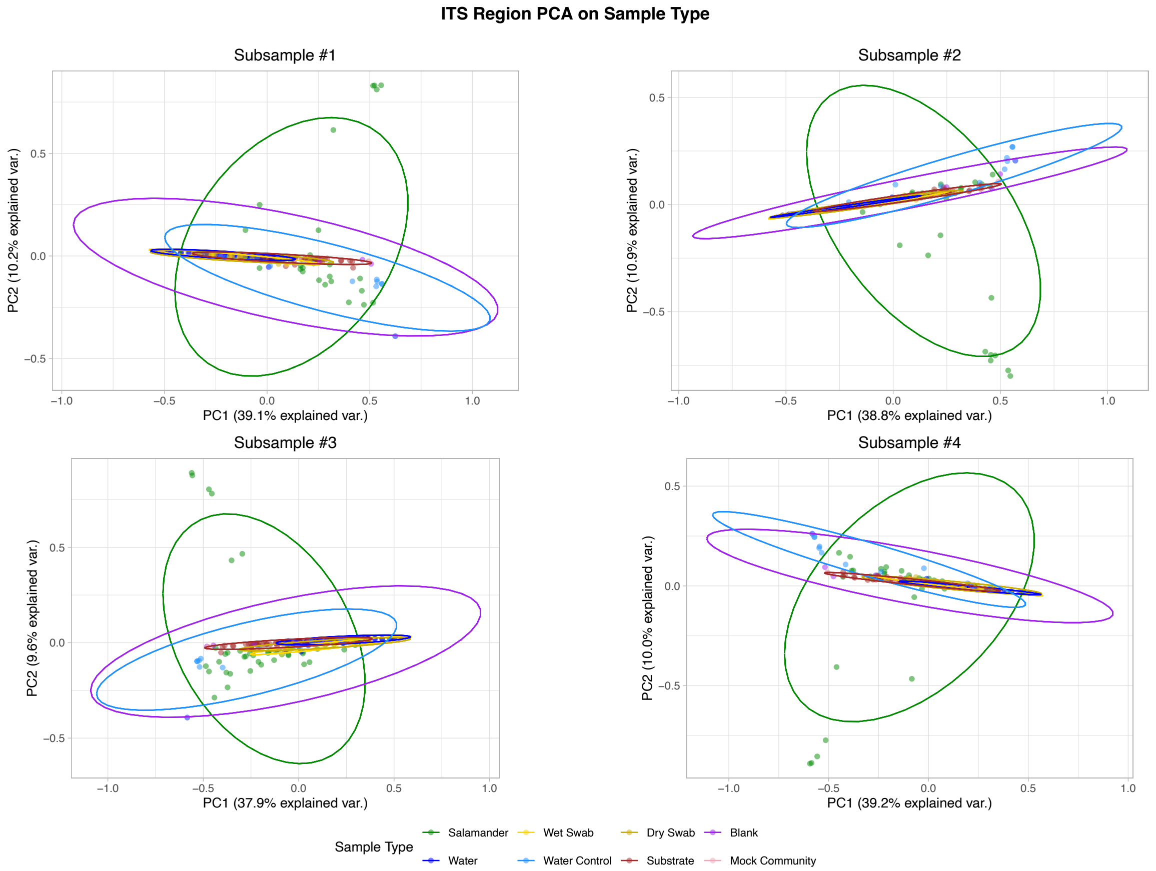

Figure S5. Principal component analyses (PCAs) on sample type with 95% ellipses for ITS data. The PCA in each plot contains a random subset of salamander samples.

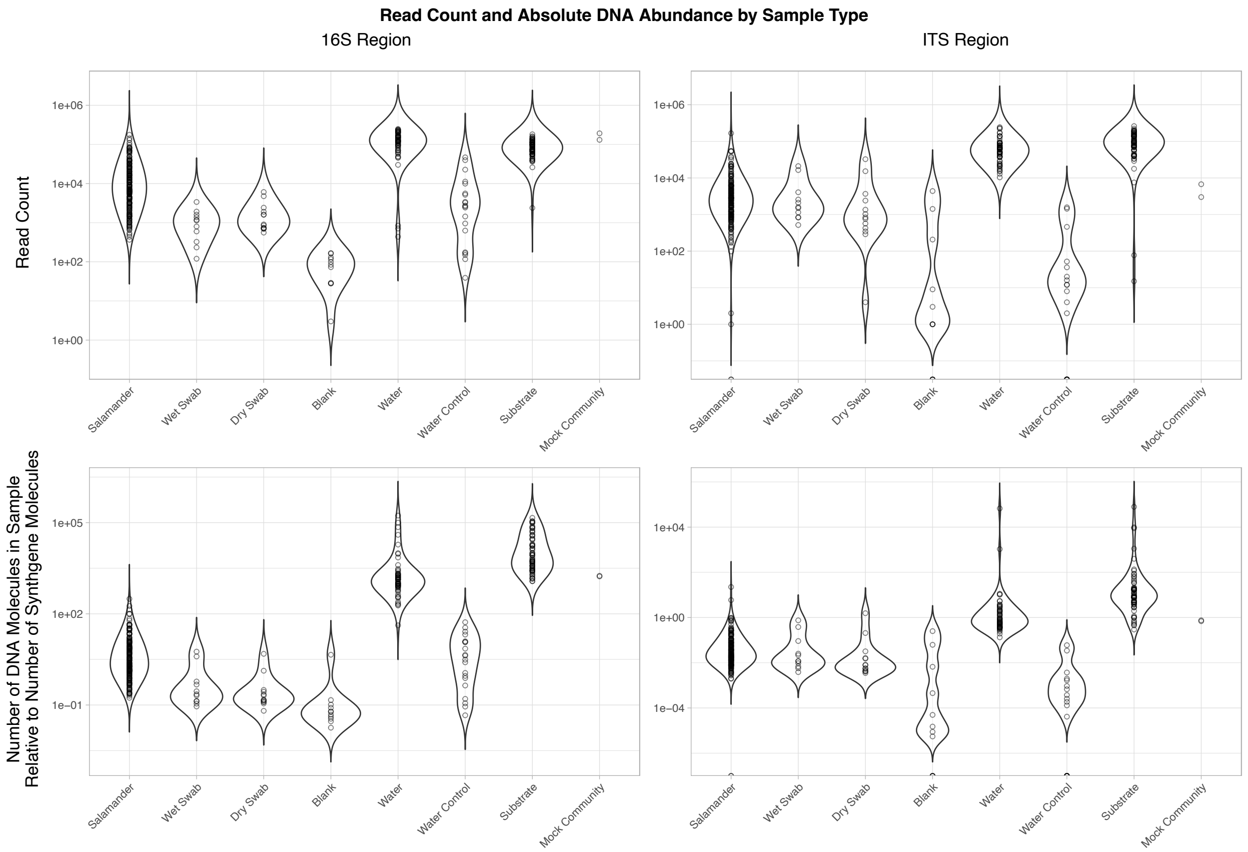

Figure S6. Sample read counts and point estimates of absolute DNA abundance by sample type and barcode region. Note the $\log_{10}$ scale on the y-axis.

#### *Antifungal prediction*

The Woodhams database of amphibian skin-associated microbiome antifungal bacterial isolates (Woodhams *et al.* 2015) was subsetted to just bacteria (using the UCLUST taxonomies provided in the database), and taxa with unknown *Batrachochytrium dendrobatidis* (*Bd*)-inhibition status were removed. The remaining Woodhams taxa that belonged to antifungal classes that were not *Bd*-inhibitory were reclassified as non-inhibitory. Woodhams sequences were trimmed to the 16S rRNA V4 region using our 16S amplification primers, and sequences without matches for both forward and reverse amplification primers were discarded. Trimmed Woodhams sequences which were outside the range of 250 to 256 bp were discarded. This left 872 Woodhams sequences available for alignment of the 1,944 sequences originally in the Woodhams database.

We used Clustal Omega (version 1.2.4; Sievers *et al.* 2011) with default settings to align the MiSeq 16S sequences of taxa detected in our NextSeq 16S non-control samples (*i.e.*, salamander, water, and substrate samples) with the Woodhams sequences, and we used FastTree 2 (version 2.1.11; Price *et al.* 2010) with a GTR+CAT model to create a phylogenetic tree. The tree was midpoint-rooted using midpoint.root in the phytools package (version 0.7.70; Revell 2012), and polytomies were randomly resolved into dichotomies (with zero-length branches) using multi2di in the ape package (version 5.4.1; Paradis & Schliep 2019). Edges of zero length were set to $10^{-6}$ times the total tree length. An equal-rates transition matrix was estimated with fitMk (phytools package), and a stochastic character mapping model was fit with this transition matrix using make.simmap (phytools package) and mclapply (parallel package) to simulate 5,000 phylogenetic trees with mapped *Bd*-inhibition statuses. Our NextSeq 16S taxa received a prior probability of being *Bd*-inhibitory of 0.5, while the Woodhams taxa received a zero (non-inhibitory) or a one (inhibitory). The posterior probabilities of our taxa being *Bd*-inhibitory were extracted from the fitted trees using describe.simmap in the phytools package.

##### *Microbial composition modeling*

Following Harrison *et al.* (2020), we added one to all read counts in the datasets (after the selection of the top 100 taxa and the creation of "other" categories) to avoid errors which occur when Dirichlet parameters in the model approach zero. Our Dirichlet-multinomial regression model is adapted from the Dirichlet regression model of Sennhenn-Reulen (2018), and our model takes the form:

Sample read counts $c_{i_k}$ are distributed according to the Dirichlet-multinomial distribution with expected proportions $p_{i_k}$ and precision parameter $\theta$, which controls the degree of overdispersion relative to the multinomial distribution.

$$c_{i_1} \dots c_{i_K} \sim \text{Dirichlet-multinomial}\left(p_{i_1} \dots p_{i_K} * e^{\theta}, \sum_{k=1}^{K} c_{ik}\right)$$

The softmax function normalizes linear predictor combinations $\eta_{i_k}$ into proportions $p_{i_k}$.

$$p_{i_1} \dots p_{i_K} = \text{softmax}(\eta_{i_1} \dots \eta_{i_K})$$

$$\text{softmax}(\eta_{i_1} \dots \eta_{i_K}) = \frac{e^{\eta_{i_1}}}{\sum_{k=1}^{K} e^{\eta_{i_k}}} \dots \frac{e^{\eta_{i_K}}}{\sum_{k=1}^{K} e^{\eta_{i_k}}}$$

$$\eta_{i_k} = \beta_{0_k} + B_{\text{Stratum}_k} * X_{\text{Stratum}_i}^{\top} + B_{\text{Non-stratum}_k} * X_{\text{Non-stratum}_i}^{\top}$$

$$B_{\text{Stratum}_k} = \left[\beta_{\text{Stratum}_{1_k}} \dots \beta_{\text{Stratum}_{S_k}}\right]$$

$$B_{\text{Non-stratum}_k} = \left[\beta_{\text{Non-stratum}_{1_k}} \dots \beta_{\text{Non-stratum}_{Q_k}}\right]$$

$$X_{\text{Stratum}_i} = \left[x_{\text{Stratum}_{1_i}} \dots x_{\text{Stratum}_{S_i}}\right]$$

$$X_{\text{Non-stratum}_i} = \left[x_{\text{Non-stratum}_{1_i}} \dots x_{\text{Non-stratum}_{Q_i}}\right]$$

The intercept terms $\beta_0$, stratum regression coefficients $\beta_{\text{Stratum}_s}$, and non-stratum regression coefficients $\beta_{\text{Non-stratum}_q}$ of the last taxon $K$ are set to zero to allow for the model to be identifiable.

$$\beta_{0_K} = 0$$

$$\beta_{\text{Stratum}_{s_K}} = 0$$

$$\beta_{\text{Non-stratum}_{q_K}} = 0$$

A normal prior is used on precision $\theta$.

$$\theta \sim \text{normal}(0, 1)$$

Except for the last taxon $K$, weakly informative priors are used on the intercept terms $\beta_{0_k}$ and non-stratum regression coefficients $\beta_{\text{Non-stratum}_{q_k}}$.

$$\beta_{0_{k \neq K}} \sim \text{normal}(0, 1)$$
$$\beta_{\text{Non-stratum}_{q_{k \neq K}}} \sim \text{normal}(0, 1)$$

Stratum is treated as a hierarchical effect. For each taxon except the last taxon $K$, stratum regression coefficients $\beta_{\text{Stratum}_{s_k}}$ share a common variance term ${\sigma^2}_{\text{Stratum}_k}$, except for the last stratum regression coefficient $\beta_{\text{Stratum}_{S_k}}$, which is set to the negative sum of the preceding stratum regression coefficients $\beta_{\text{Stratum}_{s \neq S_k}}$. This applies sum-to-zero constraints to allow for identifiability.

$$\beta_{\text{Stratum}_{s \neq S_{k \neq K}}} \sim \text{normal}\left(0, {\sigma^2}_{\text{Stratum}_k}\right)$$
$${\sigma^2}_{\text{Stratum}_k} \sim \text{inverse-gamma}(0.01, 0.01)$$
$$\beta_{\text{Stratum}_{S_{k \neq K}}} = -1 * \sum_{s=1}^{S-1} \beta_{\text{Stratum}_{s_k}}$$

Where $c_{i_k}$ is the read count,
$i = 1 \ldots N$ is the sample number,
$k = 1 \ldots K$ is the taxon number,
$s = 1 \ldots S$ is the stratum number,
$x_{\text{Stratum}_{s_i}}$ is the stratum predictor value,
$q = 1 \ldots Q$ is the non-stratum predictor number,
and $x_{\text{Non-stratum}_{q_i}}$ is the non-stratum predictor value.

For the salamander samples, stratum, site, and life stage were treated as categorical predictors and were coded as binary dummy variables. Stratum was coded as nine dummy variables to represent the four strata in Gibson Lakes and the five strata in Ponds Lake. These categorical predictors were mean-centered to aid in model convergence. Since mean centering does not affect the range in the dummy predictor values (*i.e.*, the range remains one), the interpretation of modeled effects is not impacted (Gelman 2008) and the sum-to-zero constraints on hierarchical stratum effects remain unaffected. Age and time were treated as continuous predictors. These predictors were mean-centered and scaled by one standard deviation to aid in model convergence and to reduce predictor collinearity. Age was coded from zero (age-0) to two (age-2+). Time was coded as the number of weeks since June 9th, 2018 and was also given a quadratic term. After centering, scaling, or both, interactions between all of the non-stratum predictors were created such that the non-stratum predictor matrix included four-way interactions between age, life stage, site, and a second-degree polynomial for week, all lower-level interactions, and the individual predictors.

Backwards variable selection by WAIC operated by fitting an initial model with all 24 predictors and calculating the model's WAIC (using method 2 in Gelman *et al.* 2014, as recommended). The ddirmnom function in the extraDistr package (version 1.8.11; Wolodzko

2019) was used to compute log probability masses during WAIC calculations. A set of 23 models were subsequently fit by removing a single predictor from the initial model. WAICs were calculated for these models and compared to the initial model's WAIC. If the lowest WAIC in the new model set was less than or equal to the initial model's WAIC, then the model in the model set with the lowest WAIC was considered to be the new benchmark for best predictive accuracy. The process repeated by fitting another set of models with a single predictor removed from the benchmark model and comparing WAICs. When removing additional predictors failed to reduce or match WAIC, or after an intercept-only model was fit and no more predictors could be removed, then the variable selection process was finished and the model with the lowest WAIC of all the models fit was considered to have the best predictive accuracy. We considered the model with the best predictive accuracy to be the best-fit model. During backwards variable selection, the stratum dummy variables were considered to be a single predictor and were removed as a group.

The same Dirichlet-multinomial regression model and backwards variable selection approach were used to fit models to water and substrate read counts. Models for water samples were treated separately from models for substrate samples. For both water and substrate models, the non-stratum predictor matrix included site, a second-degree polynomial for week, and all interactions between these predictors. Centering and scaling of categorical and continuous predictors was performed in the same way as for the salamander models.

These same steps were performed for data from each of the 16S and ITS barcode regions, and for salamander samples, additional models were fit which substituted spatiotemporal covariates (*i.e.*, stratum, site, and time) with water quality predictors. The predictor matrix for the water quality models included a five-way interaction between age, life stage, temperature (°C), pH, and dissolved oxygen (ppm), all lower-level interactions, and the individual predictors. Conductivity (μS) was excluded from the analysis due to being highly colinear with pH (Pearson correlation coefficient of 0.817). We missed dissolved oxygen measurements (both ppm and percent) on August $11^{th}$ at Ponds Lake due to a faulty dissolved oxygen meter. For this analysis, we estimated dissolved oxygen (ppm) on this day by taking the average of the dissolved oxygen (ppm) measurements at each stratum for the weeks sampled immediately before and after at Ponds Lake. Quadratic terms were not introduced for the water quality models both because the Dirichlet-multinomial regression model is already quite flexible and because including such terms would have greatly increased the run-time required to perform backwards variable selection. Water quality predictors were mean-centered and scaled by one standard deviation prior to the creation of interaction terms. For the water quality models, parts of the Dirichlet-multinomial regression model associated with stratum were removed, and the predictor matrix was treated as the non-stratum predictor matrix.

The Dirichlet-multinomial regression models were fit in Stan (version 2.21.0; Carpenter *et al.* 2017) using the rstan R interface (version 2.21.2; Stan Development Team 2020) for Stan's Hamiltonian Monte Carlo (HMC) No-U-Turn sampler. Each model had four HMC chains run in parallel on four cores, and each chain had 500 warmup iterations and 500 sampling iterations without thinning. The average probability of accepting a posterior draw was set to 95%, and maximum tree depth was set to 20. When models were fit which excluded stratum predictors, which excluded all non-stratum predictors, or both, appropriate Stan models were used in which model parts associated with these excluded predictors were removed. The number of $\hat{R}$

convergence diagnostics in each model which were > 1.05 (disregarding the $\hat{R}$s of fixed parameters set to zero) were monitored, and for the final models selected by backwards variable selection, trace plots of the posteriors were used to assess convergence for parameters with $\hat{R}$s > 1.05. The vast majority of all models fit during the backwards variable selection process had zero $\hat{R}$s > 1.05, and many of these models had thousands of parameters each. The selected models for the water 16S region, salamander ITS region, and substrate ITS region each had at least one parameter with $\hat{R}$ > 1.05. The selected water 16S region model had 19 parameters with $\hat{R}$s > 1.05, and six parameters with $\hat{R}$s > 1.1 (all < 1.11). Examining the trace plots of parameters with $\hat{R}$s > 1.05 revealed that these parameters had some spurious draws which quickly returned towards the other chains, and otherwise convergence appeared fine. The selected salamander ITS region model had 46 parameters with $\hat{R}$s > 1.05, and two parameters with $\hat{R}$s > 1.1 (1.107 and 1.127). The trace plots of parameters with $\hat{R}$s > 1.05 appeared fine. The selected substrate ITS region model had 3 parameters with $\hat{R}$s > 1.05 (all < 1.08), and the trace plots of these parameters appeared fine. The selected water quality salamander models for both barcode regions had no parameters with $\hat{R}$s > 1.05.

#### *Microbial composition predictions*

Due to the softmax function and the compositional nature of the Dirichlet-multinomial regression model, the effect of regression coefficients on proportional abundances is difficult to interpret. While taxa with the same regression coefficients within a predictor experience the same percent change in proportional abundances with changes in that predictor, resulting in the same shape of relationship, additional interpretation is limited since the effect of one regression coefficient on proportional abundances is dependent on the values of all other regression coefficients in the model. The modeled proportional abundances do not change linearly with changes in predictor values, and changes in predictor values elsewhere in the model can change whether proportional abundances are increasing or decreasing at a given value of another predictor. Furthermore, the regression coefficients do not represent changes in underlying absolute abundances since a value of zero represents the regression coefficient values of the reference taxon, the selection of which is arbitrary.

Instead of attempting a direct interpretation of the Dirichlet-multinomial regression coefficients, we opted for a graphical interpretation of the best-fit models using predicted proportional abundances. For the spatiotemporal salamander, water, and substrate models, we generated posterior proportional abundance predictions for each combination of non-stratum predictors observed in the datasets. For water, proportional abundance predictions were also made for July 14$^{th}$ at Gibson Lakes, a sampling event for which water samples had to be discarded. Non-stratum predictor matrices were prepared as in the compositional modeling section with the following differences. Mean centering and standard deviation scaling were performed using the same mean and standard deviation values used to center and scale the models' original predictors, and non-stratum predictors were subsetted to just those included in the best-fit models. If stratum was included as a predictor in the best-fit model, then a stratum predictor matrix was prepared as described above, and all dummy variables were set to zero prior to centering with the same mean values used to center the model's original stratum predictors. Due to the sum-to-zero constraint on all stratum regression coefficients within a taxon, the average effect of stratum for a taxon is zero, and setting stratum predictor values to either all zero or all one prior to centering results in predictions for the average stratum. This is analogous to leaving out random effects from frequentist models to achieve group-level predictions. These predictions represent the posterior distributions of taxa proportional abundances.

#### *Salamander length-weight regression*

In order to model microbial absolute abundance in terms of density, and because we did not measure swabbed area in the field, we derived estimates proportional to swabbed area based on a regression between salamander length and weight. Snout-vent length (SVL) can be related to weight via a power function (Hile 1936) which takes the form:

$$\text{Weight (g)} = a * \text{SVL (mm)}^{b}$$

Where $a$ and $b$ are constants. If $b$ = 3, then the organism exhibits isometric growth where the body grows proportionally, and the organism exhibits allometric growth otherwise. The relationship can be linearized by taking the logarithm of both sides:

$$\log(\text{Weight (g)}) = \log(a) + b * \log(\text{SVL (mm)})$$

Fitting the linearized model between salamander SVL and weight while including an interaction term between SVL and site yielded an insignificant interaction p-value of 0.166. Since there was insufficient evidence to suggest a unique length-weight relationship for each site, we fit a common length-weight relationship for both sites by excluding the interaction term, and we performed a t-test to determine whether $b$ was significantly different than 3 using the hoCoef function in the FSA package (version 0.8.30; Ogle *et al.* 2020). A t-test p-value of $< 0.001$ suggested that $b$ differed significantly from 3 and that salamander growth was allometric.

#### *Estimating swabbed area*

To express microbial absolute abundance in terms of density, we derived a proportional relationship between salamander length and swabbed area based on our earlier length-weight regression (see *Salamander length-weight regression* section above). Under isometric growth, area is proportional to length squared. However, our length-weight regression suggested allometric growth in which a salamander's body does not grow proportionally in all dimensions.

Our earlier length-weight model yielded the following relationship:

$$\text{Weight (g)} = 0.000669 * \text{SVL (mm)}^{2.350}$$

$\text{SVL (mm)}^{b}$ is proportional to the volume of the salamander, with $\text{length} * \text{width} * \text{height}$ being proportional to $\text{length}^{3}$ if growth is isometric. We can decompose the exponent term into three dimensions as:

$$\text{Length}^{2.350} \propto \text{Length} * \text{Width} * \text{Height}$$

$$\text{Length}^{1.350} \propto \text{Width} * \text{Height}$$

We make the assumption that salamander width and height grow proportionally:

$$\text{Width} \propto \text{Height}$$

This assumption simplifies $\text{Length}^{1.350} \propto \text{Width} * \text{Height}$ to:

$$D^2 \propto \text{Length}^{1.350}$$
$$D \propto \text{Length}^{0.675}$$
Where D is width or height.

Assuming that swabbed area is proportional to salamander length times width, we can derive a proportional relationship between length and swabbed area:
$$\text{Area} \propto \text{Length} * \text{Width}$$
$$\text{Area} \propto \text{Length} * \text{Length}^{0.675}$$
$$\text{Area} \propto \text{Length}^{1.675}$$

#### *Microbial absolute abundance modeling*

Since the swabbed area (the belly) differs between salamanders of different sizes, we express microbial absolute abundance in terms of density (count per unit area). While we did not measure swabbed area in the field, we derived estimates proportional to swabbed area based on a salamander length-weight regression and the assumption that salamander width and height grow proportionally (see *Estimating swabbed area* section above). To model the density of microbes on salamander skin, we used a Bayesian negative binomial LASSO model for each taxon. For each barcode region, we fit models to the same 100 microbial taxa included in the composition modeling, plus the "other" category. In order to incorporate many predictors into the model without overfitting, and because we had a group of categorical predictors (*i.e.*, stratum), we borrowed from the Bayesian LASSO (Park & Casella 2008) and Bayesian group LASSO (Xu & Ghosh 2015) in formulating our model. Specifically, our model takes the form:

Taxon read counts $c_i$ are distributed according to the negative binomial distribution with probability parameter $p_i$ and size parameter $r$, which controls the degree of overdispersion relative to the Poisson distribution.
$$c_i \sim \text{negative binomial}(p_i, r)$$

The probability parameter $p_i$ is derived from the size parameter $r$ and the expected value of the negative binomial distribution $\mu_i$.
$$p_i = \frac{r}{r + \mu_i}$$

As parameterized in JAGS (version 4.3.0; Plummer 2003), the negative binomial distribution has:
$$\mathbb{E}[c] = \frac{r * (1 - p)}{p} = \mu$$
and
$$\text{Var}[c] = \frac{(1 - p) * r}{p^2} = \mu + \frac{\mu^2}{r}$$

The expected value of the negative binomial distribution $\mu_i$ is the product of taxon density (arbitrary units), synthgene count, and a value proportional to swabbed area (estimated as SVL (mm)$^{1.675}$).
$$\mu_i = \text{density}_i * \text{synthgene}_i * \text{area}_i$$

Taxon density is related to a linear predictor combination via a log link.

$$\log(\text{density}_i) = \beta_0 + B_{\text{Stratum}} * X_{\text{Stratum}_i}{}^{\mathsf{T}} + B_{\text{Non-stratum}} * X_{\text{Non-stratum}_i}{}^{\mathsf{T}}$$
$$B_{\text{Stratum}} = \left[\beta_{\text{Stratum}_1} \cdots \beta_{\text{Stratum}_S}\right]$$
$$B_{\text{Non-stratum}} = \left[\beta_{\text{Non-stratum}_1} \cdots \beta_{\text{Non-stratum}_Q}\right]$$
$$X_{\text{Stratum}_i} = \left[x_{\text{Stratum}_{1_i}} \cdots x_{\text{Stratum}_{S_i}}\right]$$
$$X_{\text{Non-stratum}_i} = \left[x_{\text{Non-stratum}_{1_i}} \cdots x_{\text{Non-stratum}_{Q_i}}\right]$$

A weakly informative prior is used on the intercept term $\beta_0$.

$$\beta_0 \sim \text{normal}(0, 10^6)$$

Following the Bayesian LASSO and Bayesian group LASSO, the priors for regression coefficients are a scale mixture of normals. Following other applications of the Bayesian LASSO to generalized linear models, we do not condition the regression coefficient priors on the residual variance $\sigma^2$ (*e.g.*, Huang & Cai 2013; Tang *et al.* 2017), as is the case for linear models. Following the Bayesian group LASSO, a gamma mixing density is used for grouped predictors (*i.e.*, stratum). $\lambda$ serves as the model's overall shrinkage parameter, and $\tau_{\text{Stratum}}$ serves as a stratum-specific shrinkage parameter. A Bernoulli-distributed binary inclusion variable $\text{IV}_{\text{Stratum}}$ allows for exact zero estimates.

$$\beta_{\text{Stratum}_{s \neq S}} \sim \text{normal}(0, \tau_{\text{Stratum}}{}^2) * \text{IV}_{\text{Stratum}}$$
$$\tau_{\text{Stratum}}{}^2 \sim \text{gamma}\left(\frac{S}{2}, \frac{\lambda^2}{2}\right)$$
$$\text{IV}_{\text{Stratum}} \sim \text{Bernoulli}(\text{IP})$$

Since we view stratum as a random effect, we center the stratum effects around zero by applying a sum-to-zero constraint on the last stratum regression coefficient.

$$\beta_{\text{Stratum}_S} = -1 * \sum_{s=1}^{S-1} \beta_{\text{Stratum}_s}$$

When there is no grouping structure, the gamma mixing density of the Bayesian group LASSO simplifies to the exponential mixing density of the Bayesian LASSO (Van Erp *et al.* 2019). The $\tau_{\text{Stratum}_q}$ parameters again serve as predictor-specific shrinkage parameters, and Bernoulli-distributed binary inclusion variables $\text{IV}_{\text{Non-stratum}_q}$ allow for exact zero estimates.

$$\beta_{\text{Non-stratum}_q} \sim \text{normal}\left(0, \tau_{\text{Non-stratum}_q}{}^2\right) * \text{IV}_{\text{Non-stratum}_q}$$
$$\tau_{\text{Non-stratum}_q}{}^2 \sim \text{exponential}\left(\frac{\lambda^2}{2}\right)$$
$$\text{IV}_{\text{Non-stratum}_q} \sim \text{Bernoulli}(\text{IP})$$

We provide weakly informative gamma priors for the size parameter $r$ and the squared overall shrinkage parameter $\lambda^2$.

$$r \sim \text{gamma}(0.01, 0.01)$$
$$\lambda^2 \sim \text{gamma}(0.01, 0.01)$$

A beta prior is provided for inclusion probability IP with shape parameters 0.02 and 1.98. With these shape parameters, the prior expectation is to select 1% of the predictors.

$$\text{IP} \sim \text{beta}(0.02, 1.98)$$

Where $c_i$ is the taxon read count,
$i = 1 \dots N$ is the sample number,
$s = 1 \dots S$ is the stratum number,
$x_{\text{Stratum}_{s_i}}$ is the stratum predictor value,
$q = 1 \dots Q$ is the non-stratum predictor number,
and $x_{\text{Non-stratum}_{q_i}}$ is the non-stratum predictor value.

We prepared our predictor matrices in the same manner as for the Dirichlet-multinomial regression model except for the following differences. To achieve similar scales for all our input variables, as desired for the Bayesian LASSO, we scaled continuous predictor values by two standard deviations (prior to the creation of polynomial and interaction terms) to put them on a similar scale to the categorical dummy variables (Gelman 2008). We also chose to include second- the fourth-degree polynomial terms for age and week, respectively. Our resulting non-stratum predictor matrix included four-way interactions between a second-degree polynomial for age, life stage, site, and a fourth-degree polynomial for week, all lower-level interactions, and the individual predictors. We did not add one to each of the read counts, as was done in the microbial composition modeling.

The negative binomial LASSO models were fit in JAGS (version 4.3.0; Plummer 2003) using the rjags R interface (version 4.10; Plummer 2019) and GNU parallel (version 20201022; Tange 2011). Three Markov chain Monte Carlo (MCMC) chains were run with 50,000 adaptation, 50,000 warmup, and 500,000 sampling iterations with a thinning interval of 2. Gelman-Rubin convergence diagnostics were used to assess model convergence.

#### *Microbial absolute abundance predictions*

We generated posterior predictions of microbial densities for each combination of non-stratum predictors observed in the salamander datasets, where predictions were for the average stratum. For generating absolute abundance predictions, non-stratum predictor matrices were prepared as described in the absolute abundance modeling section with the following differences. Mean centering and two standard deviation scaling were performed using the same mean and standard deviation values used to center and scale the models' original predictors. A stratum predictor matrix was prepared as described above, and all dummy variables were set to zero prior to centering with the same mean values used to center the model's original stratum predictors.

The posterior distributions for taxa densities were summarized with 95% credible intervals, 50% credible intervals, and their median values. To estimate the antifungal function of microbial communities in terms of absolute abundances, we summed the posterior density predictions of all modeled bacterial taxa belonging to each *Bd*-inhibition category, with "other" bacterial taxa belonging to none of these categories. We summarized the densities of each *Bd*-inhibition category with 95% credible intervals, 50% credible intervals, and their median values.

### Results

#### *Field sampling*

Based on observed sizes of males and females, most age-2+ salamanders are thought to have been adults. 32 of 55 (58.2%) of sexed age-2+ salamanders were female (64% for Gibson Lakes and 53.3% for Ponds Lake). Males develop swollen cloacas once sexually mature, and only one non-male age-2+ salamander (83 mm; assumed to be female) had an SVL less than the smallest male (84 mm), with other small salamanders in the range of 85 to 87 mm SVL being a mix of males (3) and females (4). Given the overlap in size between males and females, few subadults are expected to have been included in the age-2+ age class since male salamanders of this size were showing clear signs of sexual maturity.

#### *Microbial diversity across sample types*

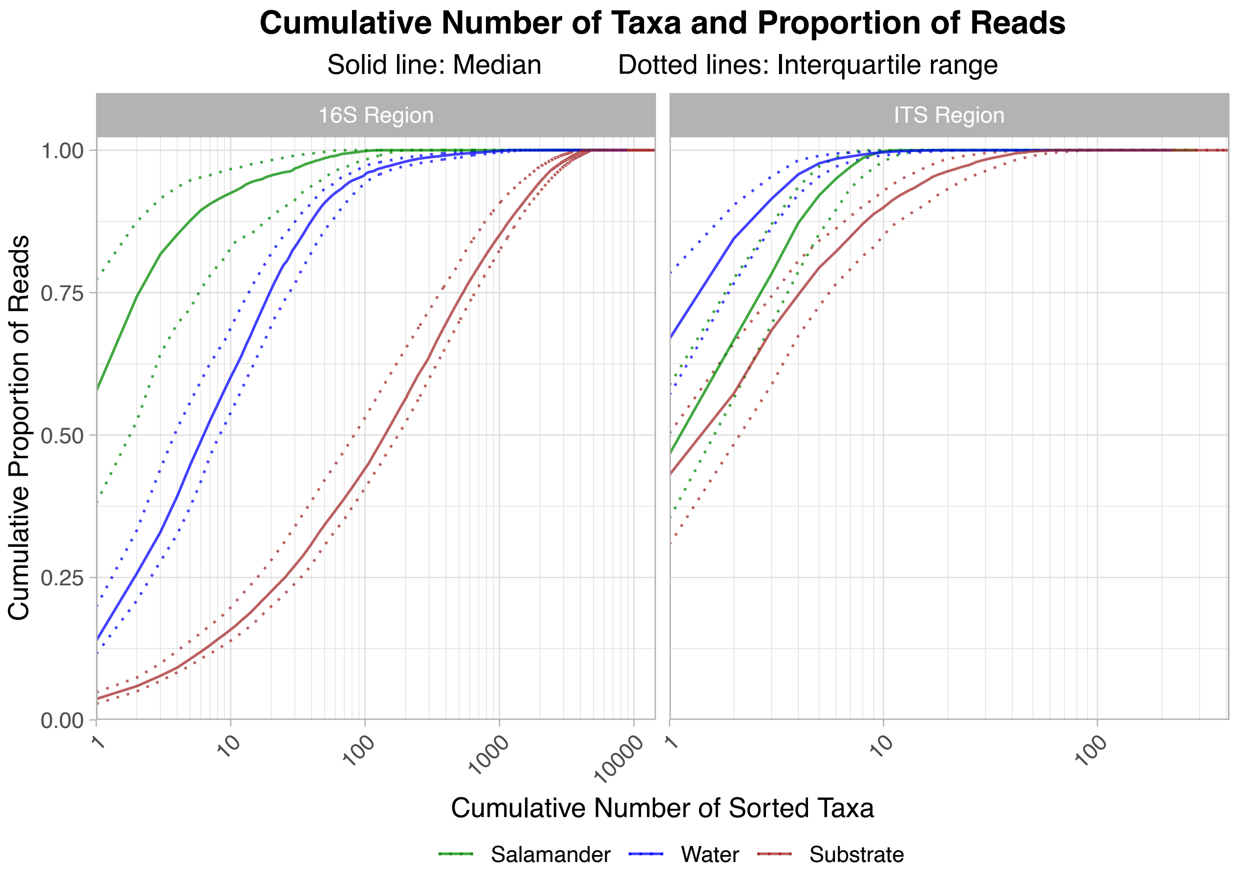

Figure S7. Cumulative proportion of sample reads by cumulative number of sorted taxa for each field sample type and barcode region. For each sample, taxa were sorted by descending read count. Solid lines represent sample medians, and dotted lines represent sample interquartile ranges. Note the $log_{10}$ scale on the x-axis.

##### *Taxonomic resolution*

Most 16S reads for most salamander samples were classified to down to at least family (Figure S8), about a quarter of 16S reads within each sample were classified down to at least genus, and a small proportion of 16S reads within each sample were classified down to species. The proportion of reads for each salamander sample which were classified down to at least a given taxonomic level was more variable for the ITS region, and the boxplots of these ITS proportions are much more uniform across taxonomic levels than for the 16S reads. Our classifier was able to classify most 16S reads within each salamander sample down to genus, after which the proportion of reads classified dropped sharply. Compared to the 16S reads, our classifier classified a lower proportion of ITS reads within each salamander sample down to any given taxonomic level above genus, but the proportion of ITS reads classified remained very similar across all taxonomic levels. Within barcode regions, differences in the proportions of reads classified down to at least a given taxonomic level within each salamander sample were minor between life stages and sites, except for ITS reads from metamorphosed individuals at Ponds Lake, which tended to be higher as *Bd* is prevalent in these samples (Figure S17).

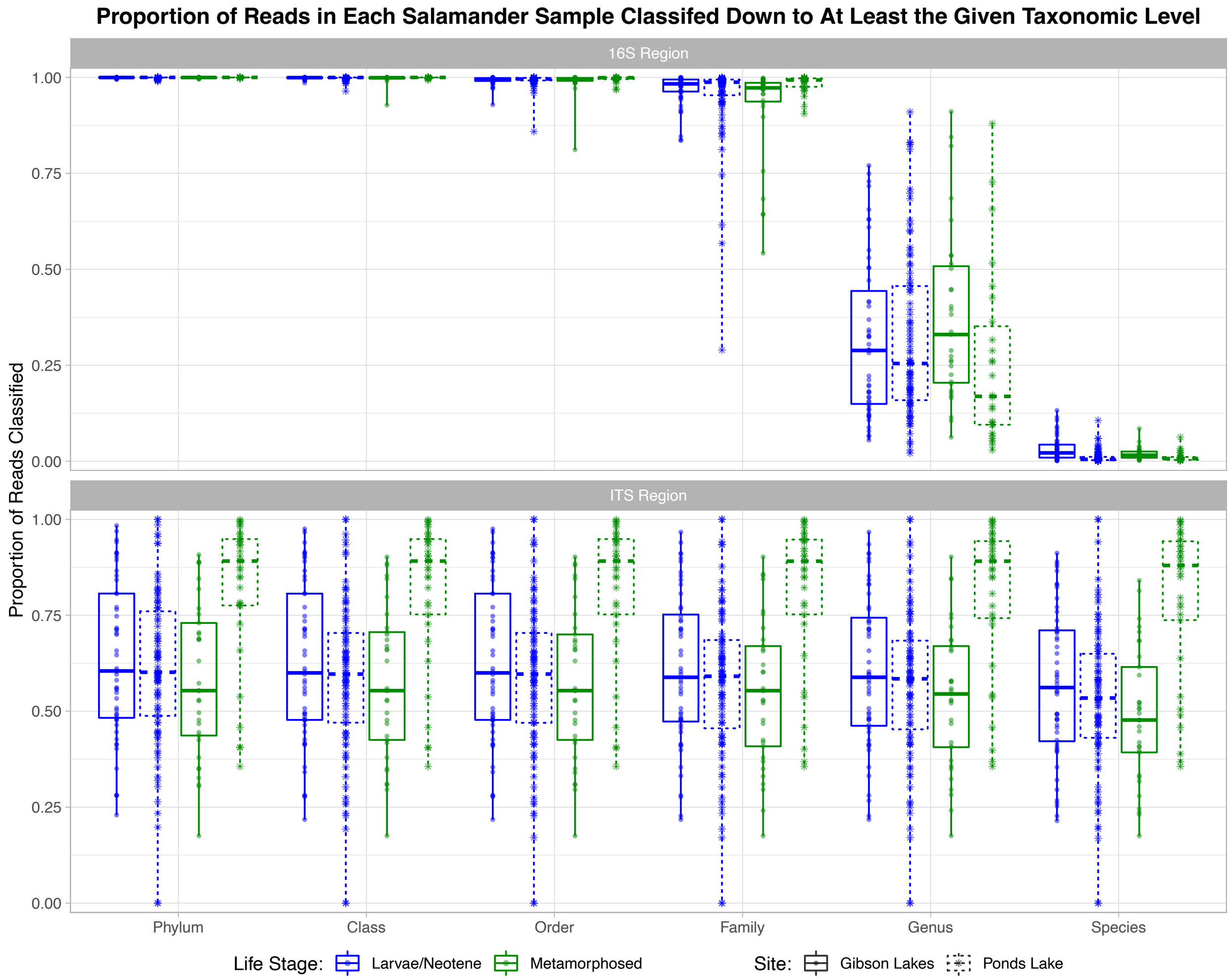

Figure S8. Proportion of salamander sample reads classified down to at least a given taxonomic level. Each point at a given taxonomic level represents a microbiome sample.

#### ***Antifungal prediction***

Table S1. Predicted *Batrachochytrium dendrobatidis* (*Bd*)-inhibition statuses of the top 100 bacterial taxa. Top 100 bacterial taxa not listed here have an uncertain *Bd*-inhibition status.

| Predicted *Bd*-Inhibition Status | Taxon | Probability of *Bd*-Inhibition |
|---|---|---|
| Inhibitory | Aeromonas 1 | 99.9% |
| | Brevibacterium 1 | 99.9% |
| | Comamonadaceae 5 | 99.8% |
| | Hafnia-Obesumbacterium 1 | 99.8% |
| | Paucibacter 2 | 99.9% |
| | Pseudarthrobacter 1 | 99.9% |
| | Pseudomonas 1 | 99.9% |
| | Pseudomonas 3 | 99.9% |
| | Pseudomonas 5 | 99.9% |
| | Staphylococcus 2 | 99.9% |
| | Stenotrophomonas 1 | 99.9% |
| Non-Inhibitory | Chryseobacterium 1 | 0.1% |
| | Enterobacterales 1 | 0.1% |
| | Flavobacterium 2 | 0.2% |
| | Yersinia 1 | 0.2% |

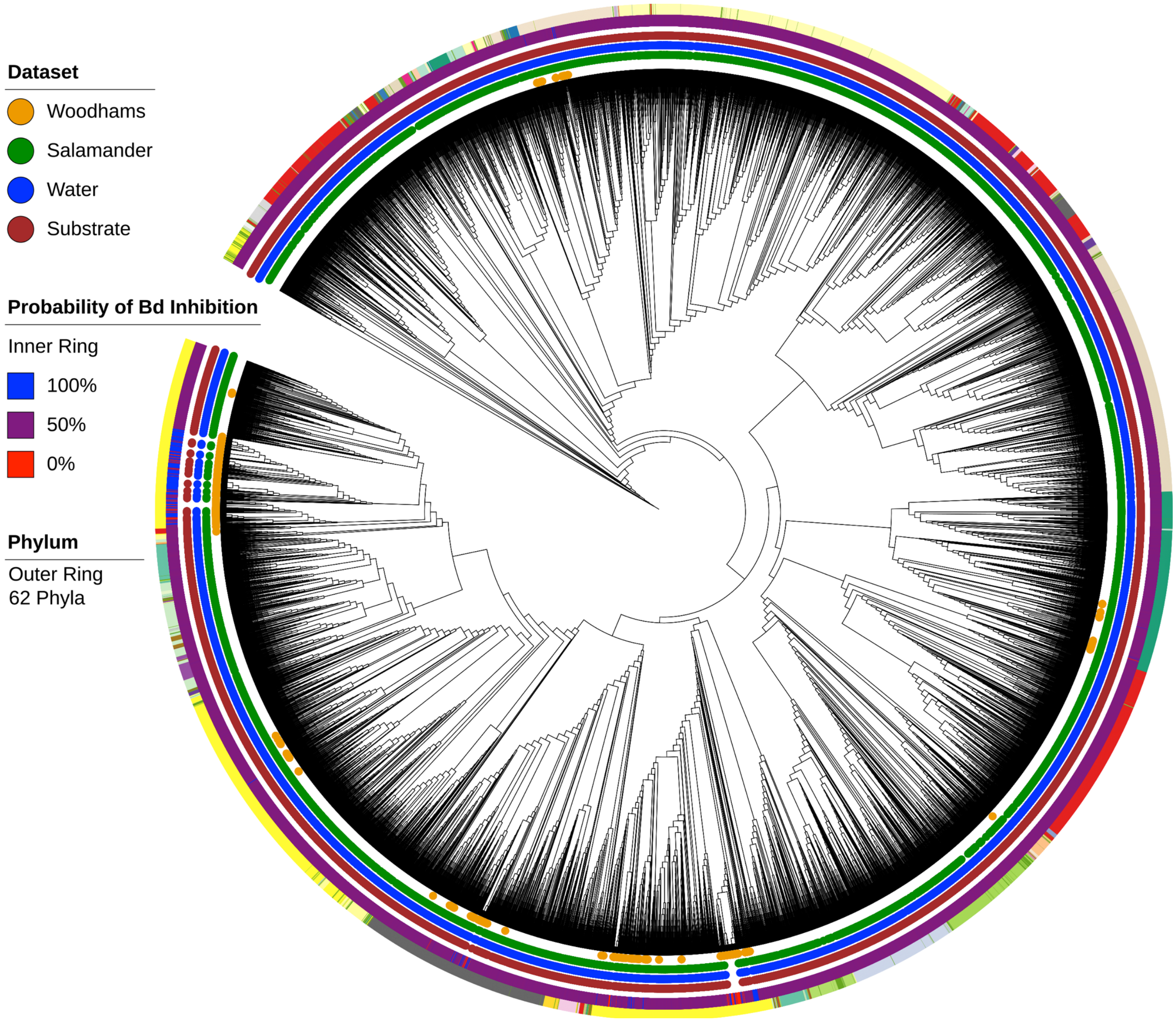

Figure S9. A phylogenetic tree of bacterial taxa observed in field samples and the Woodhams database. Displayed with the tree is which sample type or dataset each taxon was observed in, each taxon's posterior probability of being *Batrachochytrium dendrobatidis* (*Bd*)-inhibitory, and each taxon's phylum. Phyla for the Woodhams taxa are from the UCLUST taxonomies provided in the database (Bacteroidetes and Actinobacteria were changed to their synonyms Bacteroidota and Actinobacteriota to match the taxonomy of our Silva reference database). As there are many phyla, some have similar colors, and one phylum color represents taxa whose phyla are unknown.

#### *Water quality*

The field season finished on September 29$^{th}$ when Gibson Lakes became too cold to safely sample salamanders. Water quality and environmental microbiomes were still taken at Gibson Lakes on September 29$^{th}$ despite not catching any salamanders. The water level at Gibson Lakes on September 29$^{th}$ was too low to measure relative lake elevation.

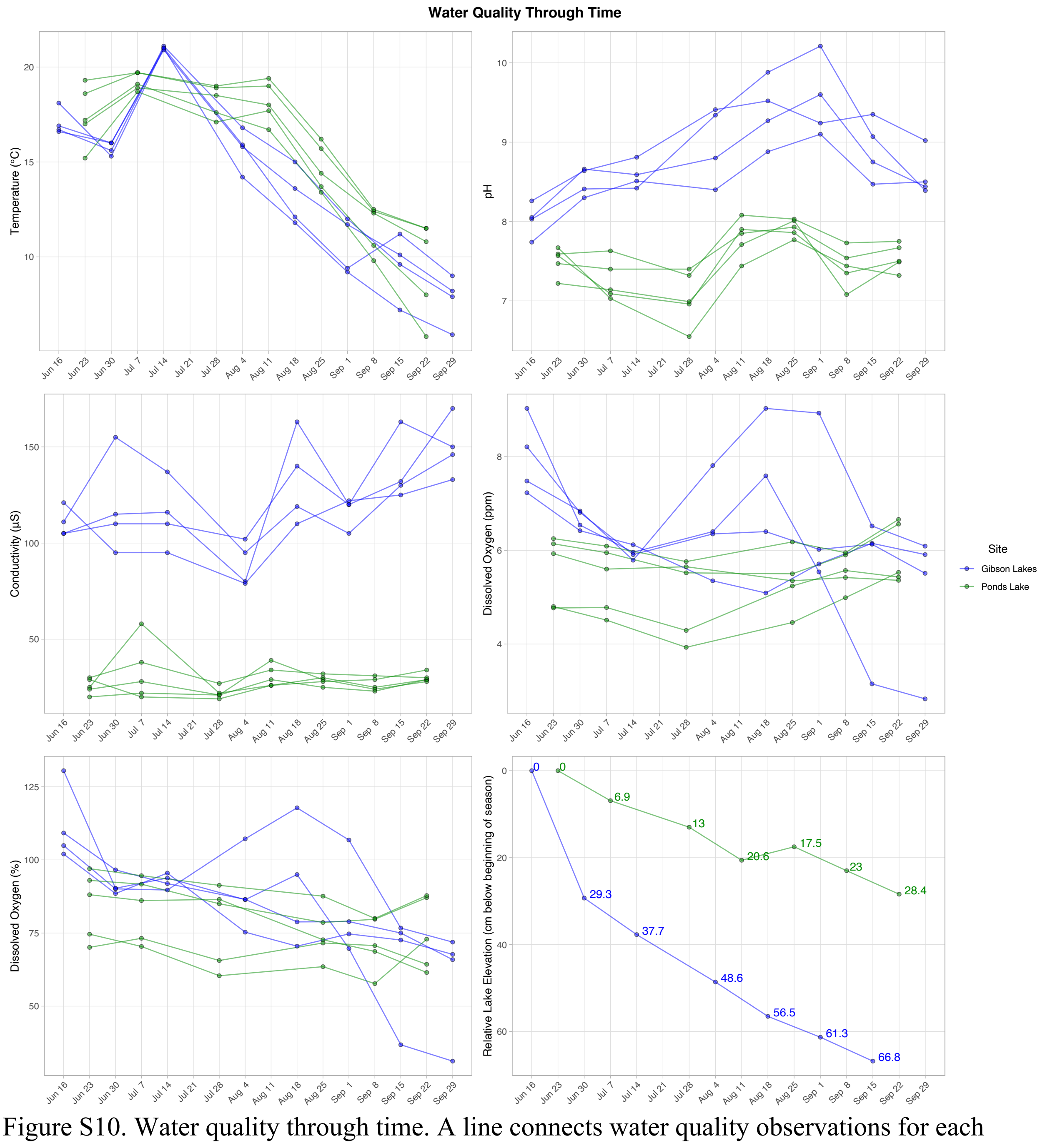

Figure S10. Water quality through time. A line connects water quality observations for each stratum.

#### *16S proportional abundance plots*

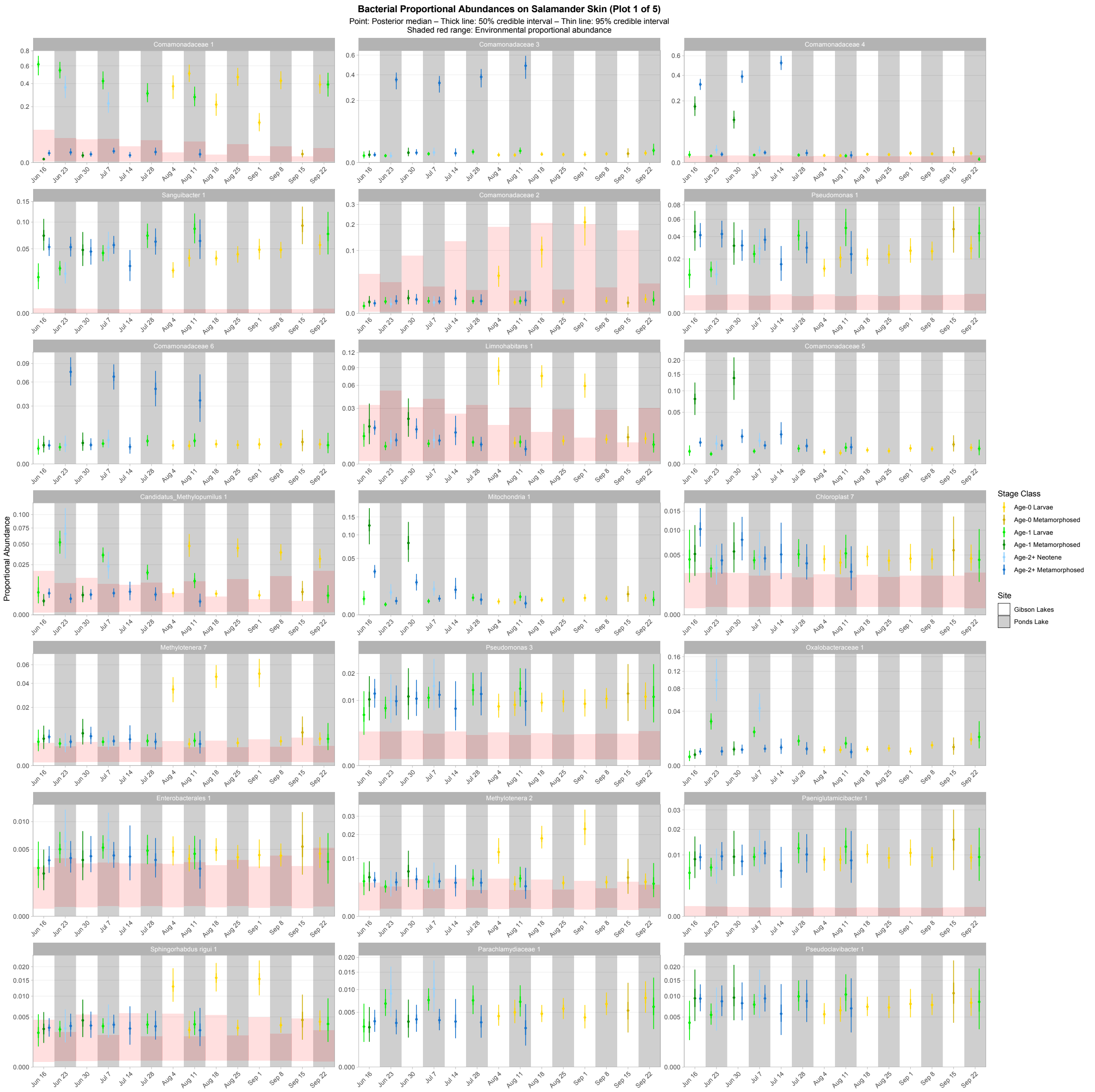

Figure S11. First plot of proportional abundance predictions for the top 100 bacterial taxa. Points, thick lines, and thin lines represent posterior medians, 50% credible intervals, and 95% credible intervals, respectively. Shaded red ranges represent environmental proportional abundances. Taxa without shaded red ranges were not detected in either water or substrate. Note the square root scale on the y-axis. Taxa are ordered (left to right, top to bottom) by descending average proportion of reads in salamander samples in which each combination of site and life stage receives equal weight.

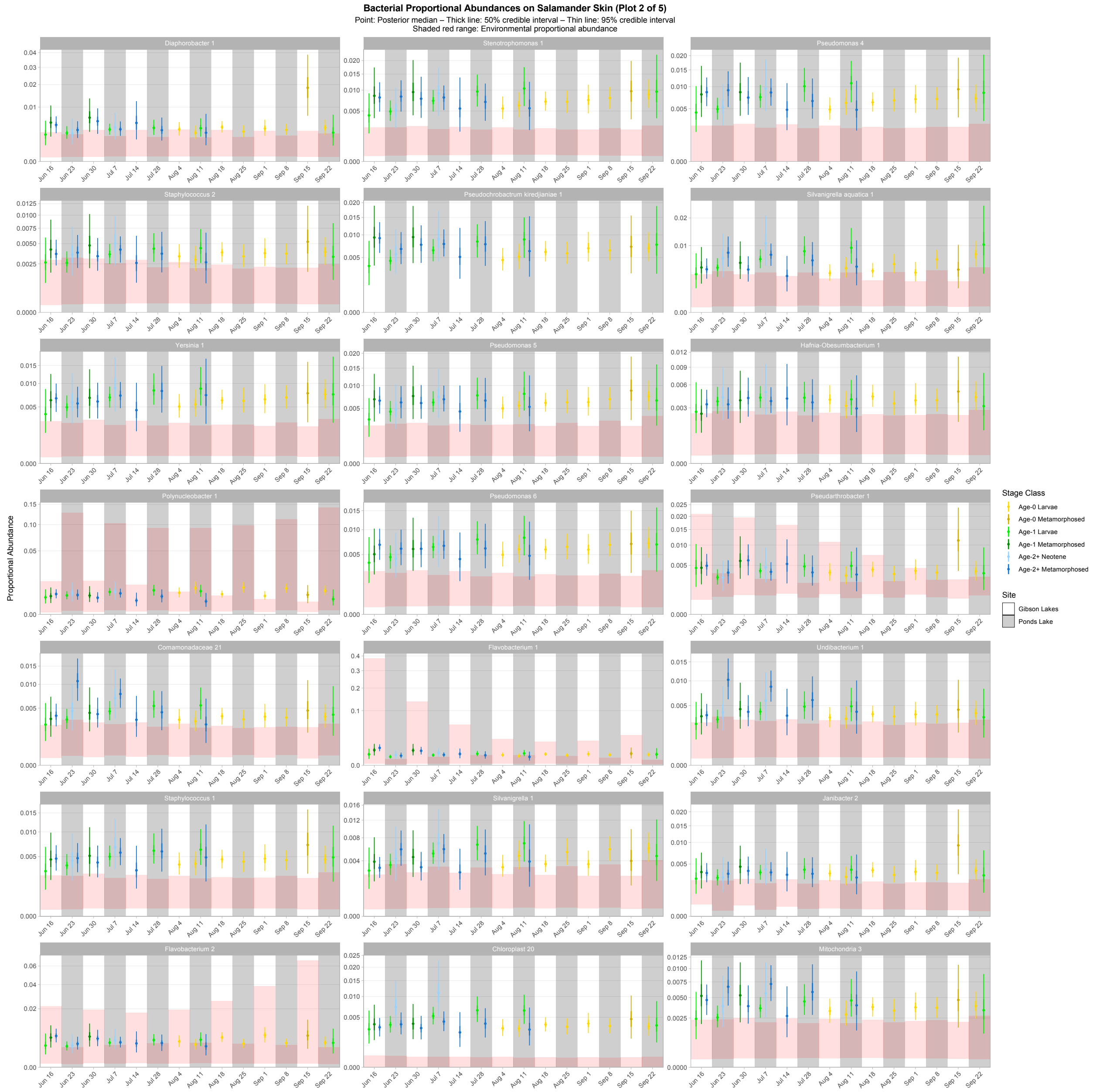

Figure S12. Second plot of proportional abundance predictions for the top 100 bacterial taxa. Points, thick lines, and thin lines represent posterior medians, 50% credible intervals, and 95% credible intervals, respectively. Shaded red ranges represent environmental proportional abundances. Taxa without shaded red ranges were not detected in either water or substrate. Note the square root scale on the y-axis. Taxa are ordered (left to right, top to bottom) by descending average proportion of reads in salamander samples in which each combination of site and life stage receives equal weight.

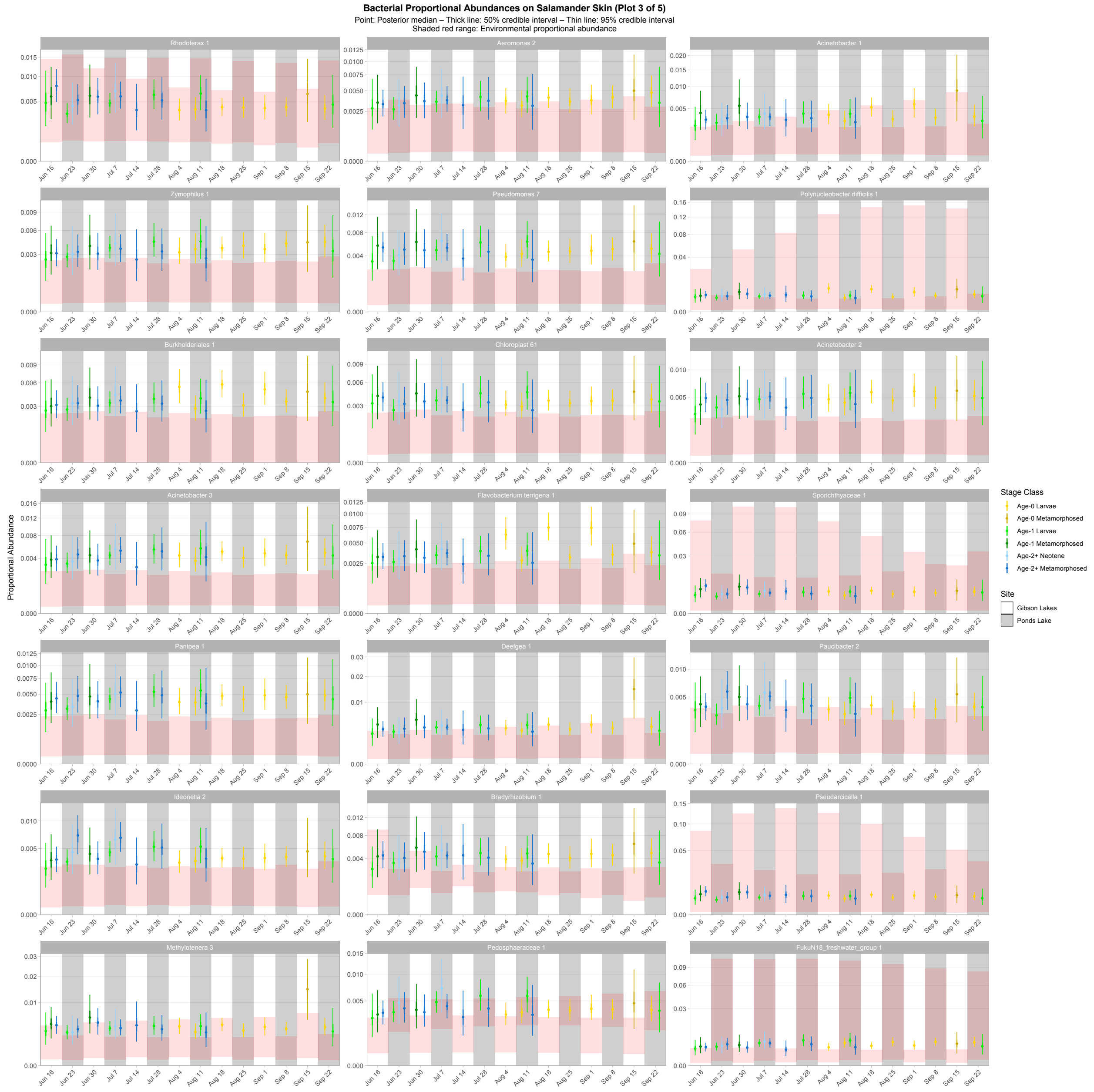

Figure S13. Third plot of proportional abundance predictions for the top 100 bacterial taxa. Points, thick lines, and thin lines represent posterior medians, 50% credible intervals, and 95% credible intervals, respectively. Shaded red ranges represent environmental proportional abundances. Taxa without shaded red ranges were not detected in either water or substrate. Note the square root scale on the y-axis. Taxa are ordered (left to right, top to bottom) by descending average proportion of reads in salamander samples in which each combination of site and life stage receives equal weight.

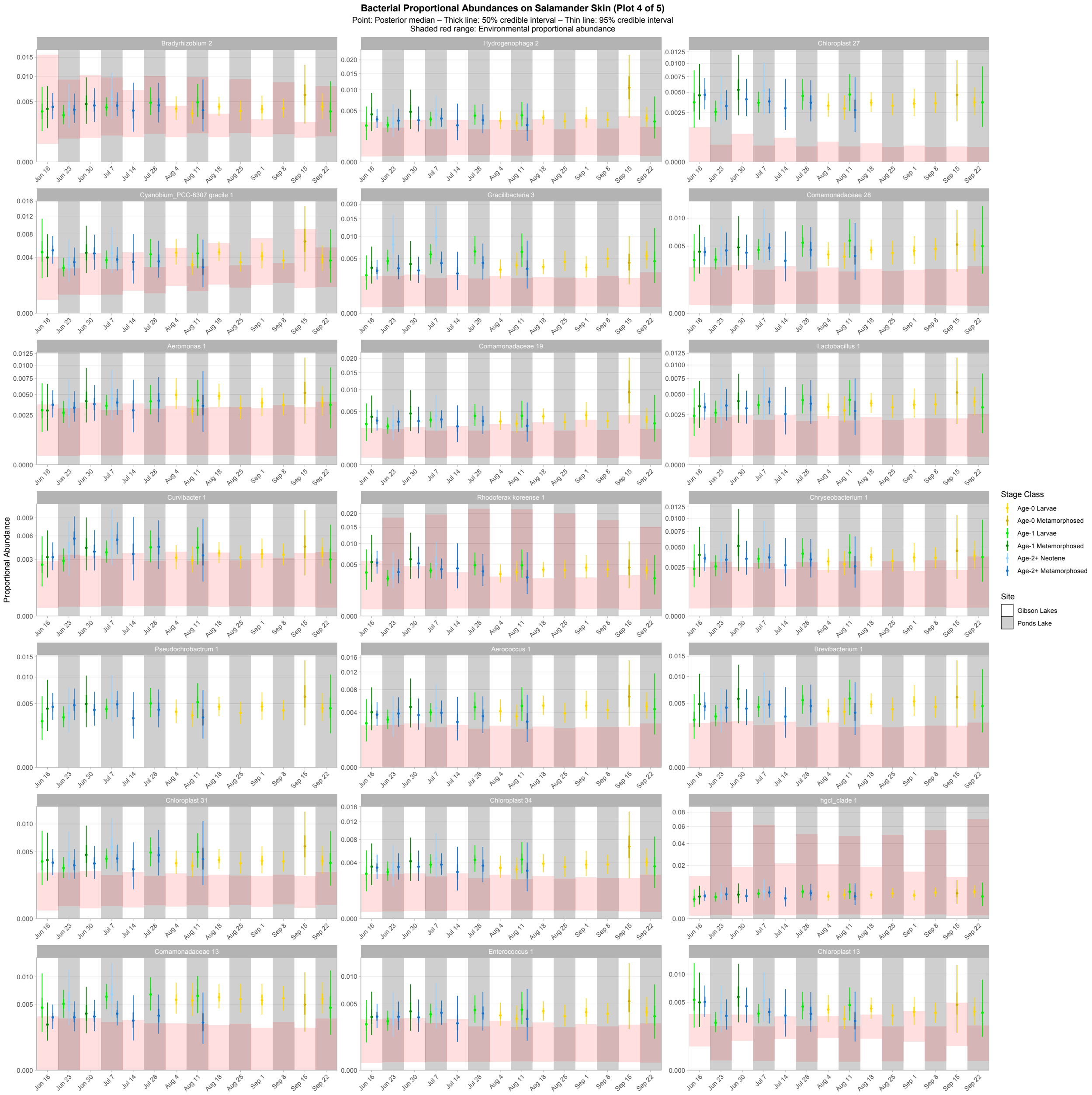

Figure S14. Fourth plot of proportional abundance predictions for the top 100 bacterial taxa. Points, thick lines, and thin lines represent posterior medians, 50% credible intervals, and 95% credible intervals, respectively. Shaded red ranges represent environmental proportional abundances. Taxa without shaded red ranges were not detected in either water or substrate. Note the square root scale on the y-axis. Taxa are ordered (left to right, top to bottom) by descending average proportion of reads in salamander samples in which each combination of site and life stage receives equal weight.

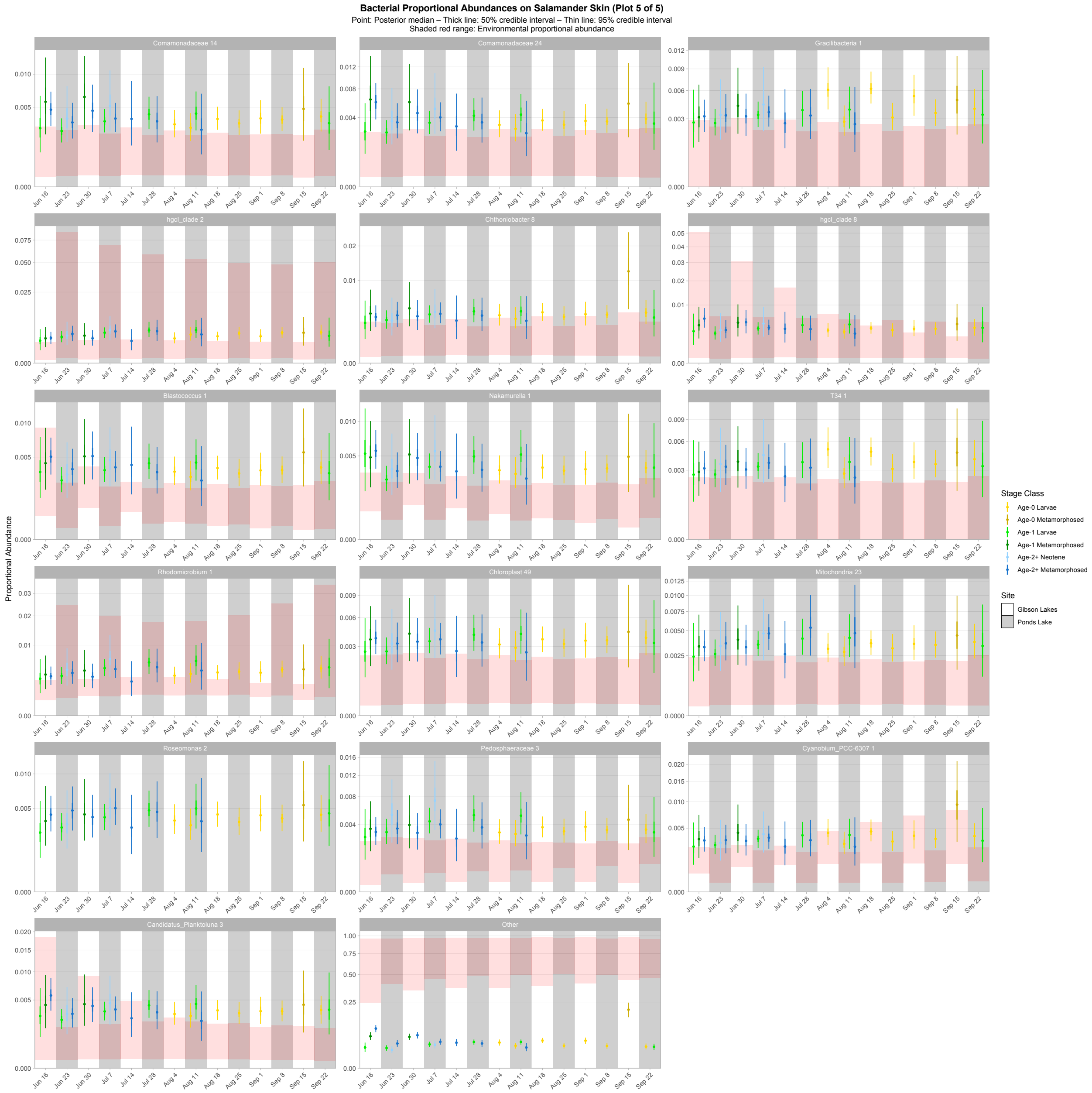

Figure S15. Fifth plot of proportional abundance predictions for the top 100 bacterial taxa. Points, thick lines, and thin lines represent posterior medians, 50% credible intervals, and 95% credible intervals, respectively. Shaded red ranges represent environmental proportional abundances. Taxa without shaded red ranges were not detected in either water or substrate. Note the square root scale on the y-axis. Taxa are ordered (left to right, top to bottom) by descending average proportion of reads in salamander samples in which each combination of site and life stage receives equal weight.

#### *16S Bray-Curtis dissimilarity*

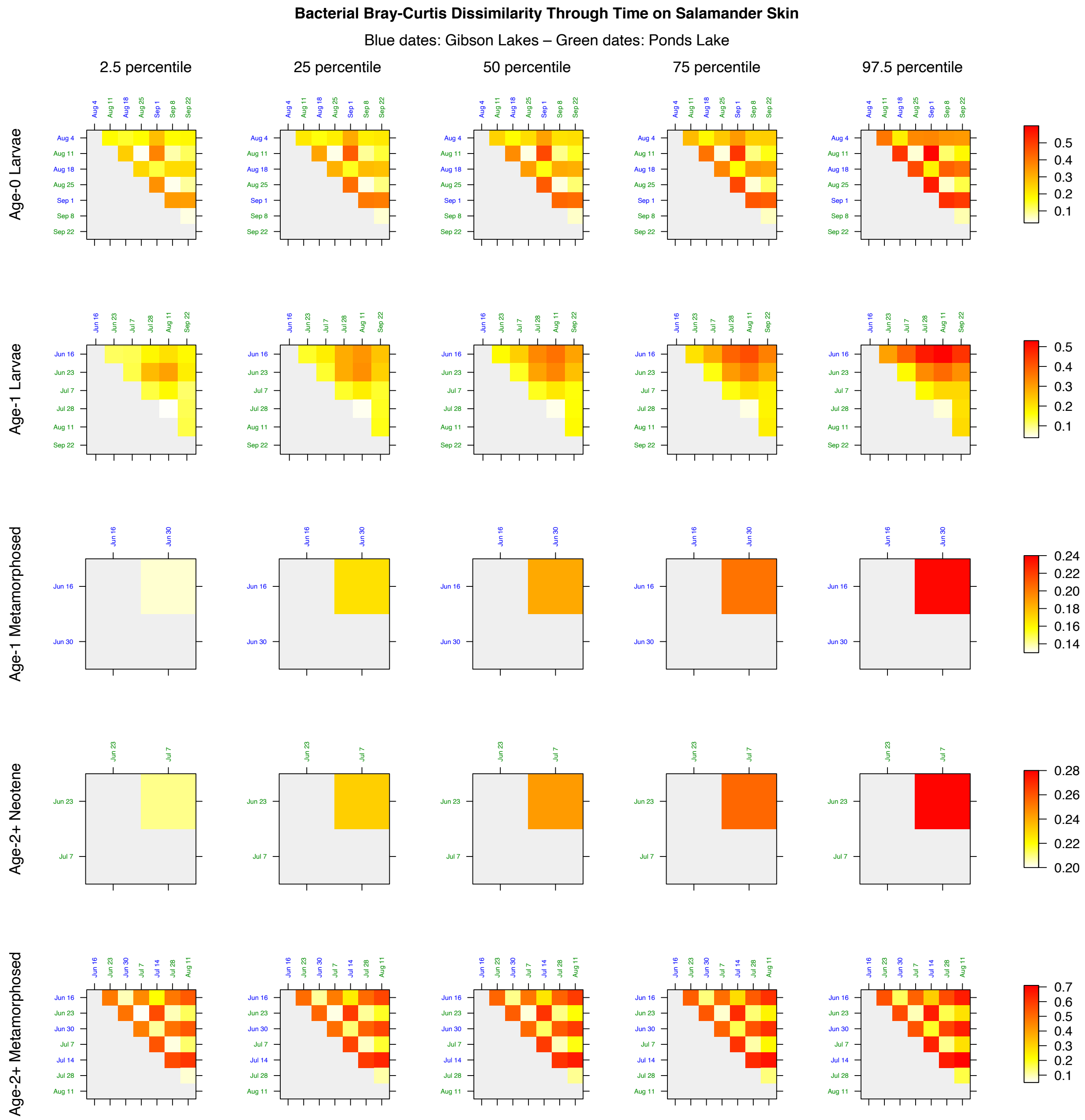

Figure S16. Bray-Curtis dissimilarity through time for bacterial communities on salamander skin.

### *ITS proportional abundance plots*

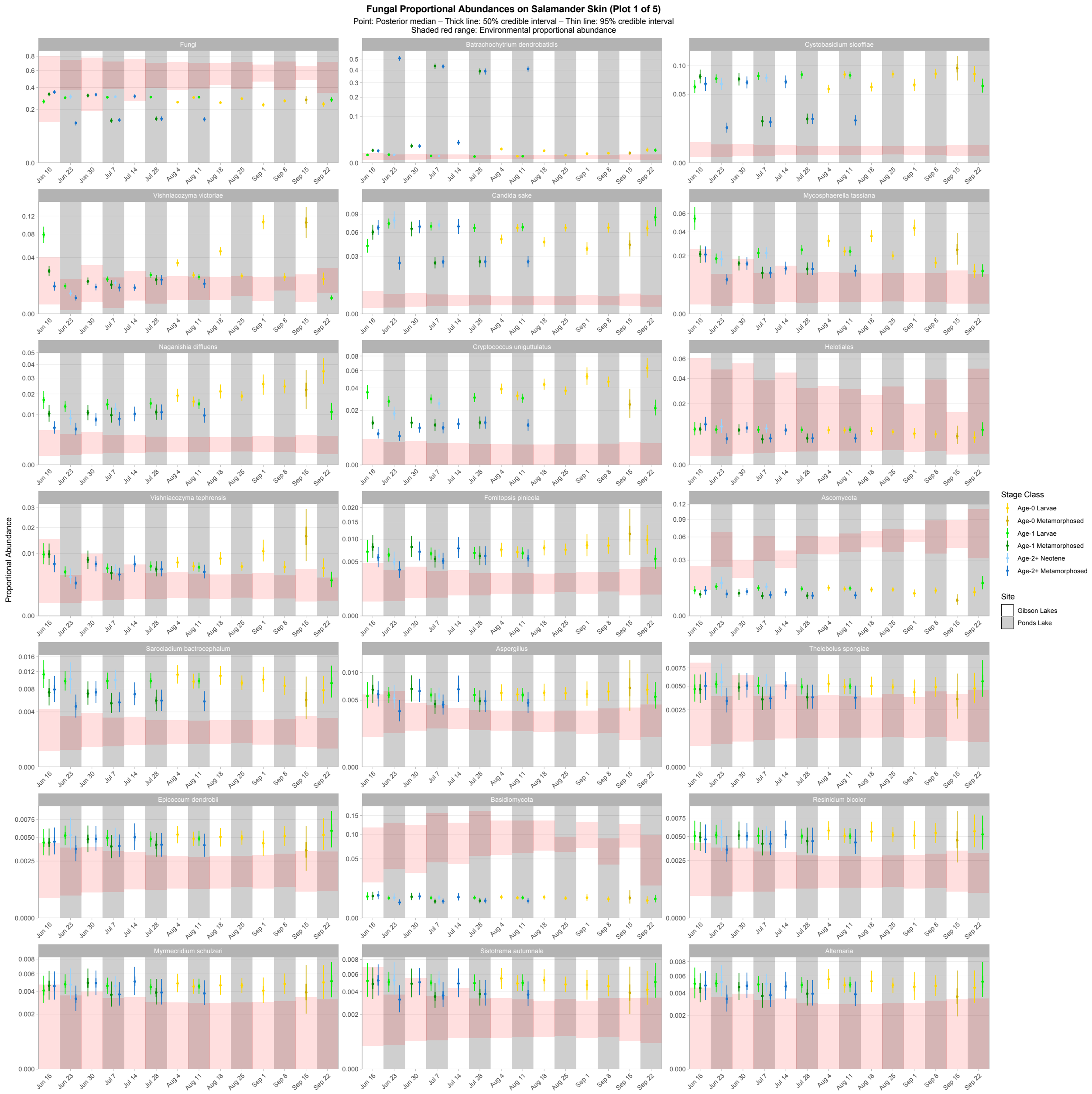

Figure S17. First plot of proportional abundance predictions for the top 100 fungal taxa. Points, thick lines, and thin lines represent posterior medians, 50% credible intervals, and 95% credible intervals, respectively. Shaded red ranges represent environmental proportional abundances. Taxa without shaded red ranges were not detected in either water or substrate. Note the square root scale on the y-axis. Taxa are ordered (left to right, top to bottom) by descending average proportion of reads in salamander samples in which each combination of site and life stage receives equal weight.

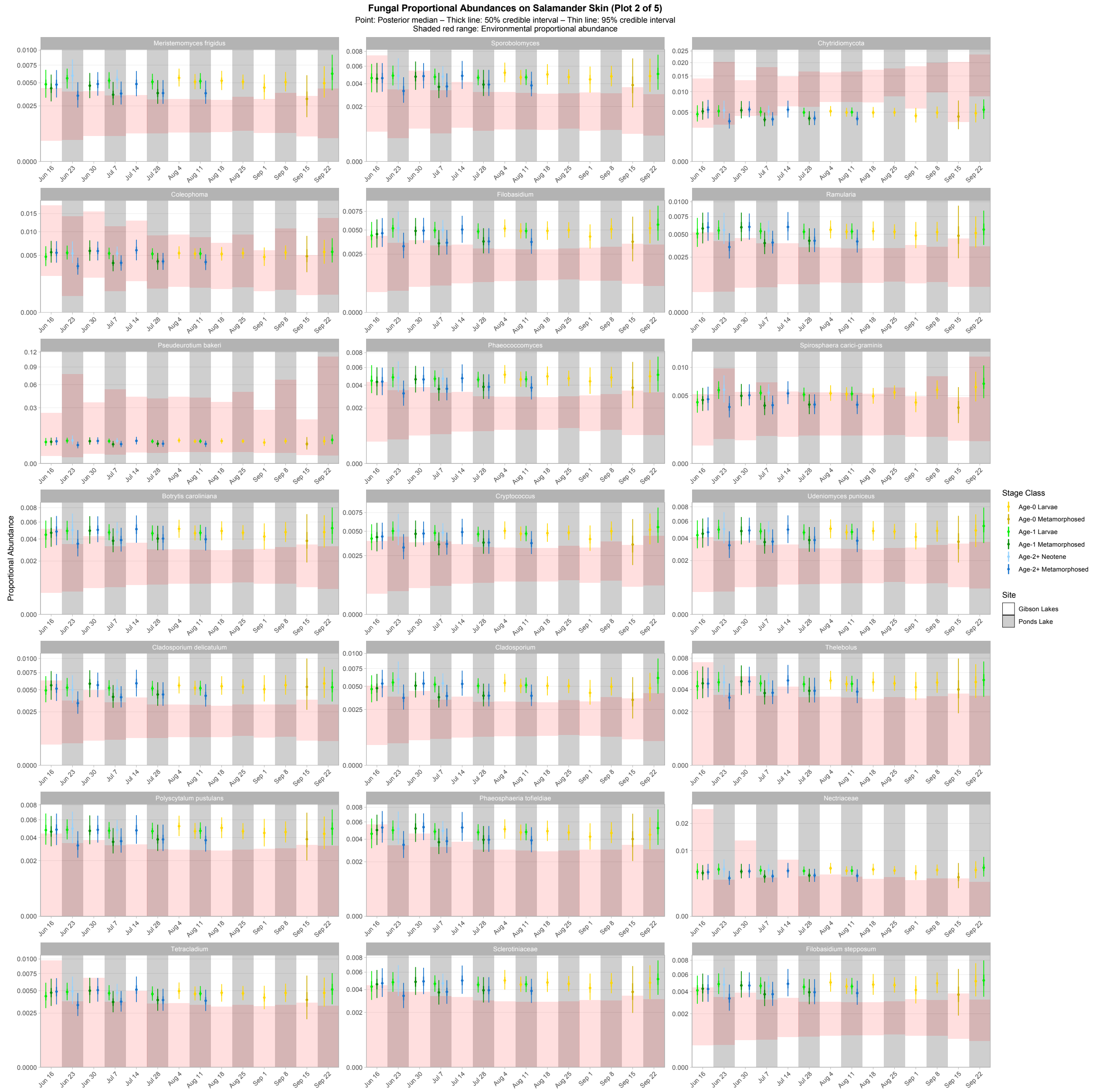

Figure S18. Second plot of proportional abundance predictions for the top 100 fungal taxa. Points, thick lines, and thin lines represent posterior medians, 50% credible intervals, and 95% credible intervals, respectively. Shaded red ranges represent environmental proportional abundances. Taxa without shaded red ranges were not detected in either water or substrate. Note the square root scale on the y-axis. Taxa are ordered (left to right, top to bottom) by descending average proportion of reads in salamander samples in which each combination of site and life stage receives equal weight.

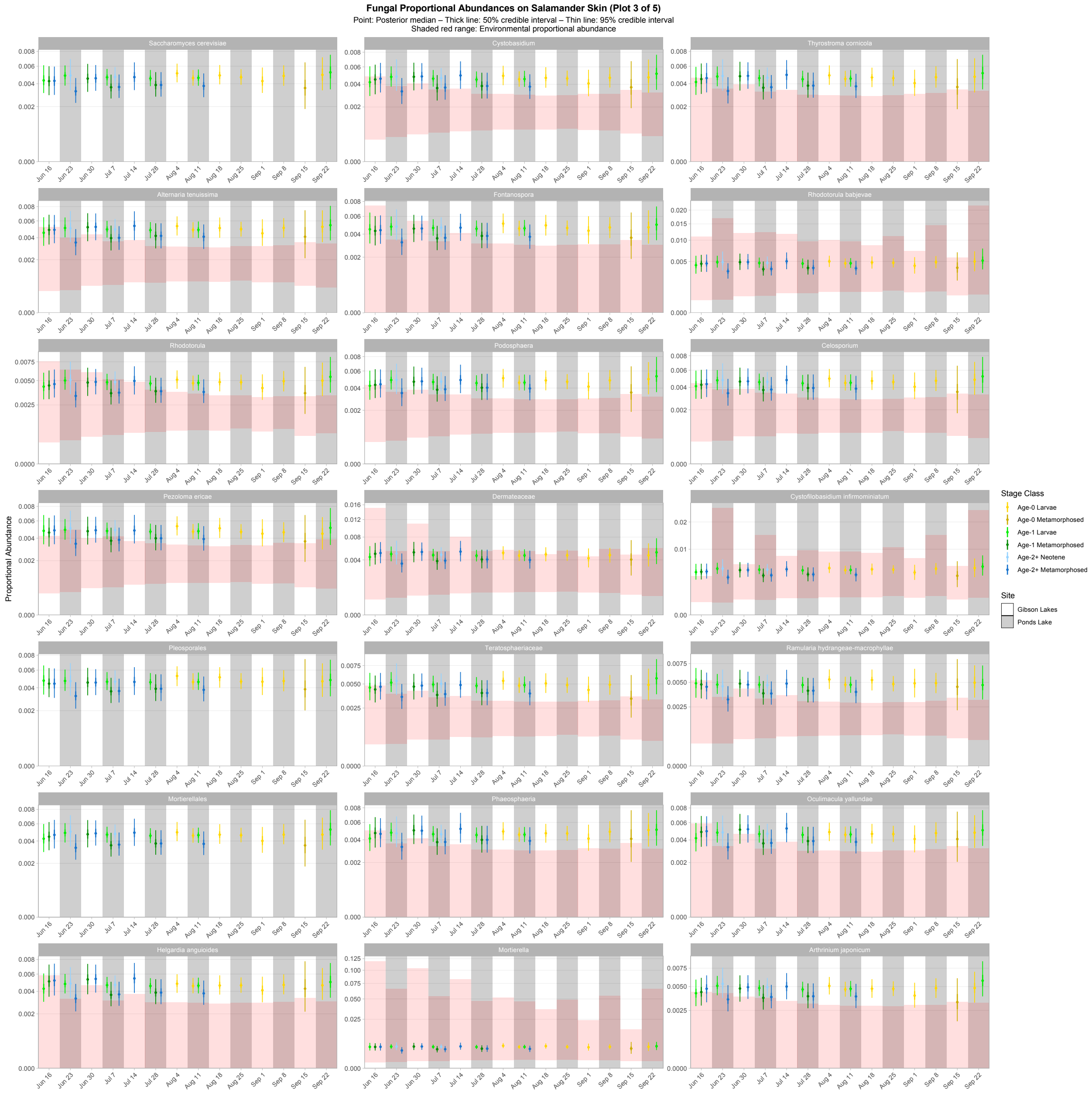

Figure S19. Third plot of proportional abundance predictions for the top 100 fungal taxa. Points, thick lines, and thin lines represent posterior medians, 50% credible intervals, and 95% credible intervals, respectively. Shaded red ranges represent environmental proportional abundances. Taxa without shaded red ranges were not detected in either water or substrate. Note the square root scale on the y-axis. Taxa are ordered (left to right, top to bottom) by descending average proportion of reads in salamander samples in which each combination of site and life stage receives equal weight.

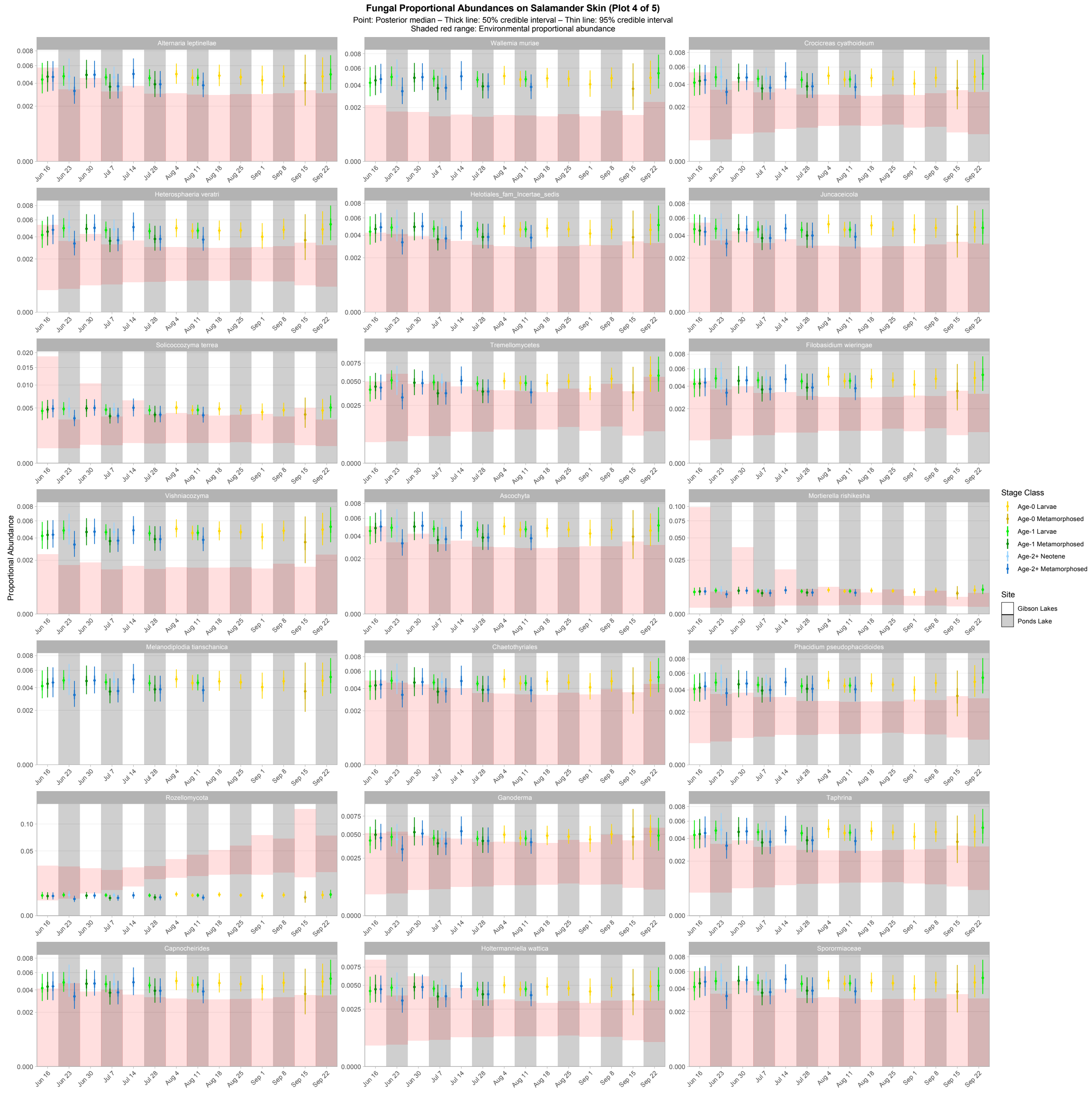

Figure S20. Fourth plot of proportional abundance predictions for the top 100 fungal taxa. Points, thick lines, and thin lines represent posterior medians, 50% credible intervals, and 95% credible intervals, respectively. Shaded red ranges represent environmental proportional abundances. Taxa without shaded red ranges were not detected in either water or substrate. Note the square root scale on the y-axis. Taxa are ordered (left to right, top to bottom) by descending average proportion of reads in salamander samples in which each combination of site and life stage receives equal weight.

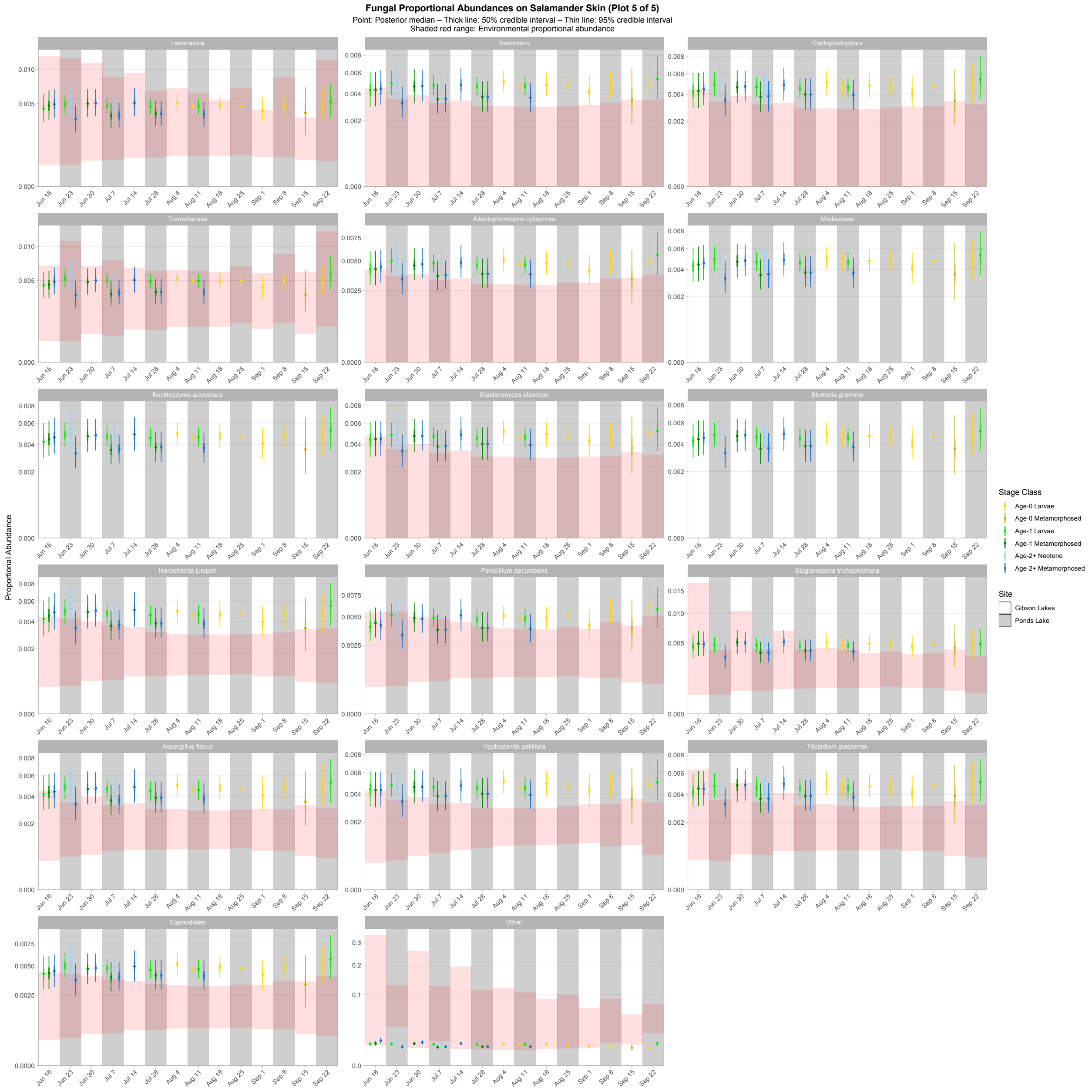

Figure S21. Fifth plot of proportional abundance predictions for the top 100 fungal taxa. Points, thick lines, and thin lines represent posterior medians, 50% credible intervals, and 95% credible intervals, respectively. Shaded red ranges represent environmental proportional abundances. Taxa without shaded red ranges were not detected in either water or substrate. Note the square root scale on the y-axis. Taxa are ordered (left to right, top to bottom) by descending average proportion of reads in salamander samples in which each combination of site and life stage receives equal weight.

#### *ITS Bray-Curtis dissimilarity*

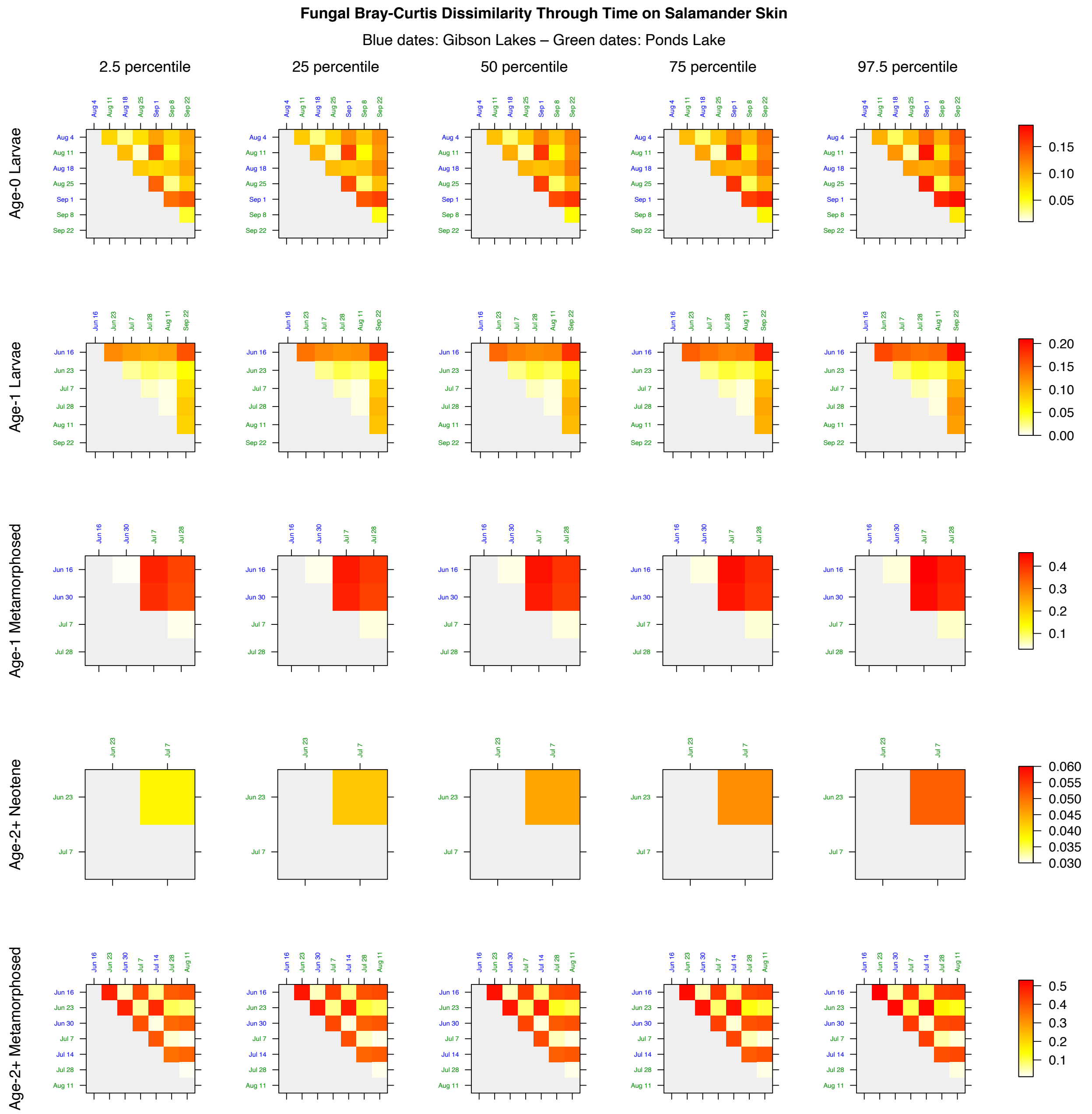

Figure S22. Bray-Curtis dissimilarity through time for fungal communities on salamander skin.

### *16S absolute abundance plots*

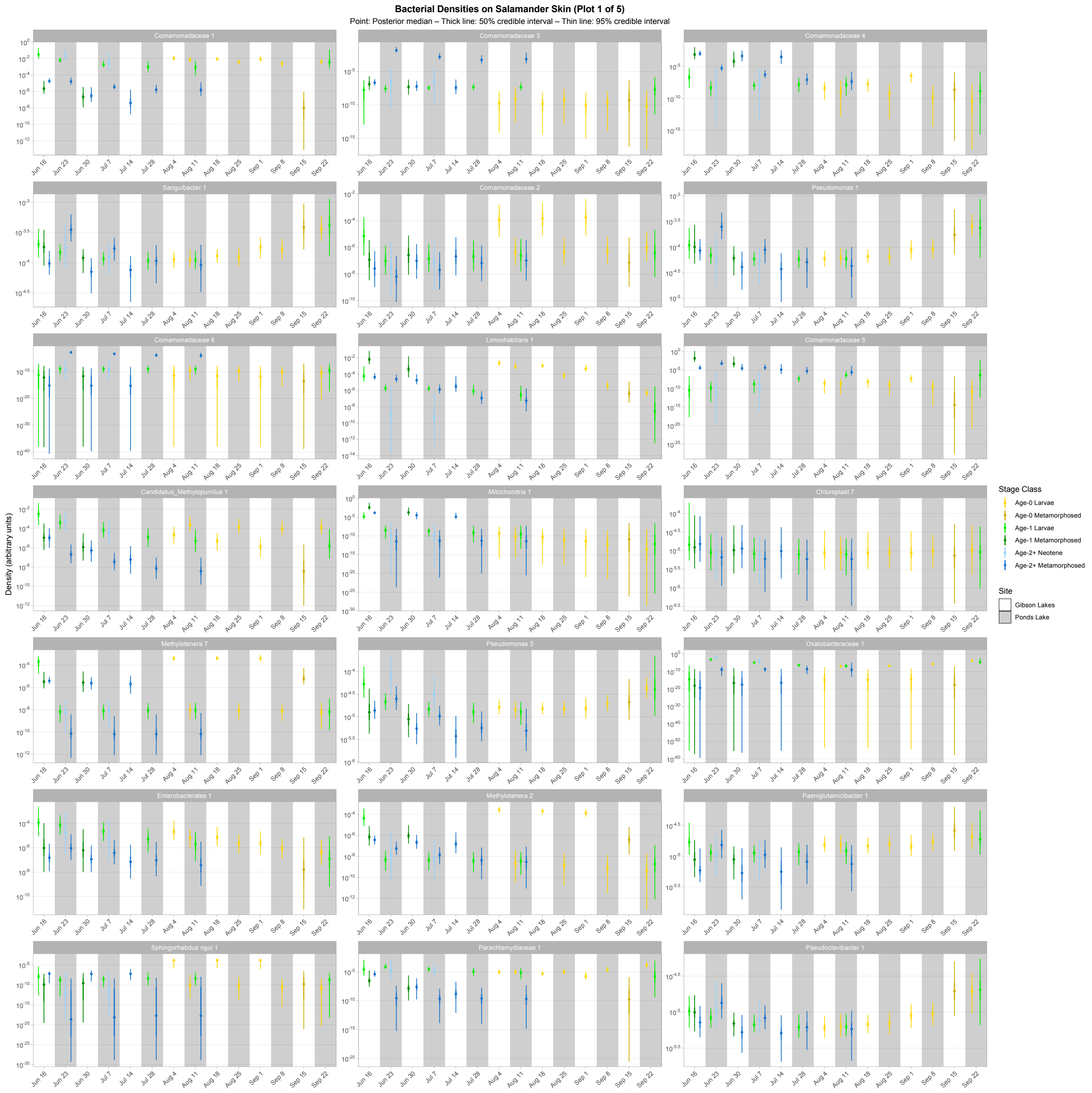

Figure S23. First plot of density predictions for the top 100 bacterial taxa on salamander skin. Points, thick lines, and thin lines represent posterior medians, 50% credible intervals, and 95% credible intervals, respectively. Note the $\log_{10}$ scale on the y-axis. Taxa are ordered (left to right, top to bottom) by descending average proportion of reads in salamander samples in which each combination of site and life stage receives equal weight.

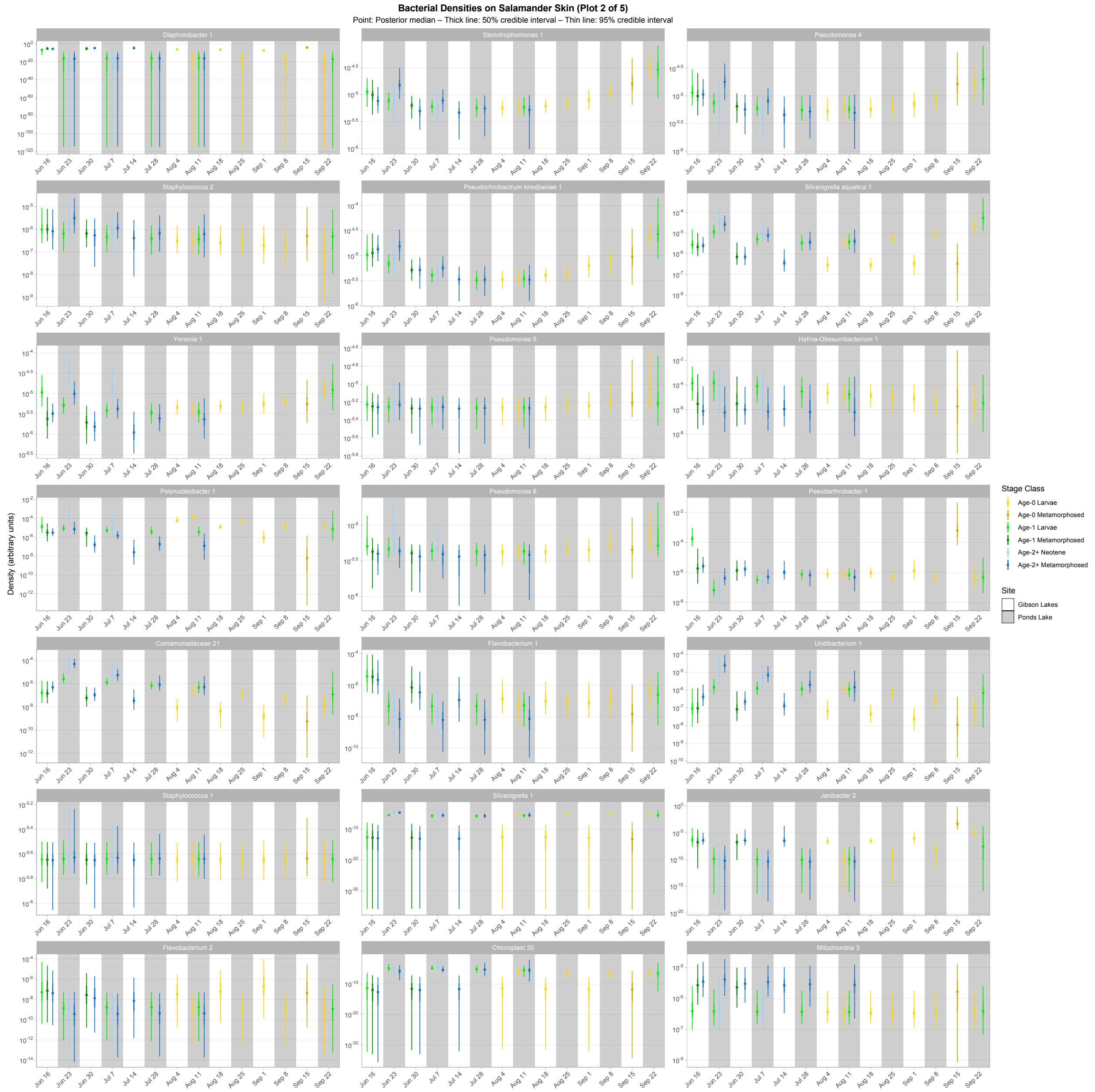

Figure S24. Second plot of density predictions for the top 100 bacterial taxa on salamander skin. Points, thick lines, and thin lines represent posterior medians, 50% credible intervals, and 95% credible intervals, respectively. Note the $\log_{10}$ scale on the y-axis. Taxa are ordered (left to right, top to bottom) by descending average proportion of reads in salamander samples in which each combination of site and life stage receives equal weight.

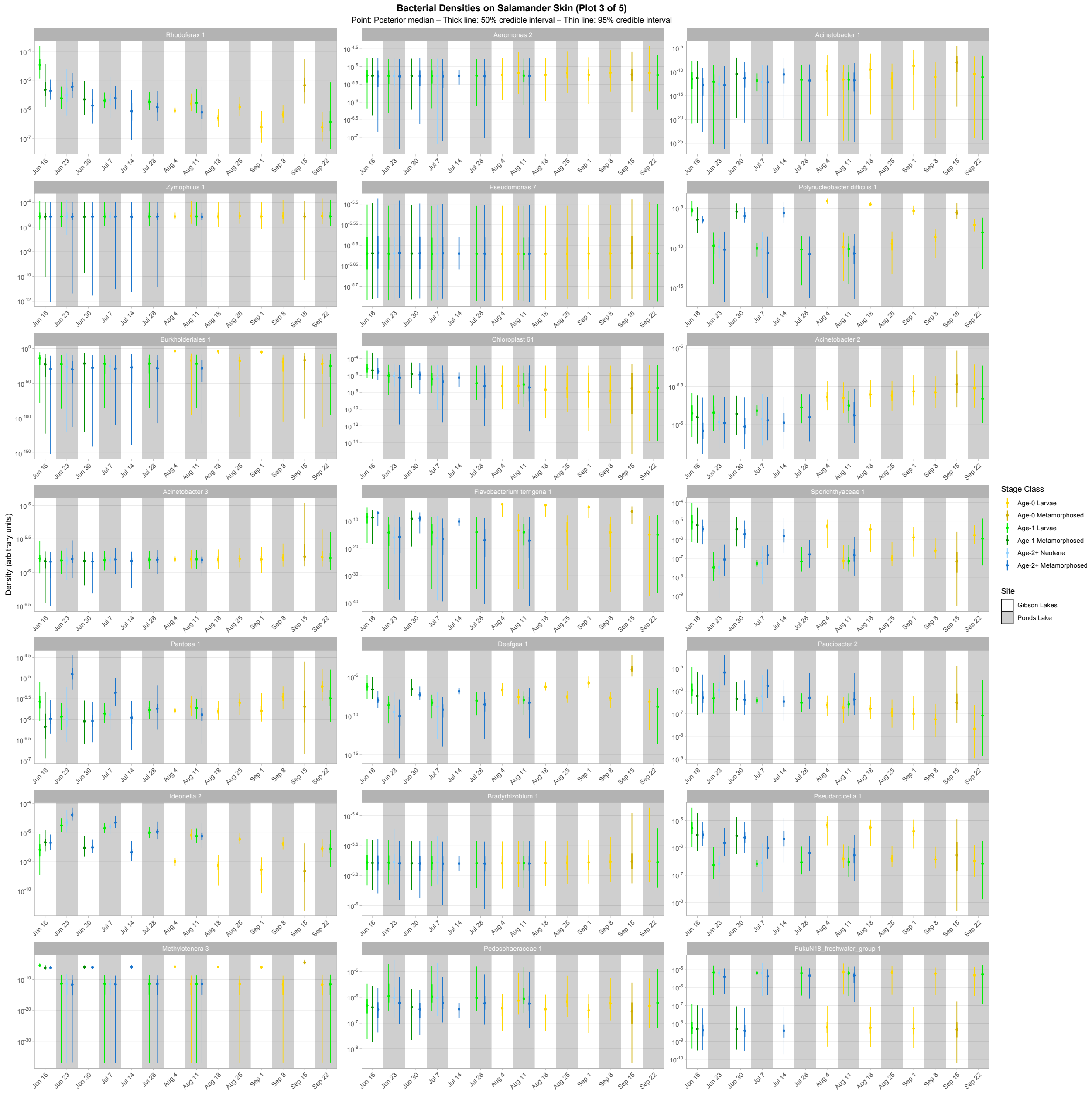

Figure S25. Third plot of density predictions for the top 100 bacterial taxa on salamander skin. Points, thick lines, and thin lines represent posterior medians, 50% credible intervals, and 95% credible intervals, respectively. Note the $\log_{10}$ scale on the y-axis. Taxa are ordered (left to right, top to bottom) by descending average proportion of reads in salamander samples in which each combination of site and life stage receives equal weight.

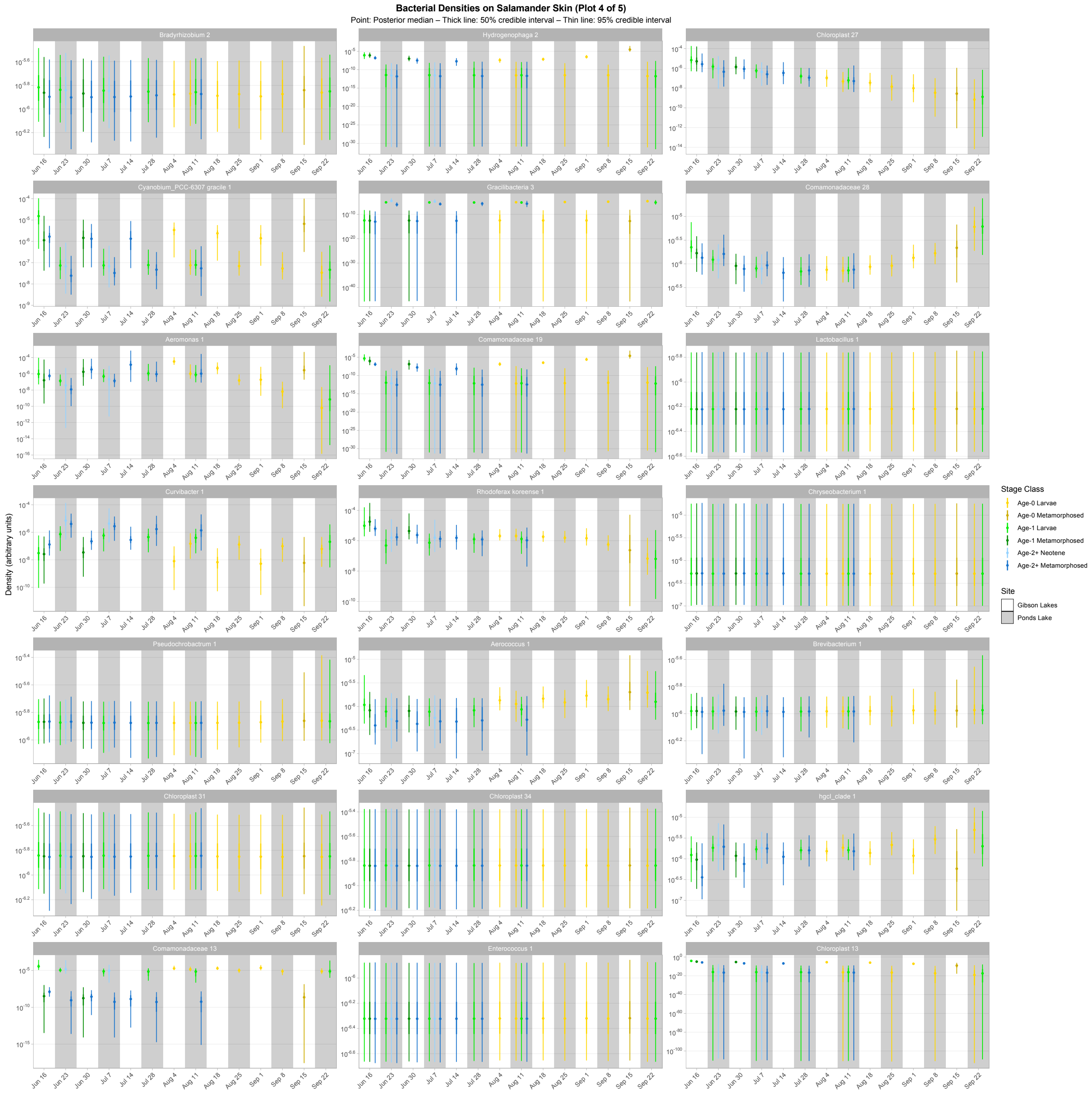

Figure S26. Fourth plot of density predictions for the top 100 bacterial taxa on salamander skin. Points, thick lines, and thin lines represent posterior medians, 50% credible intervals, and 95% credible intervals, respectively. Note the $\log_{10}$ scale on the y-axis. Taxa are ordered (left to right, top to bottom) by descending average proportion of reads in salamander samples in which each combination of site and life stage receives equal weight.

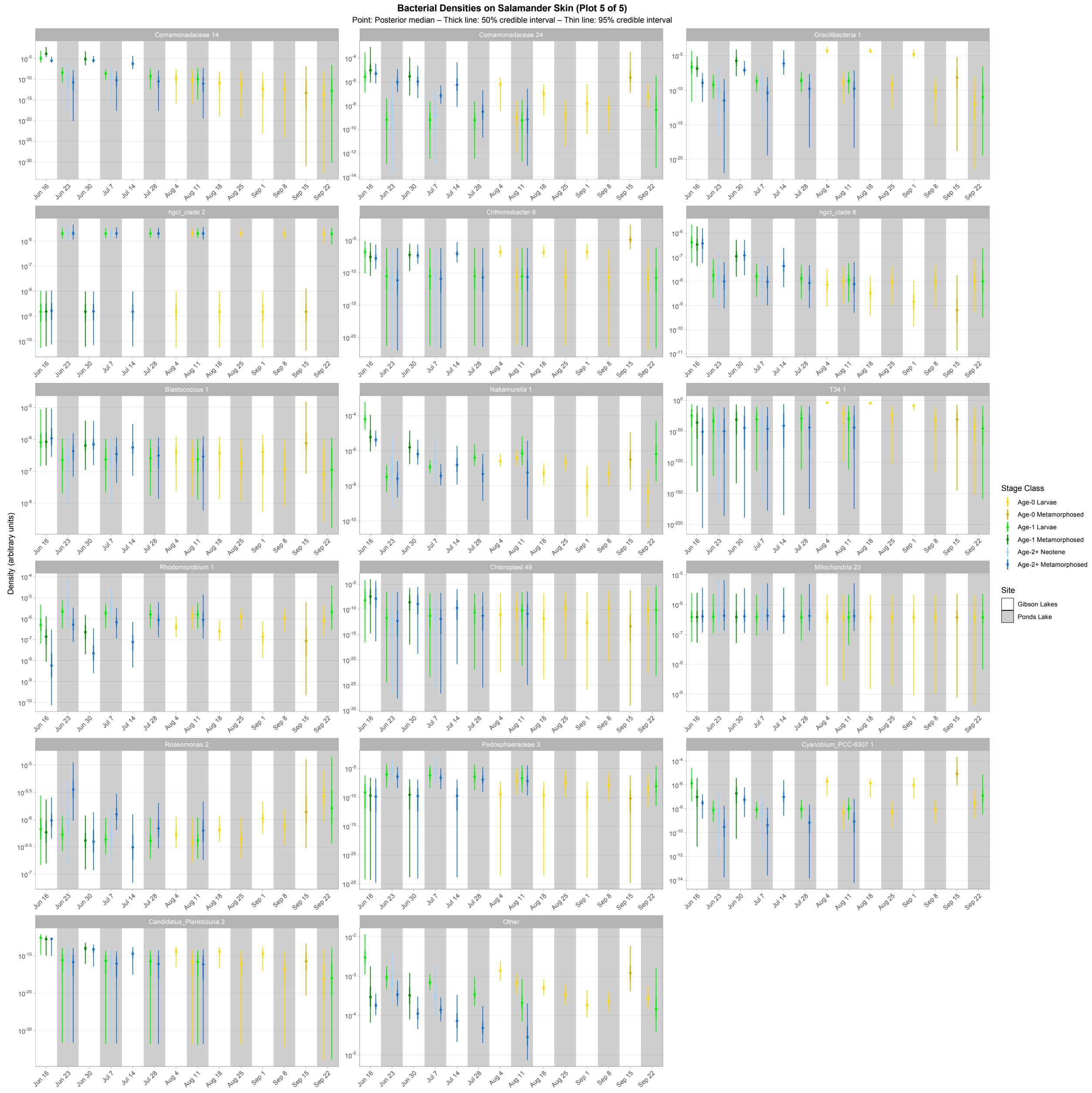

Figure S27. Fifth plot of density predictions for the top 100 bacterial taxa on salamander skin. Points, thick lines, and thin lines represent posterior medians, 50% credible intervals, and 95% credible intervals, respectively. Note the $\log_{10}$ scale on the y-axis. Taxa are ordered (left to right, top to bottom) by descending average proportion of reads in salamander samples in which each combination of site and life stage receives equal weight.

### *ITS absolute abundance plots*

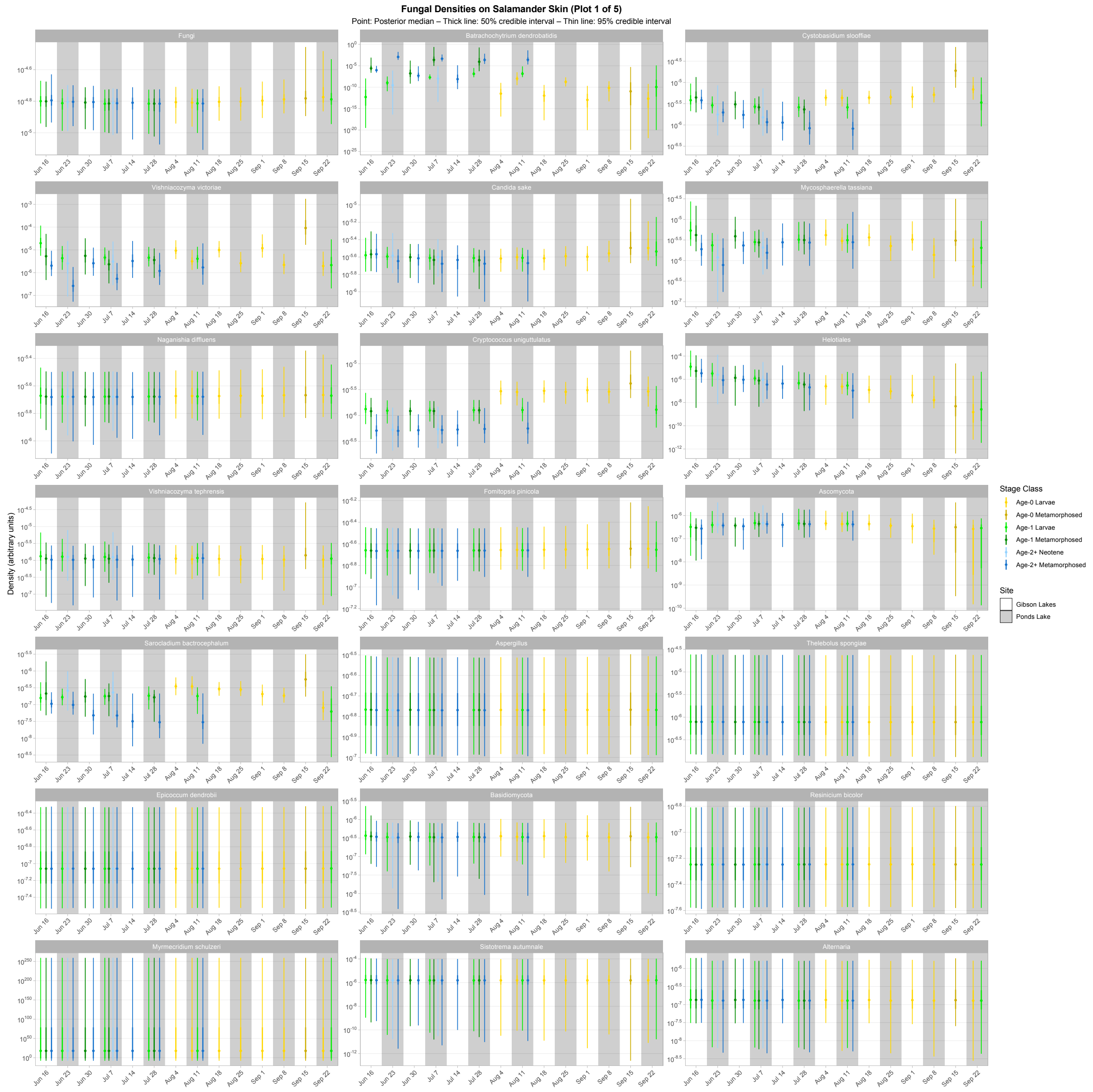

Figure S28. First plot of density predictions for the top 100 fungal taxa on salamander skin. Points, thick lines, and thin lines represent posterior medians, 50% credible intervals, and 95% credible intervals, respectively. Note the $\log_{10}$ scale on the y-axis. Taxa are ordered (left to right, top to bottom) by descending average proportion of reads in salamander samples in which each combination of site and life stage receives equal weight.

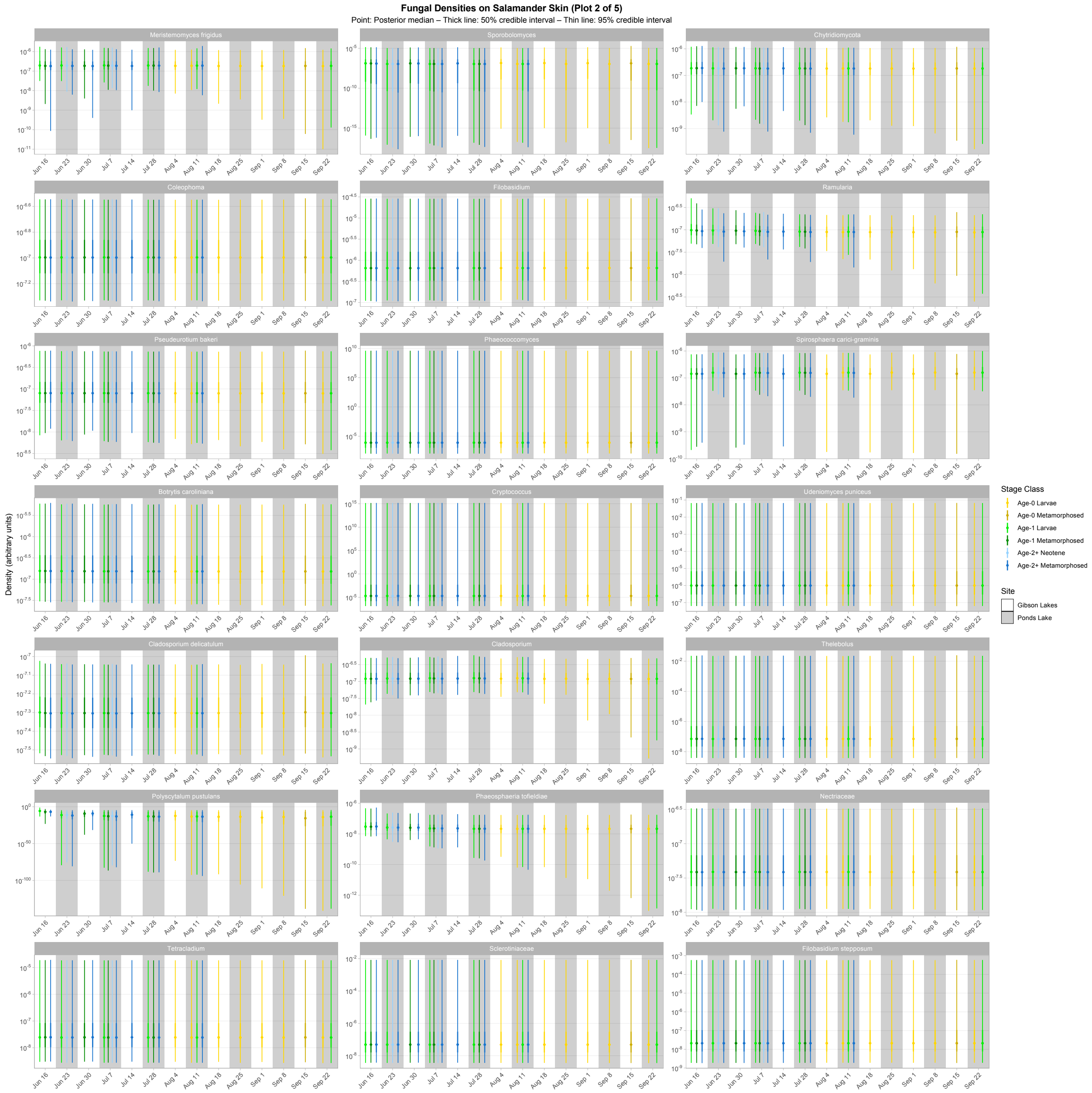

Figure S29. Second plot of density predictions for the top 100 fungal taxa on salamander skin. Points, thick lines, and thin lines represent posterior medians, 50% credible intervals, and 95% credible intervals, respectively. Note the $\log_{10}$ scale on the y-axis. Taxa are ordered (left to right, top to bottom) by descending average proportion of reads in salamander samples in which each combination of site and life stage receives equal weight.

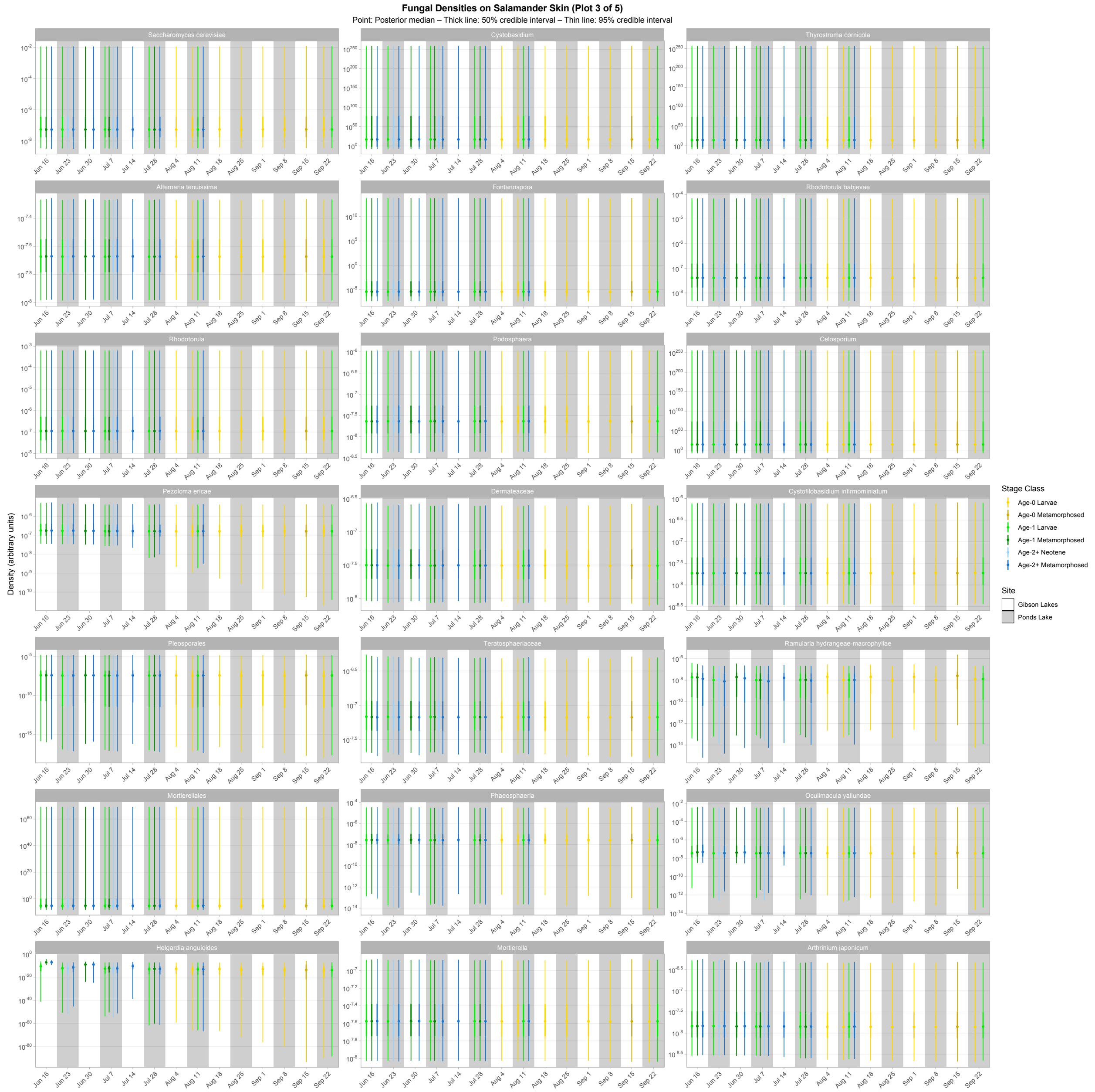

Figure S30. Third plot of density predictions for the top 100 fungal taxa on salamander skin. Points, thick lines, and thin lines represent posterior medians, 50% credible intervals, and 95% credible intervals, respectively. Note the $\log_{10}$ scale on the y-axis. Taxa are ordered (left to right, top to bottom) by descending average proportion of reads in salamander samples in which each combination of site and life stage receives equal weight.

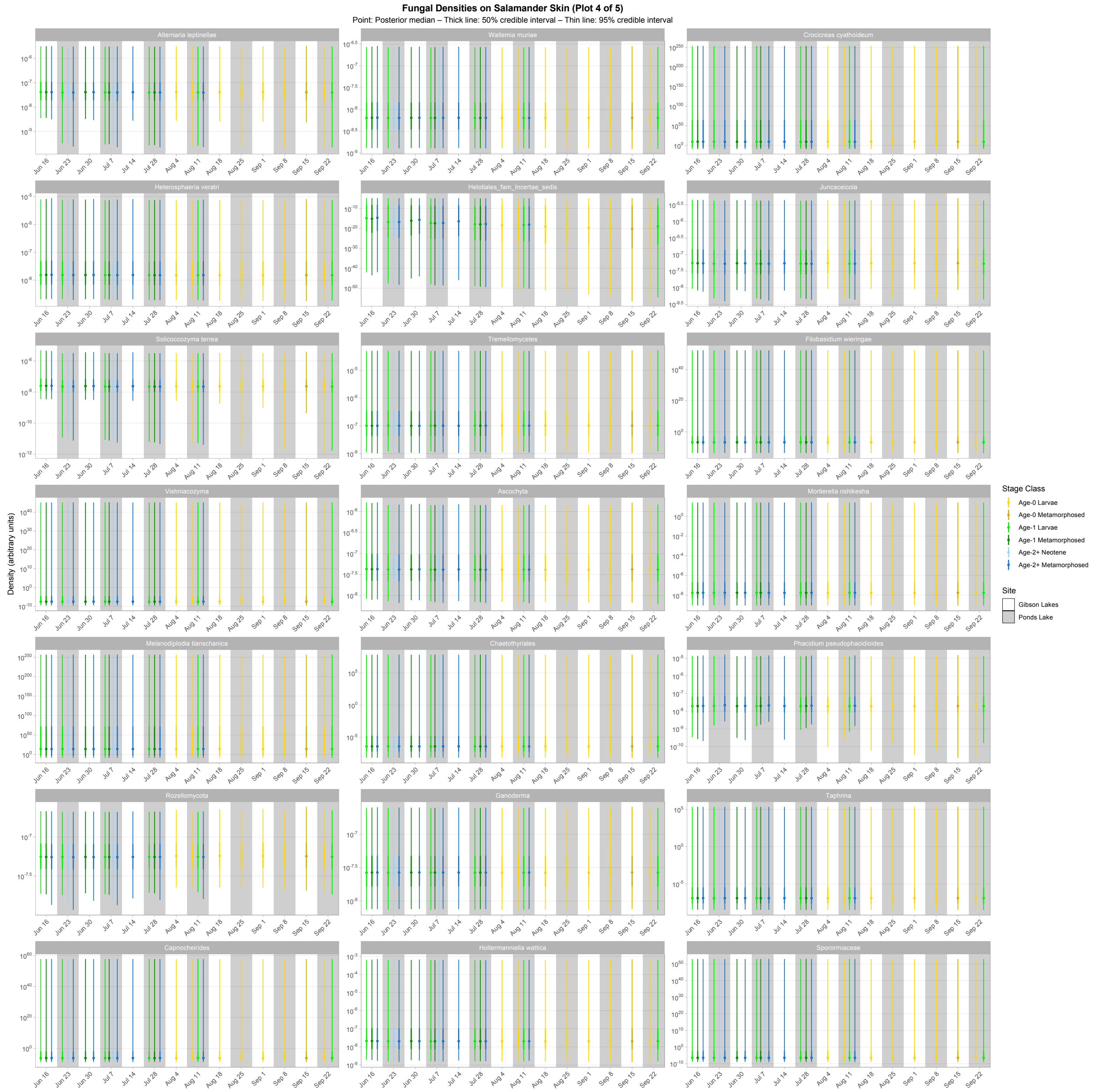

Figure S31. Fourth plot of density predictions for the top 100 fungal taxa on salamander skin. Points, thick lines, and thin lines represent posterior medians, 50% credible intervals, and 95% credible intervals, respectively. Note the $\log_{10}$ scale on the y-axis. Taxa are ordered (left to right, top to bottom) by descending average proportion of reads in salamander samples in which each combination of site and life stage receives equal weight.

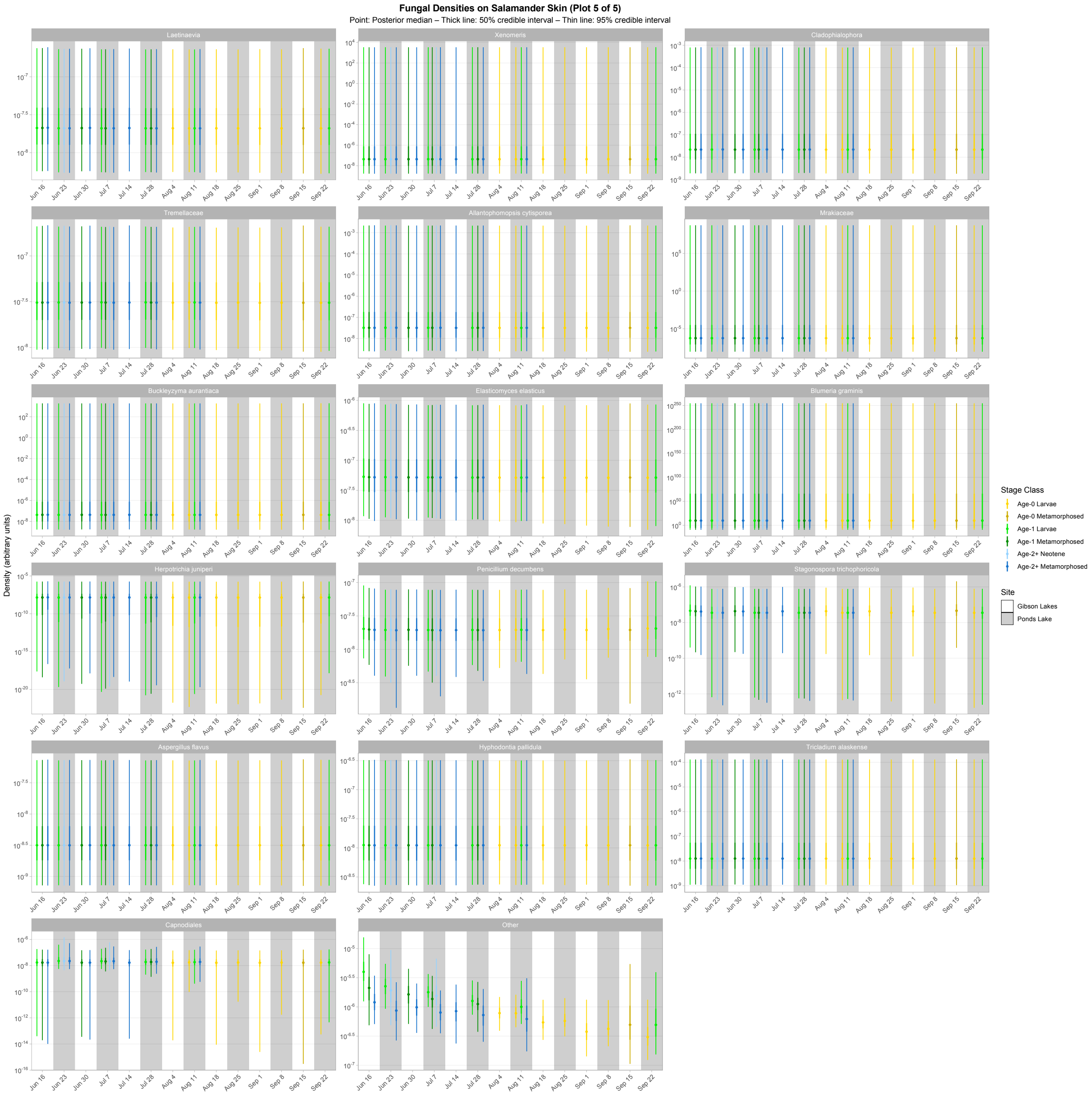

Figure S32. Fifth plot of density predictions for the top 100 fungal taxa on salamander skin. Points, thick lines, and thin lines represent posterior medians, 50% credible intervals, and 95% credible intervals, respectively. Note the $\log_{10}$ scale on the y-axis. Taxa are ordered (left to right, top to bottom) by descending average proportion of reads in salamander samples in which each combination of site and life stage receives equal weight.

#### *Absolute abundance of Bd-inhibition categories*

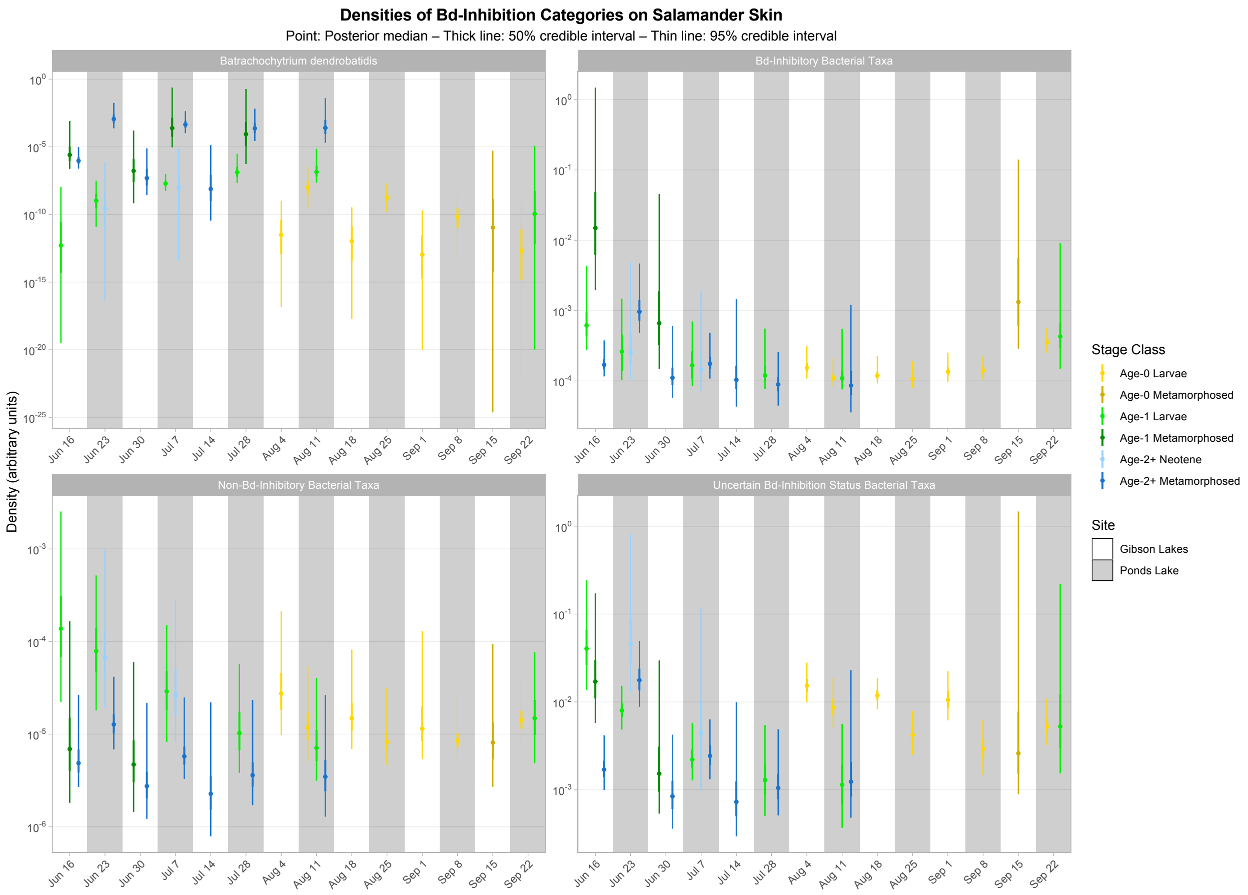

Figure S33. Density predictions of *Batrachochytrium dendrobatidis* (*Bd*) and *Bd*-inhibitory bacterial taxa from the top 100 on salamander skin. Points, thick lines, and thin lines represent posterior medians, 50% credible intervals, and 95% credible intervals, respectively. Note the $\log_{10}$ scale on the y-axis.

### Model code

#### *Stan code for Bayesian Dirichlet-multinomial regression model*

```
//// Dirichlet-Multinomial Regression Model.

data{

  //// Define variables.
  int<lower=1> NSamples; // Number of samples.
  int<lower=1> NTaxa; // Number of taxa.
  int<lower=1> NStrata; // Number of strata.
  matrix[NSamples,NStrata] StratumMatrix; // Predictor matrix for strata.
  int<lower=1> NPredictors; // Number of non-strata predictors.
```

```
  matrix[NSamples,NPredictors] PredictorMatrix; // Non-strata predictor matrix.
  int ReadsMatrix[NSamples,NTaxa]; // Reads response matrix.
  real<lower=0> sd_prior; // Regression coefficient and precision parameter standard deviation
        prior.

}

parameters{

  //// Specify parameters.
  simplex[NTaxa] p[NSamples]; // Each sample's set of modeled proportions.
  vector[NTaxa-1] beta_0_raw; // Intercept raw regression coefficient vector.
  matrix[NTaxa-1,NStrata-1] beta_stratum_raw; // Stratum raw regression coefficient matrix.
  vector<lower=0>[NTaxa-1] sigma2_stratum; // Stratum common variance vector.
  matrix[NTaxa-1,NPredictors] beta_pred_raw; // Non-stratum raw regression coefficient matrix.
  real theta; // Precision parameter.

}

transformed parameters{

  //// Specify transformed parameters.
  vector[NTaxa] beta_0; // Intercept regression coefficient vector.
  matrix[NTaxa,NStrata] beta_stratum; // Stratum regression coefficient matrix.
  vector<lower=0>[NTaxa-1] sigma_stratum; // Stratum common standard deviation vector.
  matrix[NTaxa,NPredictors] beta_pred; // Non-stratum regression coefficient matrix.
  real exptheta; // Exponentiated precision parameter.

  //// Define transformed parameters.

  // Intercept regression coefficients.
  beta_0[NTaxa]=0; // Set last taxon's intercept to zero.
  for(k in 1:(NTaxa-1)){
    beta_0[k]=beta_0_raw[k]; // Set intercepts of other taxa to the raw betas.
  }

  // Stratum regression coefficients.
  for(j in 1:NStrata){
    beta_stratum[NTaxa,j]=0; // Set last taxon's stratum regression coefficients to zero.
  }
  for(k in 1:(NTaxa-1)){
    for(j in 1:(NStrata-1)){
      beta_stratum[k,j]=beta_stratum_raw[k,j]; // Set stratum regression coefficients of other taxa
        to the raw betas (execpt for the last stratum).
    }
```

```
  beta_stratum[k,NStrata]=-sum(beta_stratum_raw[k,1:(NStrata-1)]); // Apply a sum to zero
        constraint for the last stratum.
  sigma_stratum[k]=sqrt(sigma2_stratum[k]); // Calculate stratum common standard deviation as
        the square root of the common variance.
 }

 // Non-stratum regression coefficients.
 for(j in 1:NPredictors){
  beta_pred[NTaxa,j]=0; // Set last taxon's regression coefficients to zero.
  for(k in 1:(NTaxa-1)){
   beta_pred[k,j]=beta_pred_raw[k,j]; // Set regression coefficients of other taxa to the raw
        betas.
  }
 }

 // Create exponentiated precision parameter.
 exptheta=exp(theta);

}

model{

 //// Provide priors.

 // Intercept regression coefficients.
 for(k in 1:(NTaxa-1)){
  beta_0_raw[k]~normal(0,sd_prior);
 }

 // Stratum regression coefficients.
 for(k in 1:(NTaxa-1)){
  for(j in 1:(NStrata-1)){
   beta_stratum_raw[k,j]~normal(0,sigma_stratum[k]);
  }
 }

 // Stratum common variance.
 for(k in 1:(NTaxa-1)){
  sigma2_stratum[k]~inv_gamma(0.01,0.01);
 }

 // Non-stratum regression coefficients.
 for(j in 1:NPredictors){
  for(k in 1:(NTaxa-1)){
   beta_pred_raw[k,j]~normal(0,sd_prior);
```

```
    }
  }

  // Precision parameter.
  theta~normal(0,sd_prior);

  //// State likelihood.

  // Loop through each sample.
  for(i in 1:NSamples){

    // Define local variable.
    vector[NTaxa] eta;

    // Loop through each taxon.
    for(j in 1:NTaxa){
      // Linear model for eta.

          eta[j]=beta_0[j]+beta_stratum[j,1:NStrata]*transpose(StratumMatrix[i,1:NStrata])+beta_p
          red[j,1:NPredictors]*transpose(PredictorMatrix[i,1:NPredictors]);
    }

    // Relate sample eta's to probabilities with the Dirichlet distribution.
    p[i]~dirichlet(softmax(eta[1:NTaxa])*exptheta);

    // Relate probabilities to counts with the multinomial distribution.
    ReadsMatrix[i,1:NTaxa]~multinomial(p[i]);

  }

}
```

##### ***JAGS code for Bayesian negative binomial LASSO model***

```
model{

  ##################
  ### Likelihood ###
  ##################

  # Loop through each sample.
  for(i in 1:NSamples){

    # Linear model for density (arbitrary units) with a log link.
    log(density[i])<-beta_0+
```

```
                inprod(beta_stratum[1:NStrata],StratumMatrix[i,])+
                inprod(beta_pred[1:NPredictors],PredictorMatrix[i,])

  # The taxon expected count (mu) is the density multiplied by
  # synthgene count and a value proportional to swabbed area.
  mu[i]<-density[i]*SynthgeneCount[i]*Area[i]

  # Derive the negative binomial distribution's probability parameter (p).
  p[i]<-r/(r+mu[i])

  # Relate p and r to taxon counts with the negative binomial distribution.
  TaxonCount[i]~dnegbin(p[i],r)

}

##############
### Priors ###
##############

# Intercept term.
beta_0~dnorm(0,1e-6)

# Stratum regression coefficients.
## Loop through all but last stratum regression coefficient.
for(j in 1:(NStrata-1)){
  ## Normal scale mixture for pre-selection coefficients.
  beta_stratum_pre_sel[j]~dnorm(0,inverse_tau_squared_stratum)
}
## Relate binary inclusion variable to the inclusion probability.
IV_stratum~dbern(IP)
## Apply binary inclusion variable to perform variable selection.
beta_stratum[1:(NStrata-1)]<-IV_stratum*beta_stratum_pre_sel[1:(NStrata-1)]
## Sum to zero constraint on the last stratum regression coefficient.
beta_stratum[NStrata]<- -1*sum(beta_stratum[1:(NStrata-1)])

# Non-stratum regression coefficients.
for(j in 1:NPredictors){
  ## Normal scale mixture for pre-selection coefficients.
  beta_pred_pre_sel[j]~dnorm(0,inverse_tau_squared_pred[j])
  ## Relate binary inclusion variable to the inclusion probability.
  IV_pred[j]~dbern(IP)
  ## Apply binary inclusion variable to perform variable selection.
  beta_pred[j]<-IV_pred[j]*beta_pred_pre_sel[j]
}
```

```
  # Size parameter.
  r~dgamma(0.01,0.01)

  # Gamma for tau^2.
  ## Stratum.
  inverse_tau_squared_stratum<-1/tau_squared_stratum
  tau_squared_stratum~dgamma(NStrata/2,lambda_squared/2)
  ## Non-stratum.
  for(j in 1:NPredictors){
    inverse_tau_squared_pred[j]<-1/tau_squared_pred[j]
    tau_squared_pred[j]~dexp(lambda_squared/2)
  }

  # Gamma for lambda^2.
  lambda_squared~dgamma(0.01,0.01)

  # Inclusion probability.
  # (Prior expectation is to select 1% of the predictors.)
  IP~dbeta(0.02,1.98)

}
```

### References

All citations within the Supplemental Information are included in the references section of the main text.
